# Fold recognition by scoring protein map similarities using the congruence coefficient

**DOI:** 10.1101/2020.05.20.106484

**Authors:** Pietro Di Lena, Pierre Baldi

## Abstract

**Motivation:** Protein fold recognition is a key step for template-based modeling approaches to protein structure prediction. Although closely related folds can be easily identified by sequence homology search in sequence databases, fold recognition is notoriously more difficult when it involves the identification of distantly related homologues. Recent progress in residue-residue contact and distance prediction opens up the possibility of improving fold recognition by using structural information contained in predicted distance and contact maps.

**Results:** Here we propose to use the *congruence coefficient* as a metric of similarity between maps. We prove that this metric has several interesting mathematical properties which allow one to compute in polynomial time its exact mean and variance over all possible (exponentially many) alignments between two symmetric matrices, and assess the statistical significance of similarity between aligned maps. We perform fold recognition tests by recovering predicted target contact/distance maps from the two most recent CASP editions and over 27,000 non-homologous structural templates from the ECOD database. On this large benchmark, we compare fold recognition performances of different alignment tools with their own similarity scores against those obtained using the congruence coefficient. We show that the congruence coefficient overall improves fold recognition over other methods, proving its effectiveness as a general similarity metric for protein map comparison.

**Availability:** The software CCpro is available as part of the Scratch suite http://scratch.proteomics.ics.uci.edu/

## 1 Introduction

Computational approaches for protein structure prediction generally follow one of two broad strategies Kuhlman and Bradley (2019); Kryshtafovych *et al.* (2019): *template-free* (or *ab-initio*) modeling and *template-based modeling*, which uses known protein structures as templates for the structural modeling of the unknown protein structure. While closely related templates can easily be detected by using protein sequence search methods, the detection of distantly related templates needs more sophisticated fold recognition strategies. Popular approaches make use of sequence profiles, predicted secondary structure and solvent accessibility, and exploit diverse computational methods, such as linear programming, dynamic programming, hidden Markov models, as well as other machine learning methods Jones and Thompson (1993); Lemer *et al.* (1995). However, despite considerable progress, remote homology detection remains a challenging problem.

The most recent Critical Assessment of Structure Prediction experiment (CASP13) held in 2018 reported a dramatic improvement in protein structure prediction for both template-free and template-based modeling Kryshtafovych *et al.* (2019). This improvement has been driven primarily by the successful applications of *deep-learning* approaches Di Lena *et al.* (2012); Kandathil *et al.* (2019) and *direct coupling analysis* De Juan *et al.* (2013) to predict intra-residues distances and contacts Hou *et al.* (2019); Zheng *et al.* (2019); Shrestha *et al.* (2019); Senior *et al.* (2019); Xu and Wang (2019).

The recent progress in intra-residue distance and contact prediction opens up the possibility to further improve fold recognition by database searches using predicted distance/contact maps. This requires addressing two distinct problems: i) developing efficient two-dimensional alignment procedures for map comparison; ii) developing a good scoring function to measure the fitness between target maps and templates.

In this work we deal explicitly with the second problem, and partially with the first problem, by exploiting the *congruence coefficient* Burt (1948) as a measure of similarity between aligned maps. The congruence coefficient bears some similarity to the Pearson’s correlation coefficient, since its value lies in the [−1, 1] interval and it is insensitive to multiplication (but not addition) by a constant factor. However, unlike Pearson’s correlation coefficient, its normalization factor is invariant with respect to any alignment between two maps, which makes it particularly suitable as an objective function for alignment procedures.

We prove some interesting statistical properties of the congruence coefficient, such as a measure of statistical significance and polynomial-time formulas for computing both the exact mean and variance of the coefficient over all possible (exponential number of) alignments between two symmetric matrices. Such statistical properties are complementary. The statistical significance of the congruence coefficient can be used to detect statistically significant similarities between two aligned maps, which improves template ranking for predicted target maps. Conversely, the mean and variance of the congruence coefficient over all possible alignments can be used to compute the alignment *Z*-scores, which give indications on the quality of the alignments.

We test the fold recognition performances of the congruence coefficient by recovering *predicted maps* from the last two CASP editions and over 27,000 *structural templates* from the ECOD database Cheng *et al.* (2014) that do no share sequence similarity with the CASP targets. In detail, for performance assessment with predicted contact maps we use residue-residue predictions at CASP12 and CASP13 and three contact map alignment software: AlEigen Di Lena *et al.* (2010), EigenThreader Buchan and Jones (2017) and Map_Align Ovchinnikov *et al.* (2017). All three tools return an alignment between two input maps together with a similarity score. Keeping fixed the alignments, we compute the congruence coefficient between target and structural templates. Performances have been then assessed by comparing fold recognition accuracy with the congruence coefficient versus the original similarity score. A statistical analysis of alignment quality is also provided in order to evaluate to which extend alignment quality affects fold recognition performances. Since there is no CASP category for predicted distance maps and there are no standalone tools for distance map alignment, we assess fold recognition accuracy by using regular protein structure predictions at CASP and structural alignment tools CE Shindyalov and Bourne (1998) and TM-align Zhang and Skolnick (2005). In this case, we recover predicted distance maps from predicted structures, use the structural alignments to induce alignments between distance maps and then compute the congruence coefficient between the aligned maps. Performance assessment is achieved by comparison of fold recognition accuracy with the congruence coefficient versus the specific structural alignment similarity scores. Also in this case, alignment *Z*-scores are used to asses alignment quality and its impact on fold recognition performances. Although fold recognition with distance maps recovered from structural predictions may appear artificial, it provides a fair evaluation of fold recognition by protein distance maps. Overall our tests provide a benchmark to compare the congruence coefficient to other structural alignment metrics, in both contact-based and distance-based fold recognition.

As a general conclusion, fold recognition with predicted contact maps is significantly improved by using the congruence coefficient score as a fitness function. In comparison to structural alignment metrics, the congruence coefficient shows comparable or better fold recognition accuracy, proving its potential as general similarity metric for protein map comparisons.

## 2 Materials and methods

### 2.1 Congruence coefficient

#### 2.1.1 Definition

The congruence coefficient was first introduced in Burt (1948), with the name of *unadjusted correlation*, as a measure of similarity in factor analysis.

##### Definition 2.1.

*Let X, Y ∈* ℝ^*m×n*^ *be two real matrices. The congruence coefficient between X, Y is defined by:*

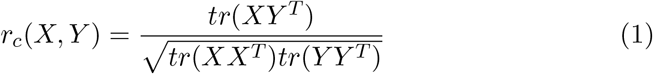

The *r*_*c*_ score is between −1 and +1, with *r*_*c*_ = 1 representing the highest degree of similarity. Although the congruence coefficient appears quite similar to the Pearson’s correlation coefficient, the latter measures the deviations from the mean whereas the congruence coefficient measures the deviations from zero. Like the correlation coefficient, the congruence coefficient is insensitive to the multiplication of the matrices *X, Y* by constant factors different from zero. Unlike the correlation coefficient, it is sensitive to the addition of constant factors.

Although the *r*_*c*_ coefficient can be computed for non-square matrices, here we focus on protein contact and distance maps, which are both represented by square (symmetric) matrices. Typically, contact and distance maps of different proteins have different sizes determined by the protein sequence lengths, thus the *r*_*c*_ coefficient between two contact/distance maps can be computed only if an *alignment* between the two matrices is provided. Aligned matrices can be simply obtained by introducing rows and (respective) columns of zeroes in the original symmetric matrices, which correspond to gaps in the alignments. Since zero (gap) rows/columns do not contribute in the trace of the products in Equation (1), an equivalent and simpler formulation of the congruence coefficient with respect to some alignment can be obtained by leaving unchanged the *X* matrix and by removing all rows/columns in the *Y* matrix that match a gap row/column in the aligned *X* matrix (Section 3, Suppl.) In this way, we can just recode the *Y* matrix as follows.

##### Definition 2.2.

*A partial function α* : {1, .., *m*} → {1, .., *n*} *is an alignment if* ∀*i* ≠ *j such that α*(*i*) ≠ ⊥ *and α*(*j*) ≠ ⊥ *then:*

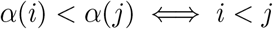

where *α*(*i*) = ⊥ means that *α* is not defined on *i*. Given a matrix *Y* ∈ ℝ^*n×n*^ and an alignment *α* : {1, .., *m*} → {1, .., *n*}, we define the new matrix *Y* ^*α*^ ∈ ℝ^*m×m*^ by:

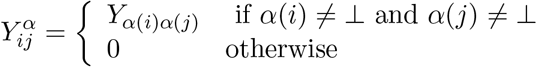

Now, let *X* ∈ ℝ^*m×m*^, *Y* ∈ ℝ^*n×n*^ be two symmetric matrices and *α* : {1, .., *m*} → {1, .., *n*} an alignment. We define (Section 3, Suppl.) the congruence coefficient with respect to the alignment *α* by

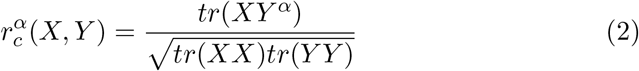

Note that, the normalization factor in Equation (2) is invariant with respect to any possible alignment *α*. Such property does not hold for Pearson’s correlation which measures the deviation from the mean value and is thus affected by the number of zero rows and columns introduced in the alignment. The alignment that maximizes the *r*_*c*_ coefficient in Equation (2) is thus simply the alignment that maximizes the trace of the product between the two aligned matrices.

#### 2.1.2 Statistical properties of the congruence coefficient

Here we show that the congruence coefficient has several desirable mathematical properties: its statistical significance can be rigorously assessed, and its mean and variance can be estimated in polynomial time. The details of our proofs are given in the Supplementary file.

Statistical hypothesis testing of the congruence coefficient between two aligned maps, under the null hypothesis that the coefficient is zero, can be reframed as a statistical hypothesis testing on the angle between two unitary vectors on some *N*-dimensional unit sphere, under the null hypothesis that the two vectors are orthogonal. The dimension *N* depends on the size and topology of the two input matrices. The p-value can be then computed as the ratio between the volume of the *N*-dimensional unit sphere and the volume of the hyper-spherical cap Li (2011) identified by the angle between the two unitary vectors. In summary, let *X* ∈ ℝ^*m×m*^ and *Y* ∈ ℝ^*n×n*^ be two symmetric matrices with zero main diagonal, and *α* : {1, .., *m*} → {1, .., *n*} an alignment. Then (Section 6 in Suppl.) the right-tailed p-value of the congruence coefficient 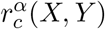 is given by:

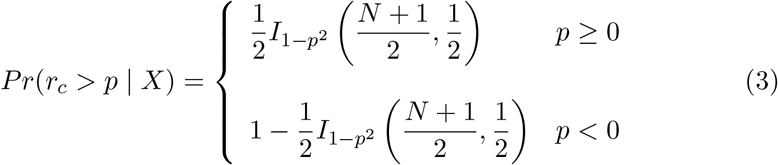

where *I* is the *regularized incomplete beta function*, 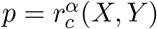, and the degree of freedom *N* is the number of non-zero elements in the upper (or lower) triangular portion of *X*. Given any symmetric matrix *X* ∈ ℝ^*m×m*^, Equation (3) gives the probability of uniformly sampling a random symmetric matrix *Y′* ∈ ℝ^*m×m*^ (with zero main diagonal), such that 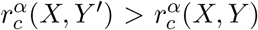. We can symmetrically use Equation (3) with known *Y*, where the degree of freedom *N* is the number of non-zero elements in the upper triangular portion of *Y*. The condition of having zero- main-diagonal is necessary, and trivially satisfied by distance maps, as well as contact maps (contacts between adjacent residues are typically ignored). In database searches, we use Equation (3) to asses whether two aligned matrices are significantly similar. That is, given a target matrix *X*, a template matrix *Y* and an alignment *α* between *X* and *Y*, we ignore template *Y* if 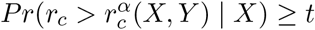 or 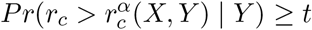, where *t* is the Bonferroni-corrected p-value cutoff 0.05.

The exact mean and variance of the *r*_*c*_ score under all possible alignments can be used to test the quality of a given alignment between two maps (i.e. Z-score). Given two symmetric matrices *X* ∈ ℝ^*m×m*^ and *Y* ∈ ℝ^*n×n*^, the expected value of the congruence coefficient between *X* and *Y* with respect to all possible alignments *α* is given by (Section 4.2 in Suppl.):

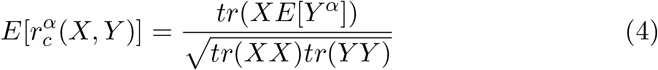

where *E*[*Y*^*α*^] ∈ ℝ^*m×m*^ is the *expectation matrix*, that averages all *Y*^*α*^ ∈ ℝ^*m×m*^ matrices. The expectation matrix can be computed from *Y* and *m*, without the need of *X*. Equivalently, the variance of the congruence coefficient between *X* and *Y* with respect to all possible alignments *α* is given by (Section 4.3 in Suppl.):

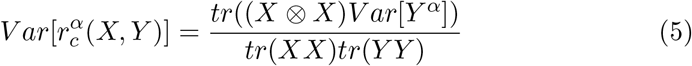

where ⊗ is the Kronecker product and:

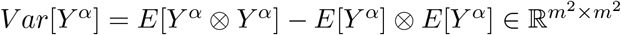

is the *variance-covariance matrix* of random matrices *Y*^*α*^ ∈ ℝ^*m×m*^. The variance-covariance matrix can also be computed using only *Y* and *m*. The computational time for the expectation matrix in Equation (4) is quadratic in the product of the lenghts *mn*, which is reasonably fast for native contact/distance maps. Instead, the computational time for the variance-covariance matrix in (5) is quartic in *mn*, which is challenging for large matrices. However, an ad-hoc sampling of alignments (i.e. proportional to the fraction of alignments of a given size) provides an almost exact estimation of the variance (Section 7.3, Suppl.).

In addition, following the combinatorial approach described in Kazi-Aouala *et al.* (1995), we can derive closed expressions for the expectation and variance of the congruence coefficient between two symmetric maps *X* and *Y* over all possible permutations of *Y*, and with respect to all possible alignments between *X* and *Y* (Section 5, Suppl.). We tried to exploit permutation statistics as a fast approach for approximating variance calculations over all possible alignments. However, tests on real protein contact/distance maps show that both expectation and variance over permutations poorly approximate expectation and variance over all possible alignments (Section 7.3, Suppl.). Thus the formula obtained do not seem to have an immediate application in protein map comparison, although they have intrinsic theoretical interest and may be useful in other contexts.

### 2.2 Template and Benchmark Data

Benchmark data sets were obtained from the CASP repository (Section 7.1 in Suppl.). For contact-based fold recognition assessment, we selected all residueresidue contact predictions submitted to the CASP12 and CASP13 experiments. For distance-based fold recognition assessment, we decided to *simulate* predicted distance maps by recovering them from the structural predictions at CASP12 and CASP13. This was necessary since distance map predictions were used as an intermediary step, rather than as a standalone problem, and such predictions were not available. We considered only the CASP targets for which the experimentally determined structure was available in the PDB and the fold annotation was available in the ECOD classification Cheng *et al.* (2014).

Template data were obtained from the ECOD database (Section 7.1 in Suppl.). ECOD protein domains are classified with respect to four groups: the *F-group* groups domains with significant sequence similarity; the *T-group* groups domains with similar topological connections; the *H-group* groups domain that are considered homologous based on different attributes (e.g. functional similarity, literature); and the *X-group* groups domains that are potentially homologous although there is not yet adeguate evidence to support their homology relationship. We downloaded the ECOD pre-filtered subset at 40% sequence identity. In order to prevent any sequence homology bias in our tests, we removed from the ECOD dataset all protein domains found by a hmmsearch Eddy (2011) scan of the CASP12 and CASP13 targets against the ECOD database. In order to identify a subset of hard targets, we matched the FM (Free Modeling) domains of the CASP targets with the domains identified in ECOD. Native contact and distance maps were extracted from the ECOD pdb domain files.

The exact number of ECOD templates used in our experiments, as well as the targets and FM targets in the two CASP benchmark datasets, are shown in Table 1.

**Table 1:**
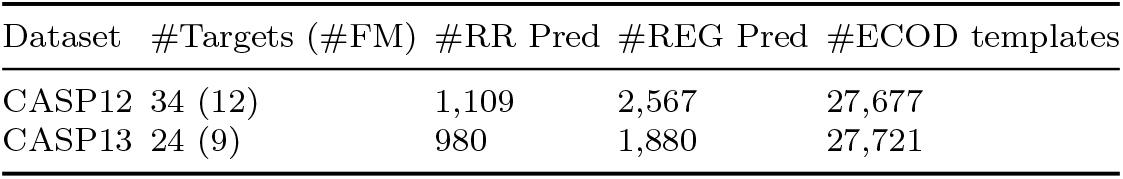
Benchmark set statistics. #Targets: number of CASP targets with fold annotation. #FM: number of targets containing FM domains. RR Pred: residue-residue contact predictions. REG Pred: regular structure predictions. #ECOD templates: number of sequence homology-free ECOD templates

### 2.3 Benchmark tools

#### Contact Maps

We considered three contact map alignment tools for performance comparison (Section 7.2 in Suppl.): AlEigen Di Lena *et al.* (2010), EigenThreader Buchan and Jones (2017), and Map_Align Ovchinnikov *et al.* (2017). We used the three tools to first align target and template maps and then to rank the templates: i) using the tool-specific scores; ii) using the congruence coefficient with respect to the alignments returned by the tools.

#### Distance Maps

Unlike contact map alignment, the standalone distance map alignment problem has received little or no attention in the literature. For this reason, for performance comparison, we decided to use two popular structural alignments tools (Section 7.2 in Suppl.), CE Shindyalov and Bourne (1998) and TM-Align Zhang and Skolnick (2005), and recover the distance map alignments from the structural alignments computed by both tools. Also in this case we rank the templates: i) using the tool-specific scores; ii) using the congruence coefficient with respect to the alignments returned by the tools.

Our choice of alignment tools has been driven mainly by speed considerations due to the very large number of comparison performed in our tests: a total of 58M contact map comparisons, and 123M structure comparisons, per method (see Table 1). The average running times of the benchmarked methods are summarized in Table 2.

**Table 2:**
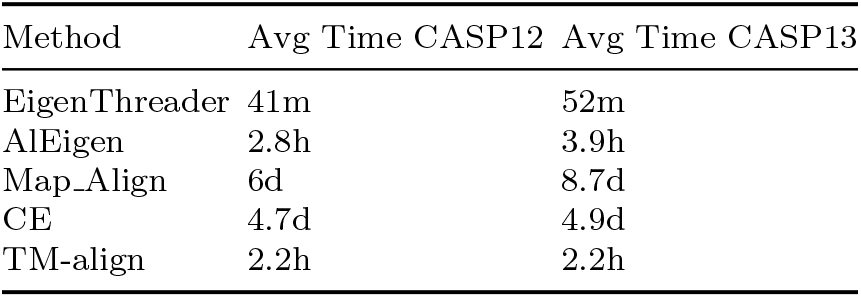
Average running time per prediction. m=minutes, h=hours, d=days

## 3 Results

### 3.1 Fold recognition with predicted contacts

For performance comparison, we search all residue-residue contact predictions submitted at CASP for a single target against the ECOD templates. This implies a maximum number of 38 predictions per target at CASP12 and 46 at CASP13, corresponding to the number of residue-residue prediction groups in the two CASP editions. True Positive Rate (TPR) fold recognition performances are assessed by selecting the top-1, top-5, top-10, and top-20 unique templates identified by the search with multiple predicted maps. For each top-*k* set, the TPR score is computed by counting the fraction of targets for which at least one template with similar fold is in the top-*k* hits. We assess the TPR performances separately for the four ECOD classes: *Family Level* (F), *Topology Level* (T), *Homology Level* (H), and *Possible Homology Level* (X). This implies that, for example, for TPR assessment at the Family Level we consider only the CASP targets that have been annotated at the Family Level in ECOD. The TPR performances on the CASP12 and CASP13 benchmark datasets, for the three map alignment tools AlEigen, EigenThreader, and Map_Align are summarized in Table 3. The table compares the performances of the three tools with their specific scoring schemes against those obtained using the congruence coefficient, indicated by AlEigen+*r*_*c*_, EigenThreader+*r*_*c*_, and Map_Align+*r*_*c*_, respectively.

**Table 3:**
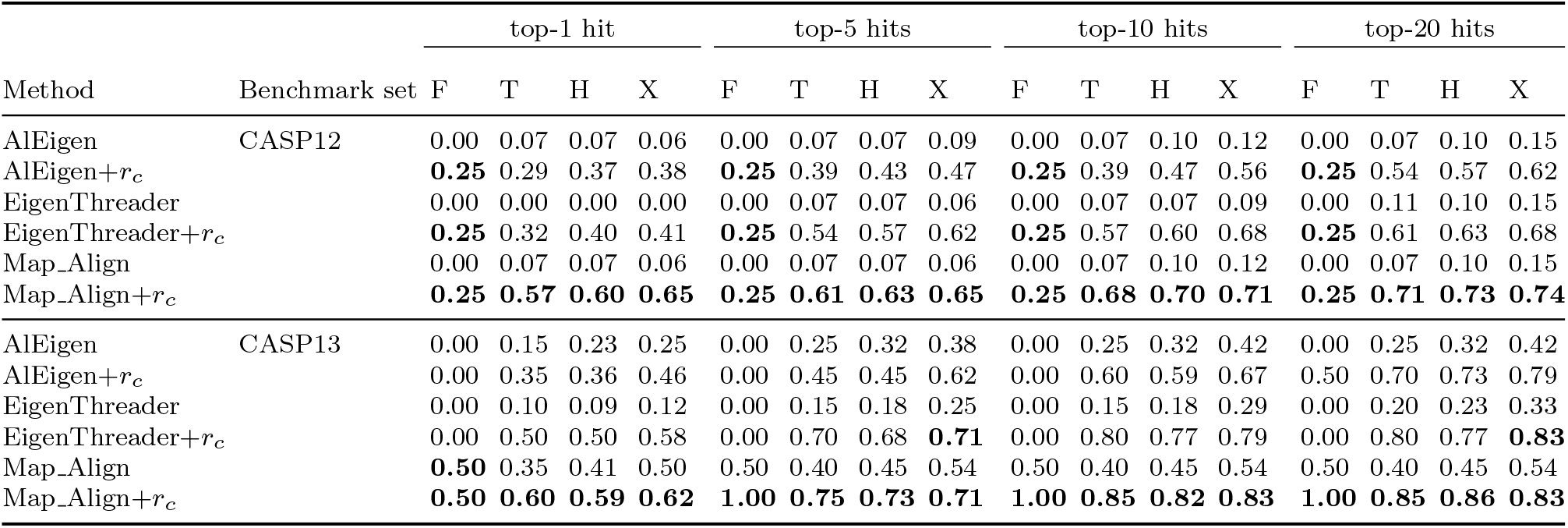
Fold recognition performances with predicted contacts. True Positive Rate (TPR) fold recognition performances on CASP12 and CASP13 benchmark sets. The TPR performances are assessed with respect to the **top-1**, **top-5**, **top10** and **top-20** ranked hits. ECOD hierarchy: (F) Family Level (4 targets in CASP12, 2 targets in CASP13), (T) Topology Level (28 targets in CASP12, 20 targets in CASP13), (H) Homology Level (30 targets in CASP12, 22 targets in CASP13), (X) Possible Homology Level (34 targets in CASP12, 24 targets in CASP13). EigenThreader, Map_Align and AlEigen use their own scoring system EigenThreader+*r*_*c*_, Map_Align+*r*_*c*_ and AlEigen+*r*_*c*_ use statistically significant congruence coefficient. Best TPR performances per column on CASP12 and CASP13 benchmark sets are highlighted in bold.

Fold recognition performances with predicted contact maps are influenced by three main factors: i) contact map prediction accuracy; ii) accurate alignments between target and templates; iii) proper scoring of the fitness between target and templates. The influence of a good scoring function is particularly evident for fold recognition performances in the CASP12 benchmark set (see Table 3), where the fold recognition precision is dramatically improved by the usage of the congruence coefficient for fitness ranking. We remark that for the computation of the *r*_*c*_ coefficient we use the alignments returned by the three packages, thus the low TPR performances of the three tools with their specific fitness functions are not an immediate consequence of poor contact map predictions or poor alignments. The improvement in fold recognition accuracy with *r*_*c*_ scoring can be observed also on the CASP13 benchmark set (see Table 3). In this case, the improvement is still significant, although less pronounced, since all the three methods show overall better performances with their own scoring functions in comparison to those achieved for CASP12. To a large extent, this can be imputed to better contact predictions for CASP13, which compensate for the lack of a good scoring function. In fact, if we restrict our tests to contact predictions submitted only by the top ranked predictors (using the official CASP rankings), we notice a general improvement in fold recognition accuracy for all methods (Section 7.4.1, Suppl.). Interestingly, the improvement is almost negligible or absent for *r*_*c*_ ranking performances, which indicates that the congruence coefficient can *filter out most of the noisy similarities*. This is partially a consequence of the statistically significant p-value cutoff applied to the rankings (Section 7.4.3, Suppl.).

For each CASP target in our benchmark set there is a highly variable number of related (i.e. similar) templates in ECOD. In particular, the number of related templates per target varies from 2 to 3540 for CASP12 targets, and from 1 to 1512 for CASP13 targets. In Table 3 we assess fold recognition performances by considering only the top-scored templates, but this does not tell us how all the templates related to a given target are ranked during a search. In Figure 1, we show the ranking distributions of all the templates related to the CASP12 and CASP13 targets. The probability density functions in Figure 1 are estimated from the observed rankings in our tests for both CASP12 and CASP13 targets, using the *density* function available in R. It is clear that the congruence coefficient shifts the ranking distribution of related templates closer towards 1, uniformly for all methods. However, in Figure 1 we can see that the peak of the *r*_*c*_-related distributions is around ranking position 607, which is still quite far from the top-20 interval considered in Table 3.

**Figure 1:**
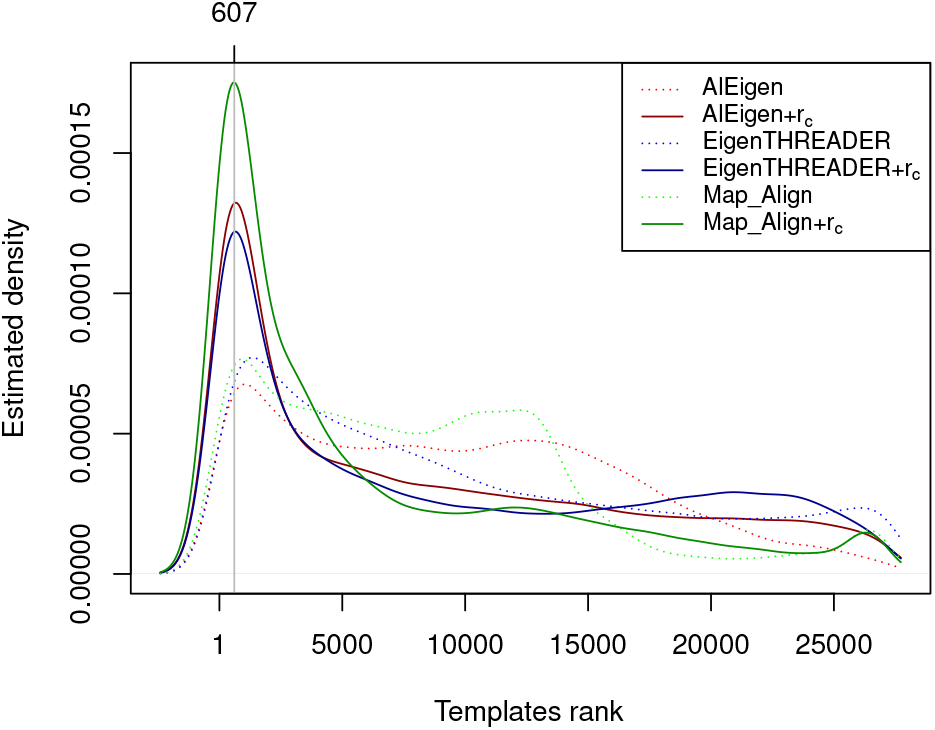
Estimated ranking distribution of templates searched with predicted contacts. Comparison of the ranking distribution for templates related to the target proteins in CASP12 and CASP13.

A more stringent analysis of the fold recognition performances can be done on CASP targets that contain at least one FM (Free Modeling) domain. There the fold recognition performances are assessed only for the FM domains of such targets. The TPR performances are summarized in Table 4. We do not consider fold recognition at the family level, since FM targets are classified as completely new folds^1^. To improve readability, we show only the results for the top-20 recovered templates (complete results in Section 7.4.1, Suppl). With Map_Align+*r*_*c*_ we can exceed 40% fold recognition accuracy on FM targets at the topology level for both CASP12 and CASP13. While leaving room for improvements, such performance is still interesting. Recall that for both CASP12 and CASP13, the ECOD database was pre-filtered by removing all domains that share a significant sequence similarity with the CASP targets. Hence, although we do not have many FM targets in our benchmark sets, the results in Table 4 suggest that contact prediction is at a sufficiently high level of accuracy to improve fold recognition for distantly related homologs.

**Table 4:**
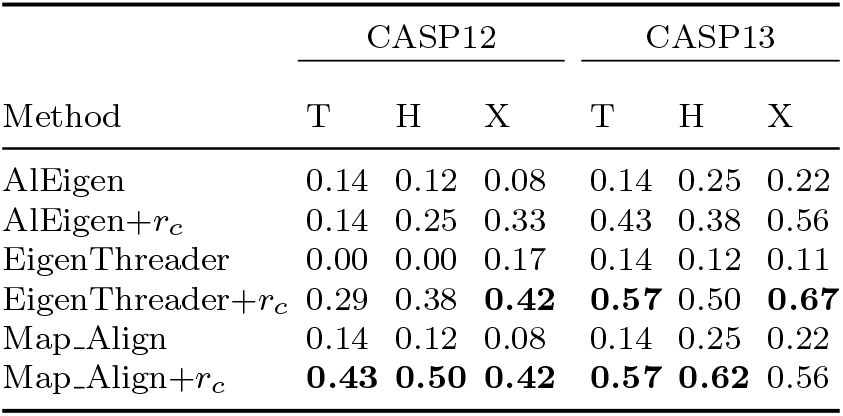
Fold recognition performances with predicted contacts on FM targets (top-20 hits). ECOD hierarchy: (T) Topology Level (7 targets in both CASP12 and CASP13), (H) Homology Level (8 targets in both CASP12 and CASP13), (X) Possible Homology Level (12 targets in CASP12, 9 targets in CASP13). Best TPR performances per column are highlighted in bold

| Method | CASP12 |  |  | CASP13 |  |  |
| --- | --- | --- | --- | --- | --- | --- |
|  | T | H | X | T | H | X |
| AlEigen | 0.14 | 0.12 | 0.08 | 0.14 | 0.25 | 0.22 |
| AlEigen+$r_c$ | 0.14 | 0.25 | 0.33 | 0.43 | 0.38 | 0.56 |
| EigenThreader | 0.00 | 0.00 | 0.17 | 0.14 | 0.12 | 0.11 |
| EigenThreader+$r_c$ | 0.29 | 0.38 | **0.42** | **0.57** | 0.50 | **0.67** |
| Map_Align | 0.14 | 0.12 | 0.08 | 0.14 | 0.25 | 0.22 |
| Map_Align+$r_c$ | **0.43** | **0.50** | **0.42** | **0.57** | **0.62** | 0.56 |

Finally, we look at the quality of the alignments provided by the three methods. We measure the alignment quality through the *Z*-score of the congruence coefficient between two aligned maps, where the mean and standard deviation are computed over all possible alignments between the two maps. The alignment quality measure is independent of the similarity between the two maps being aligned: optimal alignments can be computed for two unrelated maps, and poor alignments can be computed for similar maps, which may affect fold recognition performances. In Figure 2, we plot the Z-score distribution of all the alignments between CASP12/CASP13 targets and templates maps. The *Z*-scores are computed using the true mean of the congruence coefficient over all possible alignments, and the sampled standard deviations (Section 7.3, Suppl.). The Z-score distributions are computed separately for all the alignments between a target map and all its related templates in ECOD, and between a target map and all unrelated templates. First of all, in Figure 2 we can notice that, independently of the chosen method, the alignment *Z*-score distribution is similar for related and unrelated templates. This indicates that the alignment quality of each method is independent of the similarity between the two input maps, i.e. on average one cannot expect to see better alignments for related maps than for unrelated maps. Overall, Map_Align provides better alignments that the two other methods. This is consistent with the performances reported in Table 3, where Map_Align, especially with the correlation coefficient as its scoring function, achieves overall best fold recognition accuracy. In contrast, EigenTHREADER provides on average lower quality alignments. In particular, in a non-trivial number of cases the *r*_*c*_ scores with respect to the alignments computed by EigenTHREADER are lower that the expected mean. However, EigenTHREADER’s fold recognition performances are no dramatically affected when coupled with the congruence coefficient. This is further evidence that the congruence coefficient provides an effective measure of map similarity.

**Figure 2:**
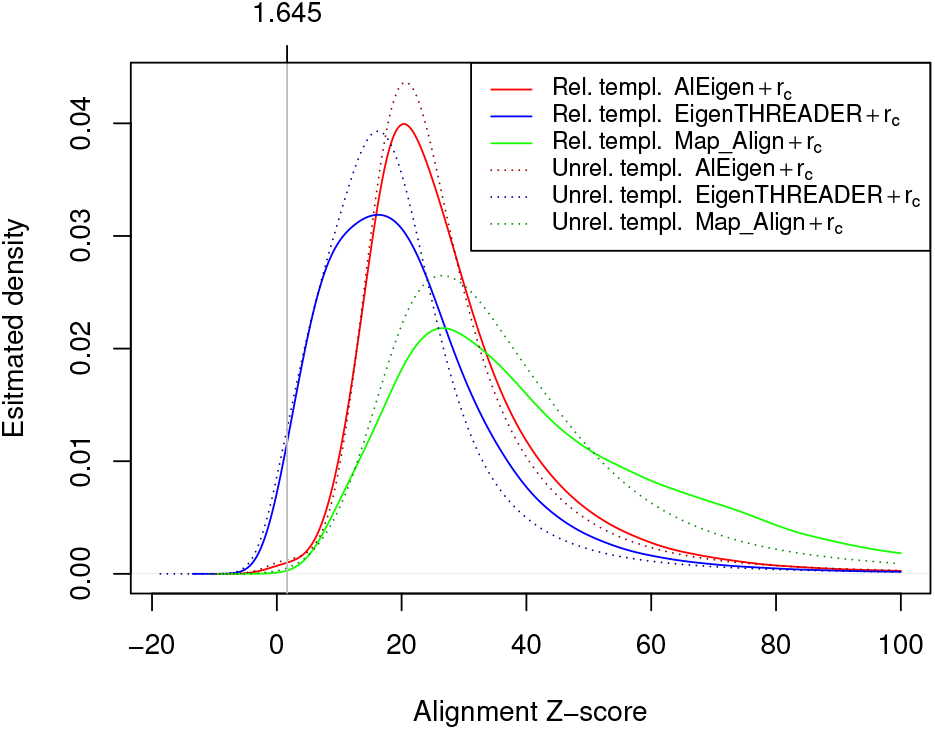
Estimated Z-score distribution of contact map alignments. Comparison of the Z-score distribution for templates related and unrelated to the target proteins in CASP12 and CASP13.

### 3.2 Fold recognition with predicted distances

For distance maps, we run tests similar to those performed with contact maps. Here we use two structural alignment tools, CE and TM-Align. Also here we compare the fold recognition capabilities of CE and TM-Align with their own scoring schemes, CE’s Z-score and TM-score, respectively, against those obtained by using the congruence coefficient, CE+*r*_*c*_ and TM-Align+*r*_*c*_, respectively. The goal of these tests is to show whether the congruence coefficient is suitable also for distance map comparisons and thus for distance map-based fold recognition. Furthermore, this provides a preliminary comparison between contact-based versus distance-based fold recognition.

The true positive rate performances on the CASP12 and CASP13 sets are summarized in Table 5, while performances on the FM targets are in Table 6 (Section 7.4.2, Suppl. for the complete table). Unlike the contact results, in these tests fold recognition capabilities are mainly affected by the quality of the predicted structures. In particular, the overall fold recognition performance for CASP13 is better than for CASP12, a direct consequence of the improvements in protein structure prediction reported at CASP13. Furthermore, the restriction to structural predictions by the top performing methods only does improve fold recognition on CASP12, but not on CASP13 (Section 7.4.2, Suppl.).

**Table 5:**
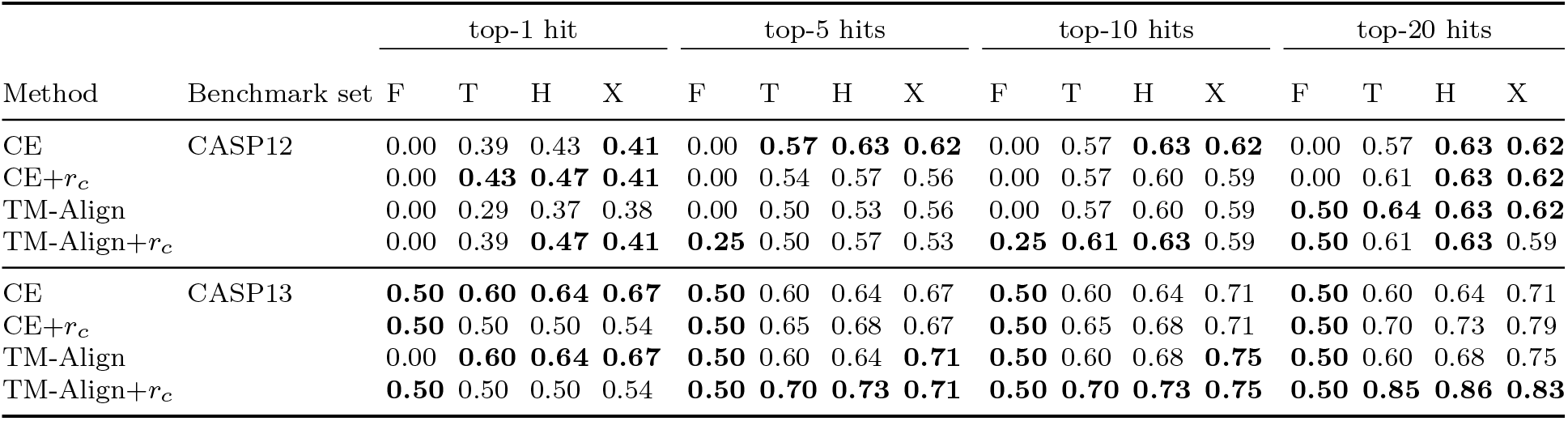
Fold recognition performances with predicted distances/structures. True Positive Rate (TPR) fold recognition performances on CASP12 and CASP13 benchmark sets. The TPR performances are assessed with respect to the **top-1**, **top-5**, **top10** and **top-20** ranked hits. ECOD hierarchy: (F) Family Level (4 targets in CASP12, 2 targets in CASP13), (T) Topology Level (28 targets in CASP12, 20 targets in CASP13), (H) Homology Level (30 targets in CASP12, 22 targets in CASP13), (X) Possible Homology Level (34 targets in CASP12, 24 targets in CASP13). CE and TM-Align use their own scoring system. CE+*r*_*c*_, and TM-Align+*r*_*c*_ use statistically significant congruence coefficient. Best TPR performances per column on CASP12 and CASP13 benchmark sets are highlighted in bold.

**Table 6:**
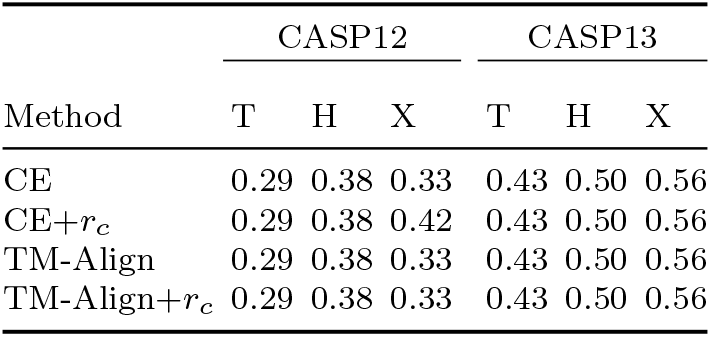
Fold recognition performances with predicted distances/structures on FM targets (top-20 hits). ECOD hierarchy: (T) Topology Level (7 targets in both CASP12 and CASP13), (H) Homology Level (8 targets in both CASP12 and CASP13), (X) Possible Homology Level (12 targets in CASP12, 9 targets in CASP13).

| Method | CASP12 |  |  | CASP13 |  |  |
| --- | --- | --- | --- | --- | --- | --- |
|  | T | H | X | T | H | X |
| CE | 0.29 | 0.38 | 0.33 | 0.43 | 0.50 | 0.56 |
| CE+$r_c$ | 0.29 | 0.38 | 0.42 | 0.43 | 0.50 | 0.56 |
| TM-Align | 0.29 | 0.38 | 0.33 | 0.43 | 0.50 | 0.56 |
| TM-Align+$r_c$ | 0.29 | 0.38 | 0.33 | 0.43 | 0.50 | 0.56 |

In terms of fold recognition performances, the congruence coefficient is comparable to TM-score and CE’s Z-score, two metrics adopted by CASP. In some cases, the congruence coefficient achieves slightly better accuracy, such as on the CASP13 benchmark set for top-5 hits and above. However, we do not observe a significantly strong differences between the TM-Align vs CE, and TM-score/Z-score vs congruence coefficient, since no approach is overall better than another in all cases. This is more evident in Figure 3, which shows that the ranking distributions of related templates are practically undistinguishable among all benchmarked approaches.

**Figure 3:**
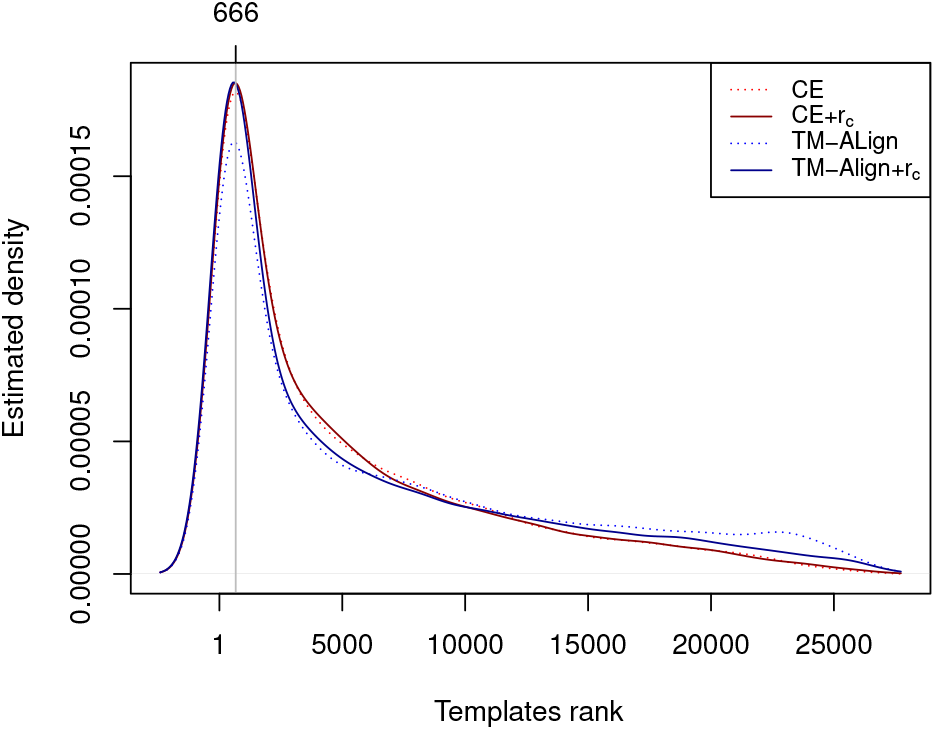
Estimated ranking distribution of templates searched with predicted structures/distances. Comparison of the ranking distribution for templates related to the target protein. Probability density function estimated from observed rankings in CASP12 and CASP13 benchmark sets.

In Figure 4, we show the *Z*-scores distributions of the alignments provided by CE and TM-Align. Unlike map alignment tools, structural alignment tools tend to compute slightly better alignments between a target and its related templates than against unrelated templates. However, not surprisingly, in most of the cases the *r*_*c*_ coefficients related to such alignments are lower than the expected coefficient over all possible alignments. This is because CE and TM-Align perform local alignments, while the maximum *r*_*c*_ score between two maps is achieved by performing a global alignment. Specific distance map alignment tools may provide better global alignments and may further improve fold recognition with predicted distances. Although most of the local alignments computed by CE and TM-Align are not optimal global alignments, the *r*_*c*_ p-value is generally statistically significant, due to the large degree of freedom associated with distance maps. Thus, unlike the case of contact-based searches, the p-value cutoff for distance maps does not seem to improve database searches.

**Figure 4:**
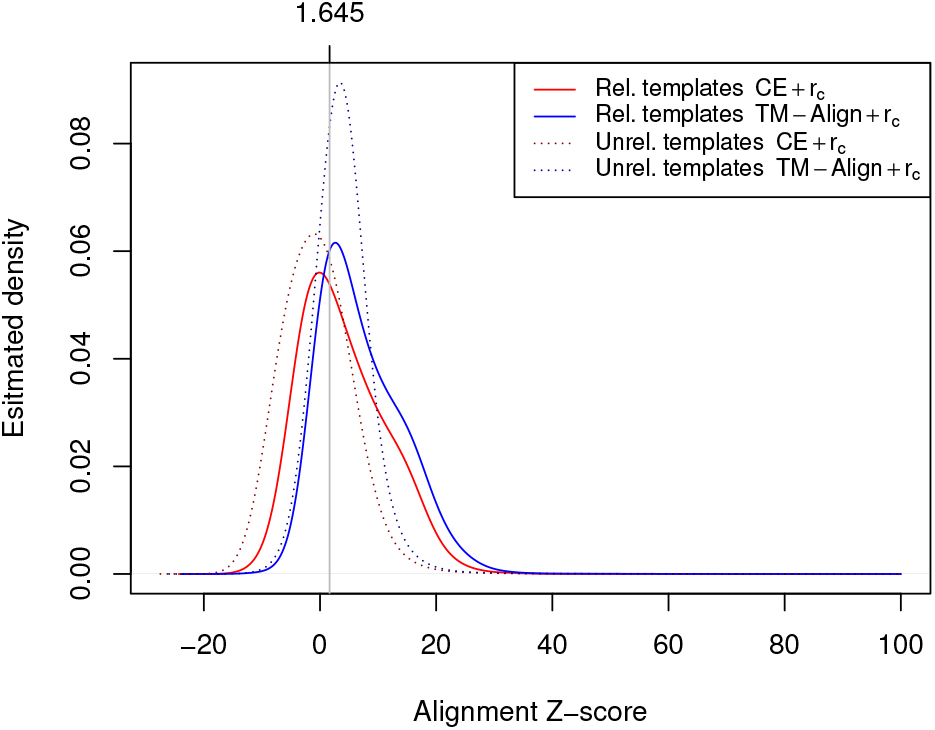
Estimated Z-score distribution of distance map alignments. Comparison of the Z-score distribution for templates related and unrelated to the target proteins in CASP12 and CASP13.

Finally, distance-based fold recognition does not outperform contact-based fold recognition. If anything, the converse is true when looking at Tables 4 and 6 summarizing fold recognition performances on FM targets. While the limited number of targets in our benchmark sets does not allow one to draw strong conclusions, these tests at least confirm that contact map comparison is a valuable approach for detecting protein structure similarities.

## 4 Conclusion

We exploited the congruence coefficient as a measure for detecting map similarities. We proved that the congruence coefficient has several important mathematical properties allowing one to rigorously assess its statistical significance and efficiently compute its average and standard deviation. We compared contact map-based and distance map-based fold recognition performances of the congruence coefficient against those of contact map alignment and structural alignment tools. Overall, the congruence coefficient score improves the fold recognition accuracy, particularly for contact-based fold recognition, proving its effectiveness as a general similarity metric for protein map comparisons. Furthermore, contact-based fold recognition accuracy is comparable or better than distance/structure-based fold recognition, suggesting its potential as a general approach for improving the detection of protein structure similarities.

## Acknowledgements

Work partially supported by the EU Horizon 2020 research and innovation programme under Marie Curie grant 777822 to PDL, and NIH grant GM123558 to PB.

## Footnotes

1 For the sake of completeness, we report that two CASP12 targets have been annotated at the Family level in ECOD. These two cases were ignored, not withstanding that no approach can detect the templates related to such targets.

