## SupplementaryFile for "Fold recognition by scoring protein map similarities using the congruence coefficient"

### Mathematical properties of the congruence coefficient

#### 1 Introduction

The congruence coefficient is a measure of matrix similarity. It was first introduced in [Burt, 1948], with the name of *unadjusted correlation*, and later popularized by Tucker [Tucker, 1951] as a measure of similarity in factor analysis. The congruence coefficient score is defined as the trace of the product between two normalized matrices. Its values range between -1 and +1, where values close to 1 indicate a high degree of similarity, and values close to 0 indicate low similarity. Other than for the range of values, the congruence coefficient shares some more similarities with the Pearson's correlation, where the main difference is that the latter measures the deviation for the mean, while the former measures the deviation from zero.

Here we exploit the congruence coefficient as a measure of similarity between (symmetric) map representations of protein structures, such as contact maps and distance maps. In particular, we prove several interesting statistical properties of the congruence coefficient: i) polynomial-time computation of its expectation and variance over all possible (exponential number of) alignments between two symmetric matrices; ii) closed formulas for computing its expectation and variance over all possible (exponential number of) permutations of one

*Department of Computer Science and Engineering, University of Bologna, Italy
†Department of Computer Science, University of California, Irvine, CA 92697
‡Institute for Genomics and Bioinformatics, University of California, Irvine, CA 92697, USA

of the two matrices; iii) P-value of the congruence coefficient between two aligned (i.e. same size) symmetric matrices. While our focus is mainly on the comparison of protein map representations, all these statistical properties hold for general symmetric matrices. In more details,

i) the expectation of the congruence coefficient over all possible alignments between two symmetric matrices $X \in \mathbb{R}^{m\times m}$ and $Y \in \mathbb{R}^{n\times n}$ can be obtained by computing the *expectation matrix* $E[Y^\alpha] \in \mathbb{R}^{m\times m}$, where $Y^\alpha \in \mathbb{R}^{m\times m}$ is the $Y$ matrix recoded with respect to alignment $\alpha$. The (normalized) trace of the product between $X$ and expectation matrix $E[Y^\alpha]$ gives the expectation of the congruence coefficient over all possible alignments between $X$ and $Y$. The expectation matrix $E[Y^\alpha]$ can be computed by just using $Y$ and size $m$ (i.e. we just need to know the size $m$ of $X$ but not $X$ itself). Equivalently, the variance of the congruence coefficient over all possible alignments between $X$ and $Y$ depends on the *variance-covariance matrix* $Var[Y^\alpha] \in \mathbb{R}^{m^2\times m^2}$, which again can be computed by just using $Y$ and size $m$. The expectation and variance-covariance matrix over all alignments cannot be easily recoded for the Pearson correlation, since, differently from the congruence coefficient, its normalization factor is not invariant with respect to the alignment size.

ii) expectation and variance over all permutations have a nice characterization in terms of closed formulas that roughly depend on the sum of elements in the $X$ and $Y$ matrices. Such statistics can be computed much more quickly than the alignment-related statistics. The approach adopted for computing expectation and variance over permutations has been previously developed in [Kazi-Aouala *et al.*, 1995] for several matrix-related metrics but not the congruence coefficient. In this case, such formulas can be easily recoded also for the Pearson correlation, since its normalization factor is invariant with respect to any matrix permutation.

iii) the congruence coefficient $P$-value of the statistical hypothesis testing, under the null hypothesis that the coefficient is zero, can be recoded as a statistical hypothesis testing on the angle between two vectors in the $N$-dimensional unit sphere (where $N$ depends on the size of the two matrices), under the null hypothesis that the two vectors are orthogonal.

These statical properties can be used to assess the statistical quality and significance of an alignment between two protein maps. In particular,

- the expectation and variance over all possible alignments between two matrices can be used to compute the $Z$-score of any given alignment between the two matrices, which gives an indication of the quality of the alignment. The distribution of the congruence coefficient over all possible alignments $\alpha$ between two symmetric matrices $X \in \mathbb{R}^{m \times m}$ and $Y \in \mathbb{R}^{n \times n}$ depends essentially on the multivariate distribution of the (aligned) random matrices $Y^\alpha \in \mathbb{R}^{m \times m}$. It is not easy to characterize such distribution, though it doubtfully follows a multivariate normal distribution, since the $Y^\alpha$ components are not independent. By empirical observations, the distribution of the congruence coefficients over all alignments between contact maps seems to be skewed to the right, thus its right tail is probably not well-approximated by a normal distribution. Although this can affect the estimation of the exact P-value for the $Z$-score, the $Z$-score alone (computed from the *exact* mean and standard deviation) can be considered as highly accurate in detecting bad-quality alignments. Unfortunately, while the expectation matrix $E[Y^\alpha]$ can be computed quickly for native protein contact/distance maps, the computation of the variance-covariance matrix $Var[Y^\alpha]$ is more challenging, since its computational complexity is quartic in the size $mn$. The $Var[Y^\alpha]$ matrices are also not easily approximable and quite large. Anyway, we can show that an ad-hoc sampling of random alignments can give almost exact estimation of the true variance over all possible alignments.

- the P-value of a congruence coefficient gives the probability of observing an equivalent or higher coefficient if one of the two maps is chosen at random (in the space of all possible symmetric matrices with same size). It can be used to detect whether there is a statistically significant similarity between two aligned maps. Note that, a high congruence coefficient score does not always indicates a significant similarity between two matrices, since the probability of observing it depends on the topology of the two matrices. An extreme example is the comparison between a constant (non-zero) matrix with a random (non-zero) matrix. In such cases, the congruence coefficient will tend to be always quite high but it will be statistically significant only if the second map is almost constant too.

The congruence coefficient's $P$-value is complementary to the $Z$-score of the alignment, since the latter indicates whether an alignment between two maps is poor or good while the former indicates whether there is a statistically significantly similarity between the aligned maps. The combination of both can be used to improve/analyze database searches based on map similarities. For example, a non-significant P-value does not necessarily imply low similarity between two maps, if the $Z$-score of the alignment between the two maps is low.

To conclude, we remark that expectation and variance over permutations do not generally provide biologically meaningful statistical information when comparing map representation of protein structures, since random permutations of protein contact/distance maps are typically not consistent with any three-dimensional set of points. We exploited permutation statistics as a possible (fast) approach for approximating variance calculation over all possible alignments. Unfortunately, test comparisons on real protein contact/distance maps show that both expectation and variance over permutations poorly approximate expectation and variance over all possible alignments. Thus, except for the theoretical interest of such formulas, they do not have an immediate application in protein map comparison. We included them here since they could still be be useful in different applications of the congruence coefficient where matrix permutations are meaningful.

The document is organized as follows. In Section 2 we give the formal definition of congruence coefficient for arbitrary matrices and introduce some of its general properties. In Section 3, we focus on symmetric matrices and give the formal definition of congruence coefficient between aligned symmetric matrices, by introducing also the notation that will be used throughout the rest of the document. Section 4 is devoted to expectation and variance of the congruence coefficient over all possible alignments between two symmetric matrices, while Section 5 is devoted to expectation and variance over all permutations. Section 6 formalizes statistical hypothesis testing of the congruence coefficient. To conclude, in Section 7 we show some experimental tests on a real biological data. In particular, in Section 7.3 we compare true expectation/variance over all possible alignments vs expectation/variance obtained by sampling and permutations. In the remaining subsections, we show some applications of the congruence coefficient for protein fold recognition.

### 2 Definition of the congruence coefficient

We give the formal definition of the congruence coefficient and introduce some basic properties.

**Definition 2.1.** *Let $X, Y \in \mathbb{R}^{m\times n}$ two matrices, the congruence coefficient $r_c$ between $X$ and $Y$ is defined by*

$$r_c(X,Y) = \frac{tr(XY^T)}{\sqrt{tr(XX^T)tr(YY^T)}} \tag{1}$$

where $Y^T$ is the transpose of $Y$ and the trace of the product is defined by

$$tr(XY^T) = \sum_{i=1}^{m}\sum_{j=1}^{n} X_{ij}Y^T_{ij}$$

The denominator in Equation 1 is a normalization factor that shifts the trace of the product into the interval $[-1, 1]$.

It shares some similarities with the Pearson's correlation coefficient but it has also important differences.

1. Similarly to the Pearson correlation, the $r_c$ coefficient is a continuous function, and can be seen as the cosine of the angle between the vectorized versions of the (normalized) $X$ and $Y$ matrices.

2. Similarly to the Pearson correlation, by the Cauchy-Schwarz inequality, the $r_c$ coefficient ranges in the $[-1, 1]$ interval, where coefficients close to 1 indicate a high degree of matrix similarity, and coefficients close to 0 a low degree of similarity. Negative coefficients indicate negative correlation, which can be reversed by simply changing the sing of all the elements in one of the two matrices $X$ and $Y$.

3. Similarly to the Pearson correlation, the congruence coefficient is invariant (up to the sign) to scalar multiplication of the matrices X, Y with constant factors different from zero.

   Let $X, Y \in \mathbb{R}^{m\times n}$ two matrices and $a, b \in \mathbb{R}$ two constant values different from zero.

$$\begin{aligned} r_c(aX, bY) &= \frac{tr(abXY^T)}{\sqrt{tr(a^2XX^T)}\sqrt{tr(b^2YY^T)}} = \\ &= \frac{ab \cdot tr(XY^T)}{|a|\,|b| \cdot \sqrt{tr(XX^T)}\sqrt{tr(YY^T)}} = \\ &= \operatorname{sgn}(ab)\frac{tr(XY^T)}{\sqrt{tr(XX^T)}\sqrt{tr(YY^T)}} \end{aligned}$$

4. Similarly to the Pearson correlation, the congruence coefficient is symmetric

$$r_c(X, Y) = r_c(Y, X)$$

5. The Pearson correlation measures the deviations from the *mean*, while the congruence coefficient measures the deviations from *zero*. This is a fundamental property of the congruence coefficient. We will show that the normalization factor of the congruence coefficient is invariant with respect to any possible alignment between the two input matrices, where an aligned map can be seen as a matrix containing additional zero rows/columns. Such property does not hold for the Pearson correlation.

   In fact, the matrix version of the Pearson correlation is

$$r(X, Y) = \frac{tr((X - \mu_X)(Y - \mu)^T)}{\sqrt{tr((X - \mu_X)(X^T - \mu_X))tr((Y - \mu_Y)(Y - \mu_Y)^T)}} \tag{2}$$

   where

$$\mu_X = \frac{1}{mn}\sum_{i=1}^{m}\sum_{j=1}^{n} X_{ij} \text{ and } \mu_Y = \frac{1}{mn}\sum_{i=1}^{m}\sum_{j=1}^{n} Y_{ij}$$

depend on the size $mn$ and change if we increase the size of the $X$ and $Y$ matrices by introducing zero rows/columns (i.e. if we align the two matrices, see Section 3).

6. The counter effect of the previous property is that, differently from the Pearson's correlation coefficient, the congruence coefficient is sensitive to additive constants. However, the new congruence coefficient under additive constant can be easily characterized. We first need to introduce some notation.

   **Definition 2.2.** *Let $\mathbf{1}_{mn} \in \{1\}^{m\times n}$ denote the $m\times n$ all-one matrix and $c\mathbf{1}_{mn} \in \{c\}^{m\times n}$ the constant matrix where every element is $c \in \mathbb{R}$.*

   **Definition 2.3.** *Let $X \in \mathbb{R}^{m\times n}$ be a matrix. We denote with*

$$\Sigma(X) = tr(X\mathbf{1}_{mn}^T) = tr(\mathbf{1}_{mn}X^T) = \sum_{i=1}^{m}\sum_{j=1}^{n} X_{ij}$$

   *the trace of the product between $X$ and the all-one matrix $\mathbf{1}_{mn}$, which is equivalent to the sum of all elements of $X$.*

   Let $X, Y \in \mathbb{R}^{m\times n}$ two matrices and $a, b \in \mathbb{R}$ two constant values.

$$\begin{aligned} r_c(X + a\mathbf{1}_{mn}, Y + b\mathbf{1}_{mn}) &= \frac{tr(XY^T + bX\mathbf{1}_{mn}^T + a\mathbf{1}_{mn}Y^T + ab\mathbf{1}_{mn}\mathbf{1}_{mn}^T)}{\sqrt{tr(XX^T + 2aX\mathbf{1}_{mn}^T + a^2\mathbf{1}_{mn}\mathbf{1}_{mn}^T)}\sqrt{tr(YY^T + 2bY\mathbf{1}_{mn}^T + b^2\mathbf{1}_{mn}\mathbf{1}_{mn}^T)}} = \\ &= \frac{tr(XY^T) + b\Sigma(X) + a\Sigma(Y) + abmn}{\sqrt{tr(XX^T) + 2a\Sigma(X) + a^2mn}\sqrt{tr(YY^T) + 2b\Sigma(Y) + b^2mn}} = \\ &= \mathcal{W}_{ab}(X, Y) \cdot r_c(X, Y) + \mathcal{K}_{ab}(X, Y) \end{aligned}$$

   where the two coefficients

$$\mathcal{W}_{ab}(X, Y) = \frac{\sqrt{tr(XX^T)}\sqrt{tr(YY^T)}}{\sqrt{tr(XX^T) + 2a\Sigma(X) + a^2mn}\sqrt{tr(YY^T) + 2b\Sigma(Y) + b^2mn}} \tag{3}$$

and

$$\mathcal{K}_{ab}(X,Y) = \frac{b\Sigma(X) + a\Sigma(Y) + abmn}{\sqrt{tr(XX^T) + 2a\Sigma(X) + a^2mn}\sqrt{tr(YY^T) + 2b\Sigma(Y) + b^2mn}} \tag{4}$$

are defined by the constants $a, b$, the matrix size $mn$, the total sum of their entries $\Sigma(X), \Sigma(Y)$, and the traces of their self products $tr(XX^T)$, $tr(YY^T)$. When $a = b = 0$, we have that

$$\mathcal{W}_{00}(X,Y) = 1 \text{ and } \mathcal{K}_{00}(X,Y) = 0$$

While the $r_c$ coefficient can be computed for arbitrary matrices, in the following we consider only symmetric matrices, i.e. matrix representations of protein structure information. In particular, we consider: i) *protein distance maps*, which are defined as symmetric matrices $X$ in which the entry $X_{ij}$ encodes the Euclidean distance between protein residues $i$ and $j$ in the protein three-dimensional structure; ii) *protein contact maps*, which are defined as symmetric binary matrices $X$ in which the entry $X_{ij}$ is 1 if and only if the the Euclidean distance between protein residues $i$ and $j$ is below some threshold (typically 8Å). Although our focus is on matrix representations of protein structures, all the results in the following sections hold for general symmetric matrices.

### 3 Congruence coefficient between aligned maps

The congruence coefficient in Equation 1 can only be applied to matrices of the same size. Symmetric matrices of different sizes need to be *aligned* in order to assess their similarity through the congruence coefficient. Here we formally define matrix alignment and develop a notation that will be used in the following sections.

Since our focus is on matrix representations of protein structure information, we only consider matrix-alignments that are induced by *global sequence alignments*. In particular, we consider all possible alignments between two sequences that respect the following two general assumptions:

1. a gap cannot be matched with a gap (non-boring gaps),
2. there is a unique alignment that matches both (sub)sequences with only gaps.

Assumptions 1 and 2 impose a finite number of possible alignments between two sequences. Furthermore, they bound the maximum length of an alignment between two sequences of length $m$ and $n$ to the sum of their lengths, $m+n$. Conversely, the minimum length of an alignment is bounded by the longest sequence, $\max\{m,n\}$.

**Definition 3.1.** *A partial function*

$$\alpha : \{1,..,m\} \to \{1,..,n\}$$

*is an* **alignment** *if* $\forall i \neq j \in \{1,..,m\}$ *such that* $\alpha(i) \neq \perp \neq \alpha(j)$ *then*

$$\alpha(i) < \alpha(j) \iff i < j$$

□

The notation $\alpha(i) = \perp$ indicates that $\alpha$ is not defined on $i$. Since and alignment is a partial injective function, the inverse function, defined by $\alpha^{-1} : \{1,..,n\} \to \{1,..,m\}$, exists and it is also an alignment. We denote by $|\alpha|$ the number of matched positions in both sequences by $\alpha$, i.e.

$$|\alpha| = |\{i \mid \alpha(i) \neq \perp\}|.$$

Now, let $X \in \mathbb{R}^{m\times m}, Y \in \mathbb{R}^{n\times n}$ be two symmetric matrices. From the matrix point of view, an alignment between $X$ and $Y$ can be induced by a sequence alignment between a sequence of length $m$ and one of length $n$. In particular, an alignment $\alpha$ can be seen as a transformation of two symmetric matrices, $X$ and $Y$, that adds some zero rows and (corresponding) zero columns, such that the transformed matrices are symmetric and have the same size. The zero rows/columns correspond to gaps in the alignment. We can define this more formally.

**Definition 3.2.** *Let* $X \in \mathbb{R}^{m\times m}, Y \in \mathbb{R}^{n\times n}$ *be two symmetric matrices and consider some alignment* $\alpha : \{1,..,m\} \to \{1,..,n\}$ *between two sequences of length m and n, respectively. Let* $\sigma = (\sigma_1, \sigma_2)$ *be two surjective partial functions*

$$\sigma_1 : \{1,..,N\} \to \{1,..,m\},\ \sigma_2 : \{1,..,N\} \to \{1,..,n\}$$

*such that*

$$\sigma_2 \circ \sigma_1^{-1} = \alpha \text{ and } \sigma_1 \circ \sigma_2^{-1} = \alpha^{-1}$$

*and (non-boring gap condition)*

$$N = m + n - |\alpha|$$

*The symmetric matrices* $X^{\sigma_1}, Y^{\sigma_2} \in \mathbb{R}^{N\times N}$ *induced by alignment* $\alpha$ *are defined by*

$$X^{\sigma_1}_{ij} = \begin{cases} 0 & \text{if } \sigma_1(i) = \perp \text{ or } \sigma_1(j) = \perp \\ X_{\sigma_1(i),\sigma_1(j)} & \text{otherwise} \end{cases}$$
$$Y^{\sigma_2}_{ij} = \begin{cases} 0 & \text{if } \sigma_2(i) = \perp \text{ or } \sigma_2(j) = \perp \\ Y_{\sigma_2(i),\sigma_2(j)} & \text{otherwise} \end{cases}$$

□

Let $X \in \mathbb{R}^{m\times m}, Y \in \mathbb{R}^{n\times n}$ be two symmetric matrices, $\alpha$ an alignment of length $N$ and $\sigma$ as defined in Definition 3.2. We define the congruence coefficient with respect to the alignment $\alpha$ by

$$r^{\alpha}_{c}(X,Y) = \frac{tr(X^{\sigma_1}Y^{\sigma_2})}{\sqrt{tr(X^{\sigma_1}X^{\sigma_1})tr(Y^{\sigma_2}Y^{\sigma_2})}} = \frac{tr(X^{\sigma_1}Y^{\sigma_2})}{\sqrt{tr(XX)tr(YY)}} \tag{5}$$

Note that, by definition of aligned matrix $X^{\sigma_1}$ (resp. for $Y^{\sigma_2}$), the normalization factor in Equation 5 is invariant with respect to all possible alignments, i.e.

$$\forall \alpha, tr(XX) = tr(XX^T) = tr(X^{\sigma_1}X^{\sigma_1})$$

Such property does not hold for Pearson's correlation which measures the deviation from the mean value and is thus affected by the number of zero rows and columns introduced in the aligned matrix. Thus, the alignment that maximizes the $r_c$ coefficient in Equation 5 is simply the alignment that maximizes the trace of the product between the aligned matrices.

**Example 3.3.** *Let* $X \in \mathbb{R}^{3\times 3}$ *and* $Y \in \mathbb{R}^{4\times 4}$ *be two symmetric matrices*

$$X = \begin{pmatrix} 0 & 1 & 2 \\ 1 & 0 & 3 \\ 2 & 3 & 0 \end{pmatrix} \quad Y = \begin{pmatrix} 0 & 4 & 5 & 6 \\ 4 & 0 & 7 & 8 \\ 5 & 7 & 0 & 9 \\ 6 & 8 & 9 & 0 \end{pmatrix}$$

*Let the alignment* $\alpha : \{1, ..3\} \to \{1, ..., 4\}$ *be defined by*

$$\alpha(1) = 2, \alpha(2) = \perp, \alpha(3) = 3$$

*and* $\sigma = (\sigma_1, \sigma_2)$*, with* $\sigma_1 : \{1, ..., 5\} \to \{1, .., 3\}$ *and* $\sigma_2 : \{1, ..., 5\} \to \{1, .., 4\}$*, be defined by*

$$\sigma_1(1) = \perp, \sigma_1(2) = 1, \sigma_1(3) = 2, \sigma_1(4) = 3, \sigma_1(5) = \perp$$

$$\sigma_2(1) = 1, \sigma_2(2) = 2, \sigma_2(3) = \perp, \sigma_2(4) = 3, \sigma_2(5) = 4$$

*Then the two aligned matrices* $X^{\sigma_1}$ *and* $Y^{\sigma_2}$ *(the gap rows/columns are in blue) are*

$$X^{\sigma_1} = \begin{pmatrix} \mathbf{0} & \mathbf{0} & \mathbf{0} & \mathbf{0} & \mathbf{0} \\ \mathbf{0} & 0 & 1 & 2 & \mathbf{0} \\ \mathbf{0} & 1 & 0 & 3 & \mathbf{0} \\ \mathbf{0} & 2 & 3 & 0 & \mathbf{0} \\ \mathbf{0} & \mathbf{0} & \mathbf{0} & \mathbf{0} & \mathbf{0} \end{pmatrix} \quad Y^{\sigma_2} = \begin{pmatrix} 0 & 4 & \mathbf{0} & 5 & 6 \\ 4 & 0 & \mathbf{0} & 7 & 8 \\ \mathbf{0} & \mathbf{0} & \mathbf{0} & \mathbf{0} & \mathbf{0} \\ 5 & 7 & \mathbf{0} & 0 & 9 \\ 6 & 8 & \mathbf{0} & 9 & 0 \end{pmatrix}$$

*The congruence coefficient between the two aligned matrices is*

$$r_c^\alpha(X,Y) = \frac{tr(X^{\sigma_1}Y^{\sigma_2})}{\sqrt{tr(X^{\sigma_1}X^{\sigma_1})tr(Y^{\sigma_2}Y^{\sigma_2})}} = \frac{2\times 7 + 2\times 7}{\sqrt{26\times 542}} = 0.236$$

*Note that, the normalization factor of the congruence coefficient is invariant with respect to alignment $\alpha$:*

$$tr(XX) = 26 = tr(X^{\sigma_1}X^{\sigma_1}) \text{ and } tr(YY) = 542 = tr(Y^{\sigma_2}Y^{\sigma_2})$$

*while the normalization factor of the Pearson correlation (see Equation 2) is not:*

$$tr((X-\mu_X)(X-\mu_X) = 12 \neq 22.24 = tr((X^{\sigma_1}-\mu_{X^{\sigma_1}})(X^{\sigma_1}-\mu_{X^{\sigma_1}}))$$

*and*

$$tr((Y-\mu_Y)(Y-\mu_Y) = 161.75 \neq 298.64 = tr((Y^{\sigma_2}-\mu_{Y^{\sigma_2}})(Y^{\sigma_2}-\mu_{Y^{\sigma_2}}))$$

□

The definition of the congruence coefficient between aligned matrices in Equation 5, is somewhat inconvenient, since it forces us to consider two transformations $X^{\sigma_1}$ and $Y^{\sigma_2}$, which can complicate calculations. However, for ease of mathematical simplification, we can define the congruence coefficient with respect to an alignment in an alternative way, which will greatly simplify the formulas in the following two sections. In detail, given an alignment $\alpha$, an alternative (and equivalent) way to define the trace of the product between the aligned matrice is to leave unchanged the $X$ matrix and just recode the $Y^{\sigma_2}$ matrix by removing all rows/columns that match a gap row/column in $X_1^\sigma$. This leads us to a simpler and equivalent mathematical formulation for the congruence coefficient between aligned matrices. More formally, let $X \in \mathbb{R}^{m\times m}, Y \in \mathbb{R}^{n\times n}$ be two symmetric matrices and consider an alignment $\alpha : \{1,..,m\} \to \{1,..,n\}$ between $X$ and $Y$. We can recode the $Y$ matrix, which we call $Y^\alpha \in \mathbb{R}^{m\times m}$, by

$$Y^\alpha_{ij} = \begin{cases} Y_{\alpha(i)\alpha(j)} & \text{if } \alpha(i) \neq \perp \text{ and } \alpha(j) \neq \perp \\ 0 & \text{otherwise} \end{cases} \tag{6}$$

In the same way, we can define $X^{\alpha^{-1}}$ as the matrix $X^{\sigma_1}$ in which have been removed all rows/columns that match an gap row/column in $Y^{\sigma_2}$.

Since gap rows/columns in $X^{\sigma_1}$ are zero rows/columns, they do not give any contribution in the trace of the matrix product $X^{\sigma_1}Y^{\sigma_2}$. Then, it is easy to see that

$$tr(X^{\sigma_1}Y^{\sigma_2}) = tr(XY^{\alpha})$$

and then

$$r_c^{\alpha}(X,Y) = \frac{tr(X^{\sigma_1}Y^{\sigma_2})}{\sqrt{tr(XX)tr(YY)}} = \frac{tr(XY^{\alpha})}{\sqrt{tr(XX)tr(YY)}} = \frac{tr(X^{\alpha^{-1}}Y)}{\sqrt{tr(XX)tr(YY)}} \tag{7}$$

In this way, when considering the congruence coefficient with respect to some alignment $\alpha$, we can leave unchanged either the $X$ or the $Y$ matrix and recode uniquely the other matrix with respect to the alignment $\alpha$.

**Example 3.4.** *As in Example 3.3, let $X \in \mathbb{R}^{3\times3}$ and $Y \in \mathbb{R}^{4\times4}$ be two symmetric matrices*

$$X = \begin{pmatrix} 0 & 1 & 2 \\ 1 & 0 & 3 \\ 2 & 3 & 0 \end{pmatrix} \qquad Y = \begin{pmatrix} 0 & 4 & 5 & 6 \\ 4 & 0 & 7 & 8 \\ 5 & 7 & 0 & 9 \\ 6 & 8 & 9 & 0 \end{pmatrix}$$

*and $\alpha : \{1, ..3\} \to \{1, ..., 4\}$ the alignment defined by*

$$\alpha(1) = 2, \alpha(2) = \perp, \alpha(3) = 3$$

*The matrices $X^{\alpha^{-1}}$ and $Y^{\alpha}$ with respect to $\alpha$ are (gap row/columns are in blue)*

$$X^{\alpha^{-1}} = \begin{pmatrix} \mathbf{0} & \mathbf{0} & \mathbf{0} & \mathbf{0} \\ \mathbf{0} & 0 & 2 & \mathbf{0} \\ \mathbf{0} & 2 & 0 & \mathbf{0} \\ \mathbf{0} & \mathbf{0} & \mathbf{0} & \mathbf{0} \end{pmatrix} \qquad Y^{\alpha} = \begin{pmatrix} 0 & \mathbf{0} & 7 \\ \mathbf{0} & \mathbf{0} & \mathbf{0} \\ 7 & \mathbf{0} & 0 \end{pmatrix}$$

*The congruence coefficient is then*

$$r_c^{\alpha}(X,Y) = \frac{tr(XY^{\alpha})}{\sqrt{tr(XX)tr(YY)}} = \frac{tr(X^{\alpha^{-1}}Y)}{\sqrt{tr(XX)tr(YY)}} = \frac{2\times7+2\times7}{\sqrt{26\times542}} = 0.236$$

□

### 4 Expectation and variance over all alignments

We show how to compute the expected value and the variance of the $r_c^{\alpha}(X,Y)$ coefficient over all possible alignments $\alpha$ between $X$ and $Y$. Such calculation can be performed in polynomial-time, without actually computing the exponential number of alignments between

$X$ and $Y$. Given two symmetric matrices $X \in \mathbb{R}^{m \times m}$ and $Y \in \mathbb{R}^{n \times n}$, the expected value of $r_c^\alpha(X, Y)$ depends on the *expectation matrix* $E[Y^\alpha] \in \mathbb{R}^{m \times m}$ while the variance of $r_c^\alpha(X, Y)$ depends on the *variance-covariance* matrix $Var[Y^\alpha] \in \mathbb{R}^{m^2 \times m^2}$. In the following sections we will show how to compute such matrices.

Before to show how to compute the expectation and the variance matrices $E[Y^\alpha]$ and $Var[Y^\alpha]$, we show a closed formula for counting the total number of (non-redundant) alignments between two sequences. Such formula is necessary in order to compute expectation and variance matrices.

#### 4.1 Number of possible alignments between two sequences

We show a closed formula for counting the number of possible alignments between two matrices or sequences.

Let us denote with $A(m, n)$ the set of all alignments between two sequences of length $m$ and $n$. We need to count the cardinality of the set $A(m, n)$. Note that, if we choose $k$ distinct positions in two sequences, by definition of alignment, we can have a unique mapping between the $k$ positions in first sequence and the $k$ positions in the second sequence, i.e. we have to match the smaller position in the first sequence with the smaller position in the second sequence, and so on. If the alignment between the two sequences is defined only by such $k$ mapped position, then by the two assumptions 1 and 2, such alignment is unique (i.e. the gap mapping is unique). Now, the number of possible alignments $A_k(m, n)$ that match exactly $0 \leq k \leq \min\{m, n\}$ distinct positions in one sequence of length $m$ with $k$ distinct positions in a second sequence of length $n$ is given by the product between the number of possibile choices of $k$ positions in the first and second sequence:

$$|A_k(m, n)| = \binom{m}{k}\binom{n}{k}$$

Then, the number of distinct alignments between two sequence of length $0 < n \leq m$ is defined by

$$|A(m, n)| = \sum_{k=0}^{n} \binom{m}{k}\binom{n}{k} = \binom{m+n}{n} = \binom{m+n}{m} \tag{8}$$

which follows directly from the identity

$$\binom{n}{n-k} = \binom{n}{k}$$

and the Vandermonde's identity, by substituting $r = n$

$$\sum_{k=0}^{r} \binom{m}{k}\binom{n}{r-k} = \binom{m+n}{r}$$

#### 4.2 Expectation of the congruence coefficient over all possible alignments between two maps

We compute the expected congruence coefficient $E[r_c^\alpha(X,Y)]$ between two symmetric matrices $X$ and $Y$, over all possible alignments $\alpha$ between the two matrices. The approach is purely combinatorial and involves the computation of the *expectation matrix* $E[Y^\alpha] \in \mathbb{R}^{m\times m}$, where $Y^\alpha \in \mathbb{R}^{m\times m}$ is defined in Equation 6 in Section 3.

The expectation of the congruence coefficient is defined by:

$$\begin{aligned}
E[r_c^\alpha(X,Y)] &= \frac{\sum\limits_{\alpha\in A(m,n)} r_c^\alpha(X,Y)}{|A(m,n)|} && = \\
&= \frac{\sum\limits_{\alpha\in A(m,n)} \frac{tr(XY^\alpha)}{\sqrt{tr(XX)tr(YY)}}}{|A(m,n)|} && = \quad \text{(since } \sqrt{tr(XX)tr(YY)} \text{ is constant)} \\
&= \frac{E\left[tr(XY^\alpha)\right]}{\sqrt{tr(XX)tr(YY)}} && = \quad \text{(since } tr \text{ is linear and } X \text{ constant)} \\
&= \frac{tr\left(XE[Y^\alpha]\right)}{\sqrt{tr(XX)tr(YY)}}
\end{aligned}$$

where the *expectation matrix* $E[Y^\alpha]$, which we denote with $\overline{Y}$, is defined by

$$\overline{Y} = E[Y^\alpha] = \frac{\sum\limits_{\alpha\in A(m,n)} Y^\alpha}{|A(m,n)|} \in \mathbb{R}^{m\times m}$$

and, entrywise, it is defined by

$$\overline{Y}_{ij} = \frac{\sum\limits_{\alpha\in A(m,n)} Y_{ij}^\alpha}{|A(m,n)|} = \frac{\sum\limits_{\alpha\in A(m,n),\alpha(i)\neq\perp,\alpha(j)\neq\perp} Y_{\alpha(i)\alpha(j)}}{|A(m,n)|}$$

The $\overline{Y}$ matrix is a symmetric matrix that averages the $Y^\alpha \in \mathbb{R}^{m\times m}$ matrices over all possible alignments $\alpha \in A(m,n)$. We can obtain the expectation matrix of the normalized $Y$ matrix by just dividing each element in $\overline{Y}$ by the constant norm $\sqrt{tr(YY)}$. In order to compute the $\overline{Y}_{ij}$ values we need to distinguish two possible cases.

1. **Diagonal elements**. Let $\alpha$ be an alignment such that

$$\alpha(i) = k, \text{ for some } 1 \le i \le m \text{ and } 1 \le k \le n$$

That is, the alignment $\alpha$ matches the the $i$-th position in the first sequence with the $k$-th position in the second sequence. Then, the summation in the entry $\overline{Y}_{ii}$ contains the value $Y_{kk}$ for a number of times that depends on the total number of alignments $\alpha' \in A(m,n)$ such that $\alpha'(i) = k$. We can compute such number of alignments with the following expression:

$$\binom{s_1+s_2}{s_1}\binom{e_1+e_2}{e_1} = \binom{i-1+k-1}{i-1}\binom{m-i+n-k}{m-i}$$

where

- $s_1 = i-1, s_2 = k-1$ are the lengths of the sub-sequences before $i$ and $k$ in the fist and second sequence, respectively,
- $e_1 = m-i, e_2 = n-k$ are the lengths of the sub sequences starting immediately after $i$ and $k$ up to the end of the first and second sequence, respectively.

Since for every $1 \le i \le m$ and for every $1 \le k \le n$ there is at least one alignment $\alpha \in A(m,n)$ such that $\alpha(i) = k$, we have that

$$\overline{Y}_{ii} = \sum_{k=1}^{n} \frac{\binom{i-1+k-1}{i-1}\binom{m-i+n-k}{m-i}}{\binom{m+n}{n}} Y_{kk} \tag{9}$$

Such summation need to be performed only for the indices $k$ such that $Y_{kk} \neq 0$. When comparing protein maps we usually assume that the diagonal elements are zero. So such term can be ignored and the $\overline{Y}_{ii}$ entries are zero for every index $i$.

2. **Non diagonal elements**. Let $\alpha$ be an alignment such that

$$\alpha(i) = k, \alpha(j) = l, \text{ for some } 1 \le i < j \le m \text{ and } 1 \le k < l \le n$$

Then, the summation in the entry $\overline{Y}_{ij}$ contains the value $Y_{kl}$ for a number of times that depends on the total number of alignments $\alpha$ such that $\alpha(i) = k$ and $\alpha(j) = l$. We can compute such number of alignments with the following expression:

$$\binom{s_1 + s_2}{s_1}\binom{c_1 + c_2}{c_1}\binom{e_1 + e_2}{e_1} =$$

$$\binom{i-1+k-1}{i-1}\binom{j-i-1+l-k-1}{j-i-1}\binom{m-j+n-l}{m-j}$$

where

- $s_1 = i-1, s_2 = k-1$ are the lengths of the sub-sequences before $i$ and $k$, respectively,
- $c_1 = j-i-1, c_2 = l-k-1$ are the lengths of the sub sequences between $i, j$ and $k, l$ , respectively,
- $e_1 = m-j, e_2 = n-l$ are the lengths of the sub sequences starting immediately after $j$ and $l$ up to the end of the first and second sequence, respectively.

Since for every $1 \leq i < j \leq m$ and for every $1 \leq k < l \leq n$ there is at least one alignment $\alpha \in A(m, n)$ such that $\alpha(i) = k, \alpha(j) = l$ we have that

$$\overline{Y}_{ij} = \overline{Y}_{ji} = \sum_{k=1}^{n}\sum_{l=k+1}^{n} \frac{\binom{i-1+k-1}{i-1}\binom{j-i-1+l-k-1}{j-i-1}\binom{m-j+n-l}{m-j}}{\binom{m+n}{n}} Y_{kl} \qquad (10)$$

If we assume that $Y$ is a binary symmetric matrix (e.g. a native contact map), such summation need to be performed only for the pairs $1 \leq k < l \leq n$ such that $Y_{kl} \neq 0$.

The computation of each entry of the expectation matrix $\overline{Y}$ can be done with Equations 9 and 10. The calculation of Equation 10 need to be done for $m(m-1)/2$ entries of the $\overline{Y}$ matrix. The calculation of each $\overline{Y}_{ij}$ entry (with $i < j$) involves a summation of $n(n-1)/2$ terms. Then, the computational complexity of the $\overline{Y}$ calculation is bounded by $O(m^2n^2)$. This can be computationally intensive when $m$ is large and $Y$ is a dense matrix. On the other end, if the $Y$ matrix is sparse, the computational cost can be bounded by $O(m^2k)$, where $k$ is the number of non-zero entries of the $Y$ matrix. However, in practice, the calculation of the $\overline{Y}$ matrix can be performed quickly for both native contact/distance protein maps.

#### 4.3 Variance of the congruence coefficient over all possible alignments between two maps

By using the same combinatorial approach adopted for the expectation, we compute the variance $Var[r_c^\alpha(X,Y)]$ of the congruence coefficient over all possible alignments $\alpha$ between two symmetric matrices $X$ and $Y$. In this case the calculation involves the computation of the variance-covariance matrix:

$$Var[Y^\alpha] = E[Y^\alpha \otimes Y^\alpha] - E[Y^\alpha] \otimes E[Y^\alpha]$$

where $\otimes$ is the the Kronecker product. The variance-covariance matrix of the normalized $Y$ matrix, can be obtained by dividing the $Var[Y^\alpha]$ matrix with constant factor $tr(YY)$.

The variance of the congruence coefficient is, as usual, defined by:

$$Var[r_c^\alpha(X,Y)] \;\; = \;\; E[r_c^\alpha(X,Y)^2] - E[r_c^\alpha(X,Y)]^2$$

Then, in order to compute $Var[r_c^\alpha(X,Y)]$ we have to compute the expectation of the squared congruence coefficient $E[r_c^\alpha(X,Y)^2]$ over all possible alignments between $X$ and $Y$. Before to show how to solve $E[r_c^\alpha(X,Y)^2]$, we review the definition and some basic properties of the Kronecker product. In order to simplify the notation, we assume only square matrices, although the following properties and definitions hold for general matrices. Let $X \in \mathbb{R}^{m\times m}$ and $Y \in \mathbb{R}^{n\times n}$ two square matrices:

1. The Kronecker product $K = X \otimes Y \in \mathbb{R}^{mn\times mn}$ between $X$ and $Y$ is defined by

   $$K_{(ik)(jl)} = X_{ij}Y_{kl} \text{ with } 1 \leq i,j \leq m \text{ and } 1 \leq k,l \leq n,$$

   where the indices $(ik)$ (row indices in $X_{ij}Y_{kl}$) and $(jl)$ (column indices in $X_{ij}Y_{kl}$) of the $K$ matrix are defined by

   $$(ik) \rightarrow (i-1)*n+k \text{ and } (jl) \rightarrow (j-1)*n+l$$

2. $tr(XY \otimes XY) = tr(XY)tr(XY) = tr(XY)^2$
3. $(X \otimes X)(Y \otimes Y) = (XY) \otimes (XY)$

By using the Kronecked product notation we can easily transform $E[r_c^\alpha(X,Y)^2]$ into a more manageable form.

$$
\begin{aligned}
E[r_c^\alpha(X,Y)^2] &= \frac{\sum_{\alpha\in A(m,n)} r_c^\alpha(X,Y)^2}{|A(m,n)|} = \\
&= \frac{\sum_{\alpha\in A(m,n)} \left(\frac{tr(XY^\alpha)}{\sqrt{tr(XX)tr(YY)}}\right)^2}{|A(m,n)|} = \qquad \text{(since } tr(XX)tr(YY) \text{ is constant)} \\
&= \frac{E[tr(XY^\alpha)^2]}{tr(XX)tr(YY)} = \qquad \text{(by Kronecker's product property 2)} \\
&= \frac{E[tr((XY^\alpha)\otimes(XY^\alpha))]}{tr(XX)tr(YY)} = \qquad \text{(by Kronecker's product property 3)} \\
&= \frac{E[tr((X\otimes X)(Y^\alpha\otimes Y^\alpha))]}{tr(XX)tr(YY)} = \qquad \text{(since } tr \text{ is linear and } X\otimes X \text{ constant)} \\
&= \frac{tr((X\otimes X)E[Y^\alpha\otimes Y^\alpha])}{tr(XX)tr(YY)}
\end{aligned}
$$

where the *expectation of the square matrix* $E[Y^\alpha \otimes Y^\alpha]$, which we call $\tilde{Y}$, is defined by

$$
\tilde{Y} = E[Y^\alpha \otimes Y^\alpha] = \frac{\sum_{\alpha\in A(m,n)} Y^\alpha \otimes Y^\alpha}{|A(m,n)|} \in \mathbb{R}^{m^2\times m^2}
$$

and, entrywise, it is defined by

$$
\tilde{Y}_{(ik)(jl)} = \frac{\sum_{\alpha\in A(m,n)} Y_{ij}^\alpha Y_{kl}^\alpha}{|A(m,n)|} = \frac{\sum_{\alpha\in A(m,n),\alpha(i)\neq\perp,\alpha(j)\neq\perp,\alpha(k)\neq\perp,\alpha(l)\neq\perp} Y_{\alpha(i)\alpha(j)}Y_{\alpha(k)\alpha(l)}}{|A(m,n)|}
$$

The variance-covariance matrix for random matrices $Y^\alpha$

$$
Var[Y^\alpha] = E[Y^\alpha \otimes Y^\alpha] - E[Y^\alpha] \otimes E[Y^\alpha]
$$

can be seen as the matrix equivalent of the variance-covariance matrix for random vectors. In particular, $Var[Y^\alpha]$ contains the variance of all the elements $Y^\alpha_{ij}$, $Var[Y^\alpha_{ij}] = Cov[Y^\alpha_{ij}, Y^\alpha_{ij}]$ and the covariances $Cov[Y^\alpha_{ij}, Y^\alpha_{kl}]$ between every pair of matrix elements $Y^\alpha_{ij}$, $Y^\alpha_{kl}$, where $i \neq k$ or $j \neq l$. Note that, differently from the vector case, the variance values $Var[Y^\alpha_{ij}]$ are not all on the main diagonal of $Var[Y^\alpha]$, since the expected value $E[Y^\alpha_{ij} Y^\alpha_{ij}]$ in $\tilde{Y} = E[Y^\alpha \otimes Y^\alpha]$ is in position $\tilde{Y}_{(ii)(jj)}$, which is on the main diagonal only when $i = j$.

In order to compute $\tilde{Y}$, we need to consider four possible cases.

1. **One match**. Let $1 \leq i, j, k, l \leq m$ be four indices, such that $|\{i, j, k, l\}| = 1$. Without loss of generality, assume that

$$\{i, j, k, l\} = \{a\} \text{ with } 1 \leq a \leq m$$

   Now, for every $1 \leq b \leq n$ there is at least one alignment $\alpha \in A(m, n)$ such that $\alpha(a) = b$. The total number of such alignments can be computed by

$$\binom{s_1 + s_2}{s_1}\binom{e_1 + e_2}{e_1} = \binom{a - 1 + b - 1}{a - 1}\binom{m - a + n - b}{m - a}$$

   where

   - $s_1 = a - 1, s_2 = b - 1$ are the lengths of the sub-sequences before $a$ and $b$, respectively
   - $e_1 = m - a, e_2 = n - b$ are the lengths of the sub sequences starting immediately after $a$ and $b$ up to the end of the first and second sequence, respectively.

   Then if $1 \leq i, j, k, l \leq m$, $\{i, j, k, l\} = \{a\}$ we have that

$$\tilde{Y}_{(ij)(kl)} = \tilde{Y}_{(aa)(aa)} = \sum_{b=1}^{n} \frac{\binom{a-1+b-1}{a-1}\binom{m-a+n-b}{m-a}}{\binom{m+n}{n}} Y_{bb} Y_{bb} \qquad (11)$$

   As before, if we assume that $Y$ is a binary symmetric matrix representing protein contacts, we usually avoid do not consider contacts around the main diagonal. Then the main diagonal of $\tilde{Y}$ is zero, i.e. for every $1 \leq a \leq m$, $\tilde{Y}_{(aa)(aa)} = 0$ .

2. **Two matches**. Let $1 \leq i, j, k, l \leq m$ be four indices, such that $|\{i, j, k, l\}| = 2$. Without loss of generality, assume that

$$\{i,j,k,l\} = \{a,b\} \text{ and that } 1 \le a < b \le m$$

Now, for every pair of indexes $1 \le c < d \le n$ there is at least one alignment $\alpha \in A(m,n)$ such that $\alpha(a) = c, \alpha(b) = d$. The total number of such alignments can be computed by

$$\binom{s_1+s_2}{s_1}\binom{c_1+c_2}{c_1}\binom{e_1+e_2}{e_1} =$$

$$\binom{a-1+c-1}{a-1}\binom{b-a-1+d-c-1}{b-a-1}\binom{m-b+n-d}{m-b}$$

where (as we have seen before)

- $s_1 = a-1, s_2 = c-1$ are the lengths of the sub-sequences before $a$ and $c$, respectively
- $c_1 = b-a-1, c_2 = d-c-1$ are the lengths of the sub-sequences between $a,b$ and $c,d$ , respectively
- $e_1 = m-b, e_2 = n-d$ are the lengths of the sub sequences starting immediately after $b$ and $d$ up to the end of the first and second sequence, respectively.

Then, if $\{i,j,k,l\} = \{a,b\}, 1 \le a < b \le m$, we have that

$$\tilde{Y}_{(ik)(jl)} = \sum_{c=1}^{n}\sum_{d>c}^{n} \frac{\binom{a-1+c-1}{a-1}\binom{b-a-1+d-c-1}{b-a-1}\binom{m-b+n-d}{m-b}}{|A(m,n)|} Y_{f(i)f(j)}Y_{f(k)f(l)} \tag{12}$$

where

$$f(x) = \begin{cases} c & \text{if } x = a \\ d & \text{if } x = b \end{cases}$$

3. **Three matches**. If $m \ge 3$ and $n \ge 3$, let $1 \le i,j,k,l \le m$ be four indices, such that $|\{i,j,k,l\}| = 3$. Without loss of generality, assume that

$$\{i,j,k,l\} = \{a,b,c\} \text{ and that } 1 \le a < b < c \le m$$

As before, for every set of indexes $1 \leq d < e < f \leq n$ there is at least one alignment $\alpha \in A(m,n)$ such that $\alpha(a) = d, \alpha(b) = e, \alpha(c) = f$. The total number of such alignments can be computed by

$$\binom{s_1+s_2}{s_1}\binom{c_1+c_2}{c_1}\binom{c_1'+c_2'}{c_1'}\binom{e_1+e_2}{e_1} =$$

$$\binom{a-1+d-1}{a-1}\binom{b-a-1+e-d-1}{b-a-1}\binom{c-b-1+f-e-1}{c-b-1}\binom{m-c+n-f}{m-c}$$

Then, if $\{i,j,k,l\} = \{a,b,c\}, 1 \leq a < b < c \leq m$, we have that

$$\tilde{Y}_{(ik)(jl)} = \sum_{d=1}^{n}\sum_{e>d}^{n}\sum_{f>e}^{n} \frac{\binom{s_1+s_2}{s_1}\binom{c_1+c_2}{c_1}\binom{c_1'+c_2'}{c_1'}\binom{e_1+e_2}{e_1}}{|A(m,n)|} Y_{f(i)f(j)} Y_{f(k)f(l)} \tag{13}$$

where

$$f(x) = \begin{cases} d & \text{if } x = a \\ e & \text{if } x = b \\ f & \text{if } x = c \end{cases}$$

4. **Four matches**. If $m \geq 4$ and $n \geq 4$, let $1 \leq i,j,k,l \leq m$ be four indices, such that $|\{i,j,k,l\}| = 4$. Without loss of generality, assume that

$$\{i,j,k,l\} = \{a,b,c,d\} \text{ and that } 1 \leq a < b < c < d \leq m$$

As before, for every set of indexes $1 \leq e < f < g < h \leq n$ there is at least one alignment $\alpha \in A(m,n)$ such that $\alpha(a) = e, \alpha(b) = f, \alpha(c) = g, \alpha(d) = h$. The total number of such alignments can be computed by

$$\binom{s_1+s_2}{s_1}\binom{c_1+c_2}{c_1}\binom{c_1'+c_2'}{c_1'}\binom{c_1''+c_2''}{c_1''}\binom{e_1+e_2}{e_1} =$$

$$\binom{a-1+e-1}{a-1}\binom{b-a-1+f-e-1}{b-a-1}\binom{c-b-1+g-f-1}{c-b-1}\binom{m-d+n-h}{m-d}$$

Then, if $\{i, j, k, l\} = \{a, b, c, d\}, 1 \leq a < b < c < d \leq m$, we have that

$$\tilde{Y}_{(ik)(jl)} = \sum_{e=1}^{n} \sum_{f>e}^{n} \sum_{g>f}^{n} \sum_{h>g}^{n} \frac{\binom{s_1+s_2}{s_1}\binom{c_1+c_2}{c_1}\binom{c_1'+c_2'}{c_1'}\binom{c_1''+c_2''}{c_1''}\binom{e_1+e_2}{e_1}}{|A(m,n)|} Y_{f(i)f(j)} Y_{f(k)f(l)} \tag{14}$$

where

$$f(x) = \begin{cases} e & \text{if } x = a \\ f & \text{if } x = b \\ g & \text{if } x = c \\ h & \text{if } x = d \end{cases}$$

The calculation of each entry of the average matrix $\tilde{Y}$ can be done with Equations 11) 12, 13 and 14. The computation of the *four matches* in Equations 14 is computationally challenging for large target sizes $m$. In particular, in a $m^2 \times m^2$ symmetric matrix $\tilde{Y}$ there are $\binom{m}{4} \sim m^4$ entries $\tilde{Y}_{(ik)(jl)}$ such that $|\{i, j, k, l\}| = 4$ (i.e. the four matches case). The summation in Equation 14 generally involves $\binom{n}{4} \sim n^4$ terms, then the computational cost for generating the $\tilde{Y}$ matrix is bounded by $O(m^4 n^4)$. If $Y$ is sparse, such cost can be lowered down to only $O(m^4 k^2)$, where $k$ is the number of non-zero entries in $Y$. Thus, even when $Y$ is a native contact map, the computation of the $\tilde{Y}$ matrix is still extremely costly for large $m$.

The $\tilde{Y}$ matrix is also not easily approximable, both in terms of computational time and memory usage. We tried some approaches for approximating $\tilde{Y}$, all of which give poor estimates of the $\tilde{Y}$ matrix. Some of the tried approaches include: i) computation of upper/lower bounds by using Jensen's inequality (extremely inaccurate); ii) approximation by Taylor expansion (doesn't work since the trace is a linear function, thus it is not actually approximated by the Taylor series); iii) estimation of the (four matches) entries in Equation 14 from the (three matches) entries computed in Equation 13 (extremely inaccurate). As a further drawback, for real protein representation maps, the $\tilde{Y}$ matrices are typically huge (on the order of few hundred Mega byte) in comparison to the $\overline{Y}$ matrices (few Mega bytes). By rounding to zero all values lower than some small cutoff we can reduce the memory occupation. However, while a cutoff equal to $10^{-10}$ provides a small reduction in size but doesn't affect the variance calculation, a slightly larger cutoff of $10^{-6}$ provides a much larger reduction but totally affects the variance value, which very often becomes negative.

A different approach for estimating the variance of the $r_c^{\alpha}(X, Y)$ coefficient is through adhoc random alignment sampling (see Section 7.3). We remark that, the alignment sampling has been used to compute directly the variance of the $r_c^\alpha(X, Y)$ coefficients. Alignment sampling for the estimation of the $\widetilde{Y}$ matrix is quite costly, since it involves the computation of the Kronecked product for the aligned $Y^\alpha$ matrices. In particular, at least when $Y$ is a sparse binary matrix, the estimation of $\widetilde{Y}$ through sampling is slower than the exact computation, even when the sampled size is small (e.g. on the order of few thousands alignments).

In conclusion, a much deeper investigation need to be done to exploit possible analytical techniques that can reasonably approximate the variance calculation.

### 5 Expectation and variance over permutations

The expectation and variance over all permutations can be defined in terms of closed formulas that essentially depend on the sum of the elements in the two matrices. The expectation formula has a simple characterization, while the variance is more complicated and involves several different terms.

**Definition 5.1.** *Any surjective and injective function $\sigma : \{1, .., n\} \to \{1, .., n\}$ is a* **permutation**.

For a given $n$, we denote with $P(n)$ the set of all permutations $\sigma : \{1, .., n\} \to \{1, .., n\}$. The cardinality of the set $P(n)$ is $n!$.

Given two symmetric matrices $X, Y \in \mathbb{R}^{n\times n}$ and a permutation $\sigma \in P(n)$, we can defined the congruence coefficient with respect to permutation $\sigma$ by

$$r^\sigma(X, Y) = r_c(X, Y^\sigma) = \frac{tr(XY^\sigma)}{\sqrt{tr(XX)tr(Y^\sigma Y^\sigma)}} = \frac{tr(XY^\sigma)}{\sqrt{tr(XX)tr(YY)}} = \frac{tr(X^{\sigma^{-1}}Y)}{\sqrt{tr(XX)tr(YY)}}$$

where $Y^\sigma$ is defined in Equation 6. Note that, as it happens for alignments, the normalization factor of the congruence coefficient is invariant with respect to permutations. Furthermore, note that, if $\sigma_1, \sigma_2 \in P(n)$ are two permutations, then $\sigma_2 \circ \sigma_1 \in P(n)$, $\sigma_1^{-1}, \sigma_2^{-1} \in P(n)$ and $\sigma_2^{-1} \circ \sigma_1 \in P(n)$. Then,

$$r_c(X^{\sigma_1}, Y^{\sigma_2}) = r_c(X, Y^{\sigma_2^{-1}\circ\sigma_1}) = r_c^{\sigma_2^{-1}\circ\sigma_1}(X, Y)$$

so we just need to consider permutations for only one of the two maps.

When $X$ and $Y$ have different sizes, we can add zero rows and columns to $X$ and $Y$ in order to make them have the same sizes for calculation. The expectation and variance of the congruence coefficient are not affected by the added zero rows/columns, except for a multiplication factor that depends on the size of the recoded matrices. In detail, let $X \in \mathbb{R}^{m\times m}$, $Y \in \mathbb{R}^{n\times n}$ be two symmetric matrices and let $\sigma \in P(N)$, with $N \geq \max\{m, n\}$, be a permutation. We can recode $X$ (resp. $Y$) into a matrix $X' \in \mathbb{R}^{N\times N}$ by

$$X'_{ij} = \begin{cases} X_{ij} & \text{if } 1 \leq i, j \leq m \\ 0 & \text{if } m < i \leq N \text{ or } m < j \leq N \end{cases}$$

We can then define $r_c^\sigma(X, Y)$ by

$$r_c^\sigma(X, Y) = r_c^\sigma(X', Y') = \frac{tr(X'Y'^\sigma)}{\sqrt{tr(X'X')tr(Y'Y')}} = \frac{tr(X'Y'^\sigma)}{\sqrt{tr(XX)tr(YY)}}$$

We can show that, if we assume that the main diagonals of $X \in \mathbb{R}^{m\times m}$ and $Y \in \mathbb{R}^{n\times n}$ are zero, the expected value and the variance of the congruence coefficient with respect to all permutations $\sigma \in P(N)$, with $N \geq \max\{m, n\}$, can be defined in terms of the following quantities, that depend on $X$ and $Y$ but not on $N$:

- $\Sigma(X) = \Sigma(X')$: sum of all elements in $X$
- $\Sigma(X^2) = \Sigma(X'^2) = tr(XX)$: sum of all squared elements in $X$
- $\Sigma(XX) = \Sigma(X'X')$: sum of all elements in the matrix product $XX$

Then, if the main diagonals of $X \in \mathbb{R}^{m\times m}$ and $Y \in \mathbb{R}^{n\times n}$ are zero and $\sigma \in P(N)$, with $N \geq \max\{m, n\}$:

$$E[r_c^\sigma(X, Y)] = \frac{\Sigma(X)\Sigma(Y)}{N(N-1)\sqrt{\Sigma(X^2)\Sigma(Y^2)}} \tag{15}$$

and

$$\begin{aligned} Var[r_c^\sigma(X, Y)] \quad =& \frac{2}{N(N-1)} + \frac{4\left[\Sigma(XX) - \Sigma(X^2)\right]\left[\Sigma(YY) - \Sigma(Y^2)\right]}{N(N-1)(N-2)\Sigma(X^2)\Sigma(Y^2)} + \\ &+ \frac{\left[\Sigma(X)^2 - 4\Sigma(XX) + 2\Sigma(X^2)\right]\left[\Sigma(Y)^2 - 4\Sigma(YY) + 2\Sigma(Y^2)\right]}{N(N-1)(N-2)(N-3)\Sigma(X^2)\Sigma(Y^2)} + \\ &- \frac{\Sigma(X)^2\Sigma(Y)^2}{N^2(N-1)^2\Sigma(X^2)\Sigma(Y^2)} \end{aligned} \tag{16}$$

If $N = m + n$, it is easy to see that the set of permutations $P(m + n)$ strictly contains the set of all possible alignments between $X$ and $Y$.

The expectation and variance formulas for general symmetric matrices can be computed equivalently (the variance formula is much larger). Also for the general case, the expectation and variance are not affected by the zero rows/columns added in $X'$ and $Y'$, since their formulas are defined in terms of quantities (summations) that depend on $X$, $Y$, and of a multiplication factor that depends on $N$.

In the following sections we show how to obtain $E[r_c^\sigma(X,Y)]$ and $Var[r_c^\sigma(X,Y)]$ when $X, Y$ have zero main diagonals and also for the general case where the main diagonals are not zero.

#### 5.1 Expectation of the congruence coefficient over all possible permutations

Let $X \in \mathbb{R}^{n\times n}, Y \in \mathbb{R}^{n\times n}$ be two symmetric matrices. The expectation of the congruence coefficient $r_c^\sigma(X,Y)$ over all $n!$ permutations $\sigma \in P(n)$ is defined by

$$
\begin{aligned}
E[r_c^\sigma(X,Y)] &= \frac{\sum\limits_{\sigma\in P(n)} r_c^\sigma(X,Y)}{n!} = \\
&= \frac{\sum\limits_{\sigma\in P(n)} tr(XY^\sigma)}{n!\sqrt{tr(XX)tr(YY)}} \\
&= \frac{E[tr(XY^\sigma)]}{\sqrt{tr(XX)tr(YY)}}
\end{aligned}
$$

We need to solve the expectation of the trace of the product between $X$ and $Y$ over all permutations $\sigma$.

$$
\begin{aligned}
E[tr(X,Y^\sigma)] &= \frac{\displaystyle\sum_{\sigma\in P(n)}\sum_{i=1}^{n}\sum_{j=1}^{n} X_{ij}Y_{\sigma(i)\sigma(j)}}{n!} \\
&= \frac{\displaystyle\sum_{\sigma\in P(n)}\sum_{i=1}^{n} X_{ii}Y_{\sigma(i)\sigma(i)}}{n!} + \frac{\displaystyle\sum_{\sigma\in P(n)}\sum_{i=1}^{n}\sum_{j=1,j\neq i}^{n} X_{ij}Y_{\sigma(i)\sigma(j)}}{n!} = \\
&= \frac{\displaystyle\sum_{i=1}^{n} X_{ii}\left(\sum_{\sigma\in P(n)} Y_{\sigma(i)\sigma(i)}\right)}{n!} + \frac{\displaystyle\sum_{i=1}^{n}\sum_{j=1,j\neq i}^{n} X_{ij}\left(\sum_{\sigma\in P(n)} Y_{\sigma(i)\sigma(j)}\right)}{n!} = \\
&= \frac{\displaystyle\sum_{i=1}^{n} X_{ii}\left(\sum_{\sigma\in P(n)} Y_{\sigma(i)\sigma(i)}\right)}{n!} + \frac{\displaystyle\sum_{i=1}^{n}\sum_{j=1,j\neq i}^{n} X_{ij}\left(\sum_{\sigma\in P(n)} Y_{\sigma(i)\sigma(j)}\right)}{n!}
\end{aligned}
$$

We have two possible cases:

1. **Diagonal elements**. If $\sigma(i) = k$, there are $(n-1)!$ permutations $\sigma' \in P(n)$ such that $\sigma'(i) = k$. Then

$$\sum_{\sigma\in P(n)} Y_{\sigma(i)\sigma(i)} = (n-1)!\sum_{k=1}^{n} Y_{kk} = (n-1)!tr(Y)$$

from which we obtain:

$$\frac{\displaystyle\sum_{i=1}^{n} X_{ii}\left(\sum_{\sigma\in P(n)} Y_{\sigma(i)\sigma(i)}\right)}{n!} = \frac{(n-1)!tr(X)tr(Y)}{n!} = \frac{tr(X)tr(Y)}{n}$$

2. **Non-diagonal elements**. If $\sigma(i) = k$ and $\sigma(j) = l$, there are $(n-2)!$ permutations $\sigma' \in P(n)$ such that $\sigma'(i) = k$ and $\sigma'(j) = l$. Then

$$\sum_{\sigma\in P(n)} Y_{\sigma(i)\sigma(j)} = (n-2)!\sum_{k=1}^{n}\sum_{k\neq l=1}^{n} Y_{kl} = (n-2)!(\Sigma(Y) - tr(Y))$$

from which we obtain:

$$\frac{\sum_{i=1}^{n}\sum_{i\neq j=1}^{n} X_{ij}\left(\sum_{\sigma\in P(n)} Y_{\sigma(i)\sigma(j)}\right)}{n!} = \frac{(n-2)!(\Sigma(X)-tr(X))(\Sigma(Y)-tr(Y))}{n!} =$$

$$= \frac{(\Sigma(X)-tr(X))(\Sigma(Y)-tr(Y))}{n(n-1)}$$

Putting altogether:

$$E[tr(XY^{\sigma})] = \frac{tr(X)tr(Y)}{n} + \frac{(\Sigma(X)-tr(X))(\Sigma(Y)-tr(Y))}{n(n-1)}$$

Note that, the expected value of the trace of the product over all permutations is greatly simplified in comparison to the expected value over all alignments. The reason is that (compare the *Diagonal elements* case in this Section with case 1 in Section 4.2), if we fix a permutation for an index $i$, say $\sigma(i) = k$, then we have that the number of permutations $\sigma' \in P(n)$, such that $\sigma'(i) = k$, is equal to $(n-1)!$, independently of $i$ and $k$. On the contrary, if we fix a matching for an index $i$, say $\alpha(i) = k$, then the number of alignments $\alpha' \in A(m,n)$, such that $\alpha'(i) = k$, is equal to $\binom{i-1+k-1}{i-1}\binom{m-i+n-k}{m-i}$, which highly depends on the indices $i$ and $k$. For example, if $i = k = 1$ the number of possible alignments compatible with such match are $\binom{m-1+n-1}{m-1}$, while if $i = 1$ and $k = n$, the number of possible alignments is 1.

In order to simplify the computation (especially for the variance), we can define the expectation only in terms of sum of all the elements in the matrices $X$, $Y$ and $\mathring{X}$ and $\mathring{Y}$, where $\mathring{X}$ is a matrix equal to $X$ except for the main diagonal, which is zero:

$$\mathring{X}_{ij} = \begin{cases} X_{ij} & i \neq j \\ 0 & i = j \end{cases}$$

We also use the following trivial equivalences to simply formulas:

- $\Sigma(\mathring{X}) = \Sigma(X) - tr(X)$
- $tr(XX) = \Sigma(X^2)$

We can then define the expectation of the congruence coefficient over all permutations of the $Y$ matrix as

$$E[r_c^{\sigma}(X,Y)] = \frac{tr(X)tr(Y)}{n\sqrt{\Sigma(X^2)\Sigma(Y^2)}} + \frac{\Sigma(\mathring{X})\Sigma(\mathring{Y})}{n(n-1)\sqrt{\Sigma(X^2)\Sigma(Y^2)}} \tag{17}$$

If we assume that the diagonals of $X$ and $Y$ are zero (and then $X = \mathring{X}$, $Y = \mathring{Y}$) we get the shorter formula:

$$E[r_c^\sigma(X,Y)] = \frac{\Sigma(X)\Sigma(Y)}{n(n-1)\sqrt{\Sigma(X^2)\Sigma(Y^2)}} \tag{18}$$

#### 5.2 Variance of the congruence coefficient over all possible permutations

Let $X \in \mathbb{R}^{n\times n}, Y \in \mathbb{R}^{n\times n}$ be two symmetric matrices. The variance of the congruence coefficient $r_c^\sigma(X,Y)$ over all $n!$ permutations $\sigma \in P(n)$ is defined by

$$Var[r_c^\sigma(X,Y)] = \frac{E[tr(XY^\sigma)^2] - E[tr(XY^\sigma)]^2}{tr(XX)tr(YY)}$$

We need to solve the expectation of the squared trace of the product between $X$ and $Y$ over all permutations $\sigma$.

$$\begin{aligned} E[tr(XY^\sigma)^2] &= \frac{1}{n!}\sum_{\sigma\in P(n)}\left(\sum_{i=1}^{n}\sum_{j=1}^{n} X_{ij}Y_{\sigma(i)\sigma(j)}\right)^2 \\ &= \frac{1}{n!}\sum_{\sigma\in P(n)}\sum_{i=1}^{n}\sum_{j=1}^{n}\sum_{k=1}^{n}\sum_{l=1}^{n} X_{ij}X_{kl}Y_{\sigma(i)\sigma(j)}Y_{\sigma(k)\sigma(l)} \\ &= \frac{S_1+S_2+S_3+S_4}{n!} \end{aligned}$$

where the four quantities $S_1, S_2, S_3, S_4$ are defined as follows:

1. $S_1 = \sum_{i,j,k,l\in\{1,..,n\},|\{\mathbf{i},\mathbf{j},\mathbf{k},\mathbf{l}\}|=\mathbf{1}}^{n} X_{ij}X_{kl}\sum_{\sigma\in P(n)} Y_{\sigma(i)\sigma(j)}Y_{\sigma(k)\sigma(l)}$

2. $S_2 = \sum_{i,j,k,l\in\{1,..,n\},|\{\mathbf{i},\mathbf{j},\mathbf{k},\mathbf{l}\}|=\mathbf{2}}^{n} X_{ij}X_{kl}\sum_{\sigma\in P(n)} Y_{\sigma(i)\sigma(j)}Y_{\sigma(k)\sigma(l)}$

3. $S_3 = \sum_{i,j,k,l\in\{1,..,n\},|\{\mathbf{i},\mathbf{j},\mathbf{k},\mathbf{l}\}|=\mathbf{3}}^{n} X_{ij}X_{kl}\sum_{\sigma\in P(n)} Y_{\sigma(i)\sigma(j)}Y_{\sigma(k)\sigma(l)}$

4. $S_4 = \sum_{i,j,k,l \in \{1,..,n\}, |\{\mathbf{i,j,k,l}\}|=\mathbf{4}}^{n} X_{ij} X_{kl} \sum_{\sigma \in P(n)} Y_{\sigma(i)\sigma(j)} Y_{\sigma(k)\sigma(l)}$

Then we need to consider four distinct cases, where the first term, $S_1$, has a simple closed form, while the remaining terms are more complicated and involve complicated summations over several indices. Also in this case, we give closed formulas in terms of the matrices $X, X^2, XX, \mathring{X}, \mathring{X}^2, \mathring{X}\mathring{X}, \mathring{X}X$ (resp. for $Y$), where

- $\mathring{X}$ corresponds to the $X$ matrix with zero diagonal
- $X^2$ corresponds to the matrix $X$ in which every element has been squared
- $XX$ is the matrix product of $X$ with itself
- $X\mathring{X}$ is the matrix product between $X$ and $\mathring{X}$

The formulas can be then expressed in terms of matrix traces and sum of all elements in a matrix:

- $tr(X)$: trace of $X$
- $\Sigma(X)$: sum of all elements in $X$

The complete formula for the expectation of the squared trace is:

$$E[tr(XY^\sigma)^2] = \frac{(n-1)!S_{11} + (n-2)!(S_{21} + 2S_{22} + 4S_{23}) + (n-3)!\,(2S_{31} + 4S_{32}) + (n-4)!S_{41}}{n!}$$

where:

- $S_{11} = tr(X^2)tr(Y^2)$
- $S_{21} = \left[tr^2(X) - tr(X^2)\right]\left[tr^2(Y) - tr(Y^2)\right]$
- $S_{22} = \Sigma(\mathring{X}^2)\Sigma(\mathring{Y}^2)$
- $S_{23} = \left[\Sigma(\mathring{X}X) - \Sigma(\mathring{X}\mathring{X})\right]\left[\Sigma(\mathring{Y}Y) - \Sigma(\mathring{Y}\mathring{Y})\right]$
- $S_{31} = \left[2\Sigma(\mathring{X}\mathring{X}) + \Sigma(X)\Sigma(\mathring{X}) - 2\Sigma(X\mathring{X}) - \Sigma(\mathring{X})^2\right]\left[2\Sigma(\mathring{Y}\mathring{Y}) + \Sigma(Y)\Sigma(\mathring{Y}) - 2\Sigma(Y\mathring{Y}) - \Sigma(\mathring{Y})^2\right]$
- $S_{32} = \left[\Sigma(\mathring{X}\mathring{X}) - \Sigma(\mathring{X}^2)\right]\left[\Sigma(\mathring{Y}\mathring{Y}) - \Sigma(\mathring{Y}^2)\right]$

- $S_{41} = \left[\Sigma^2(\mathring{X}) - 4\Sigma(\mathring{X}\mathring{X}) + 2\Sigma(\mathring{X}^2)\right]\left[\Sigma^2(\mathring{Y}) - 4\Sigma(\mathring{Y}\mathring{Y}) + 2\Sigma(\mathring{Y}^2)\right]$

If the matrices $X$ and $Y$ have zero diagonal, then $tr(X) = tr(Y) = tr(X^2) = tr(Y^2) = 0$, $X = \mathring{X}$ and $Y = \mathring{Y}$ which implies that

$$S_{11} = S_{21} = S_{23} = S_{31} = 0$$

and then

$$E[tr(XY^\sigma)^2] = \frac{2(n-2)!S_{22} + 4(n-3)!S_{32} + (n-4)!S_{41}}{n!}$$

In the following we show directly the result of the summations for the $S_1, S_2, S_3, S_4$, terms. The calculation details can be found in Appendix A.

##### 5.2.1 Term $S_1$: summation over a unique index

The term $S_1$ is given by the summation over a unique index $i \in \{1, .., n\}$.

$$\begin{aligned} S_1 &= \sum_{i,j,k,l \in \{1,..,n\}, |\{\mathbf{i},\mathbf{j},\mathbf{k},\mathbf{l}\}|=\mathbf{1}}^{n} X_{ij}X_{kl} \sum_{\sigma \in P(n)} Y_{\sigma(i)\sigma(j)}Y_{\sigma(k)\sigma(l)} = \\ &= \sum_{i=1}^{n} X_{ii}X_{ii} \sum_{\sigma \in P(n)} Y_{\sigma(i)\sigma(i)}Y_{\sigma(i)\sigma(i)} = \\ &= \sum_{i=1}^{n} X_{ii}^2 \sum_{\sigma \in P(n)} Y_{\sigma(i)\sigma(i)}^2 = \quad \text{(by 1 in Appendix A)} \\ &= (n-1)!tr(X^2)tr(Y^2) \end{aligned}$$

If we put

$$S_{11} = tr(X^2)tr(Y^2)$$

we have that

$$S_1 = (n-1)!S_{11}$$

If $X$ and $Y$ have zero diagonals, then the term $S1$ is equal to zero.

##### 5.2.2 Term $S_2$: summation over two distinct indexes

The term $S_2$ is given by the summation over two distinct indices $i, j \in \{1, ..n\}$ with $i \neq j$. The number of possible combination of two distinct indices is given by:

$$
\begin{aligned}
S_2 &= \sum_{i,j,k,l \in \{1,..,n\}, |\{\mathbf{i},\mathbf{j},\mathbf{k},\mathbf{l}\}|=\mathbf{2}}^{n} X_{ij} X_{kl} \sum_{\sigma \in P(n)} Y_{\sigma(i)\sigma(j)} Y_{\sigma(k)\sigma(l)} = \\
&= \sum_{i=1}^{n} \sum_{i \neq j=1}^{n} \Bigg[ X_{ii} X_{jj} \sum_{\sigma \in P(n)} Y_{\sigma(i)\sigma(i)} Y_{\sigma(j)\sigma(j)} + \\
&\quad + X_{ij} X_{ij} \textstyle\sum_{\sigma \in P(n)} Y_{\sigma(i)\sigma(j)} Y_{\sigma(i)\sigma(j)} + X_{ij} X_{ji} \sum_{\sigma \in P(n)} Y_{\sigma(i)\sigma(j)} Y_{\sigma(j)\sigma(i)} + \\
&\quad + X_{ii} X_{ij} \textstyle\sum_{\sigma \in P(n)} Y_{\sigma(i)\sigma(i)} Y_{\sigma(i)\sigma(j)} + X_{ii} X_{ji} \sum_{\sigma \in P(n)} Y_{\sigma(i)\sigma(i)} Y_{\sigma(j)\sigma(i)} + \\
&\quad + X_{ij} X_{ii} \textstyle\sum_{\sigma \in P(n)} Y_{\sigma(i)\sigma(j)} Y_{\sigma(i)\sigma(i)} + X_{ji} X_{ii} \sum_{\sigma \in P(n)} Y_{\sigma(j)\sigma(i)} Y_{\sigma(i)\sigma(i)} \Bigg]
\end{aligned}
$$

Note that, since X (resp. Y) is symmetric, $X_{ij} = X_{ji}$ (resp. $Y_{\sigma(i)\sigma(j)} = Y_{\sigma(j)\sigma(i)}$) and, since the product is commutative,

$$X_{ii} X_{ij} = X_{ii} X_{ji} = X_{ij} X_{ii} = X_{ji} X_{ii} \text{ (resp. for } Y_{\sigma(i)\sigma(i)} Y_{\sigma(i)\sigma(j)} \text{)}$$

Then the $S_2$ summation reduces to:

$$S_2 = \sum_{i=1}^{n} \sum_{i \neq j=1}^{n} \left[ X_{ii} X_{jj} \sum_{\sigma \in P(n)} Y_{\sigma(i)\sigma(i)} Y_{\sigma(j)\sigma(j)} + 2X_{ij}^2 \sum_{\sigma \in P(n)} Y_{\sigma(i)\sigma(j)}^2 + 4X_{ii} X_{ij} \sum_{\sigma \in P(n)} Y_{\sigma(i)\sigma(i)} Y_{\sigma(i)\sigma(j)} \right]$$

where (see 5 in Appendix A)

$$\sum_{i=1}^{n} \sum_{i \neq j=1}^{n} X_{ii} X_{jj} \sum_{\sigma \in P(n)} Y_{\sigma(i)\sigma(i)} Y_{\sigma(j)\sigma(j)} = (n-2)! \left[ tr^2(X) - tr(X^2) \right] \left[ tr^2(Y) - tr(Y^2) \right]$$

and (see 8 in Appendix A)

$$
\begin{aligned}
\sum_{i=1}^{n} \sum_{i \neq j=1}^{n} 2X_{ij}^2 \sum_{\sigma \in P(n)} Y_{\sigma(i)\sigma(j)}^2 &= 2(n-2)! \left[ \Sigma(X^2) - tr(X^2) \right] \left[ \Sigma(Y^2) - tr(Y^2) \right] = \\
&= 2(n-2)! \Sigma(\mathring{X}^2) \Sigma(\mathring{Y}^2)
\end{aligned}
$$

and (see 6 in Appendix A)

$$\sum_{i=1}^{n}\sum_{i\neq j=1}^{n} 4X_{ii}X_{ij}\sum_{\sigma\in P(n)} Y_{\sigma(i)\sigma(i)}Y_{\sigma(i)\sigma(j)} = 4(n-2)!\left[\Sigma(\mathring{X}X)-\Sigma(\mathring{X}\mathring{X})\right]\left[\Sigma(\mathring{Y}Y)-\Sigma(\mathring{Y}\mathring{Y})\right]$$

If we put

- $S_{21} = \left[tr^2(X) - tr(X^2)\right]\left[tr^2(Y) - tr(Y^2)\right]$
- $S_{22} = \Sigma(\mathring{X}^2)\Sigma(\mathring{Y}^2)$
- $S_{23} = \left[\Sigma(\mathring{X}X)-\Sigma(\mathring{X}\mathring{X})\right]\left[\Sigma(\mathring{Y}Y)-\Sigma(\mathring{Y}\mathring{Y})\right]$

we have

$$S_2 = (n-2)!\,(S_{21} + 2S_{22} + 4S_{23})$$

If the main diagonals of $X$ and $Y$ are zero, and then $X = \mathring{X}$ (resp. $Y = \mathring{X}$), the $S_{21}$ and $S_{23}$ terms are zero, thus the $S_2$ term reduces to

$$S_2 = 2(n-2)!\Sigma(X^2)\Sigma(Y^2)$$

###### 5.2.3 Term $S_3$: summation over three distinct indexes

The term $S_3$ is given by the summation over three distinct indices $i, j, k \in \{1, ..n\}$ with $i \neq j \neq k$. The number of possible combination of three distinct indices is given by:

$$\begin{aligned}
S_3 &= \sum_{i,j,k,l\in\{1,..,n\},|\{\mathbf{i},\mathbf{j},\mathbf{k},\mathbf{l}\}|=\mathbf{3}}^{n} X_{ij}X_{kl}\sum_{\sigma\in P(n)} Y_{\sigma(i)\sigma(j)}Y_{\sigma(k)\sigma(l)} = \\
&= \sum_{i=1}^{n}\sum_{i\neq j=1}^{n}\sum_{i,j\neq k=1}^{n}\left[X_{ii}X_{jk}\sum_{\sigma\in P(n)} Y_{\sigma(i)\sigma(i)}Y_{\sigma(j)\sigma(k)} + X_{jk}X_{ii}\sum_{\sigma\in P(n)} Y_{\sigma(j)\sigma(k)}Y_{\sigma(i)\sigma(i)} + \right.\\
&\quad + X_{ij}X_{ik}\sum_{\sigma\in P(n)} Y_{\sigma(i)\sigma(j)}Y_{\sigma(i)\sigma(k)} + X_{ij}X_{ki}\sum_{\sigma\in P(n)} Y_{\sigma(i)\sigma(j)}Y_{\sigma(k)\sigma(i)} + \\
&\quad \left. + X_{ji}X_{ik}\sum_{\sigma\in P(n)} Y_{\sigma(j)\sigma(i)}Y_{\sigma(i)\sigma(k)} + X_{ji}X_{ki}\sum_{\sigma\in P(n)} Y_{\sigma(j)\sigma(i)}Y_{\sigma(k)\sigma(i)}\right]
\end{aligned}$$

As for $S_2$, since $X, Y$ are symmetric and the product is commutative, the above expression simplifies into

$$S_3 = \sum_{i=1}^{n} \sum_{i \neq j=1}^{n} \sum_{i,j \neq k=1}^{n} \left[ 2X_{ii}X_{jk} \sum_{\sigma \in P(n)} Y_{\sigma(i)\sigma(i)}Y_{\sigma(j)\sigma(k)} + 4X_{ij}X_{ik} \sum_{\sigma \in P(n)} Y_{\sigma(i)\sigma(j)}Y_{\sigma(i)\sigma(k)} \right]$$

where (see 18 in Appendix A)

$$\sum_{i=1}^{n} \sum_{i \neq j=1}^{n} \sum_{i,j \neq k=1}^{n} 2X_{ii}X_{jk} \sum_{\sigma \in P(n)} Y_{\sigma(i)\sigma(i)}Y_{\sigma(j)\sigma(k)} =$$

$$2(n-3)! \left[ 2\Sigma(\mathring{X}\mathring{X}) + \Sigma(X)\Sigma(\mathring{X}) - 2\Sigma(X\mathring{X}) - \Sigma(\mathring{X})^2 \right] \left[ 2\Sigma(\mathring{Y}\mathring{Y}) + \Sigma(Y)\Sigma(\mathring{Y}) - 2\Sigma(Y\mathring{Y}) - \Sigma(\mathring{Y})^2 \right]$$

and (see 15 in Appendix A)

$$\sum_{i=1}^{n} \sum_{i \neq j=1}^{n} \sum_{i,j \neq k=1}^{n} 4X_{ij}X_{ik} \sum_{\sigma \in P(n)} Y_{\sigma(i)\sigma(j)}Y_{\sigma(i)\sigma(k)} = 4(n-3)! \left[ \Sigma(\mathring{X}\mathring{X}) - \Sigma(\mathring{X}^2) \right] \left[ \Sigma(\mathring{Y}\mathring{Y}) - \Sigma(\mathring{Y}^2) \right]$$

If we put

- $S_{31} = \left[ 2\Sigma(\mathring{X}\mathring{X}) + \Sigma(X)\Sigma(\mathring{X}) - 2\Sigma(X\mathring{X}) - \Sigma(\mathring{X})^2 \right] \left[ 2\Sigma(\mathring{Y}\mathring{Y}) + \Sigma(Y)\Sigma(\mathring{Y}) - 2\Sigma(Y\mathring{Y}) - \Sigma(\mathring{Y})^2 \right]$
- $S_{32} = \left[ \Sigma(\mathring{X}\mathring{X}) - \Sigma(\mathring{X}^2) \right] \left[ \Sigma(\mathring{Y}\mathring{Y}) - \Sigma(\mathring{Y}^2) \right]$

we have

$$S_3 = (n-3)! \left( 2S_{31} + 4S_{32} \right)$$

If the main diagonals of $X$ and $Y$ are zero, and then $X = \mathring{X}$ (resp. $Y = \mathring{X}$), the $S_{31}$ term is zero, thus the $S_3$ term reduces to

$$S_3 = 4(n-3)! \left[ \Sigma(XX) - \Sigma(X^2) \right] \left[ \Sigma(YY) - \Sigma(Y^2) \right]$$

###### 5.2.4 Term $S_4$: summation over four distinct indexes

The term $S_4$ is given by the summation over four distinct indices $i, j, k, l \in \{1, ..n\}$ with $i \neq j \neq k \neq l$ (see 22 in Appendix A).

$$\begin{aligned}
S_4 &= \sum_{i,j,k,l \in \{1,..,n\}, |\{\mathbf{i},\mathbf{j},\mathbf{k},\mathbf{l}\}|=\mathbf{4}}^{n} X_{ij} X_{kl} \sum_{\sigma \in P(n)} Y_{\sigma(i)\sigma(j)} Y_{\sigma(k)\sigma(l)} = \\
&= \sum_{i=1}^{n} \sum_{i \neq j=1}^{n} \sum_{i,j \neq k=1}^{n} \sum_{i,j,k \neq l=1}^{n} X_{ij} X_{kl} \sum_{\sigma \in P(n)} Y_{\sigma(i)\sigma(j)} Y_{\sigma(k)\sigma(l)} = \\
&= (n-4)! \left[ \Sigma^2(\mathring{X}) - 4\Sigma(\mathring{X}\mathring{X}) + 2\Sigma(\mathring{X}^2) \right] \left[ \Sigma^2(\mathring{Y}) - 4\Sigma(\mathring{Y}\mathring{Y}) + 2\Sigma(\mathring{Y}^2) \right]
\end{aligned}$$

If he main diagonals of $X$ and $Y$ are zero, and then $X = \mathring{X}$ (resp. $Y = \mathring{X}$), the $S_4$ term becomes

$$S_4 = (n-4)! \left[ \Sigma^2(X) - 4\Sigma(XX) + 2\Sigma(X^2) \right] \left[ \Sigma^2(Y) - 4\Sigma(YY) + 2\Sigma(Y^2) \right]$$

### 6 Statistical significance of the congruence coefficient

We describe a statistical hypothesis test for the congruence coefficient under the null hypothesis that the coefficient is zero. The statistical significance of the congruence coefficient is determined based on the angle between two unit vectors on the $N$-dimensional unit spere, where the dimension $N$ depends on the input matrices. We first give a formula for general non-square matrices, and then we describe a statistical hypothesis test for symmetric and sparse matrices.

First of all, for notational convenience we recode an $X \in \mathbb{R}^{m \times n}$ matrix into a normalized vectors of size $nm$.

$$\vec{v}_X = \frac{vec(X)}{\|vec(X)\|}$$

where $\|\vec{v}\| = \sqrt{\vec{v} \cdot \vec{v}} = \sqrt{tr(XX)}$ is the Euclidean norm of $\vec{v}$ and

$$vec(X) = [X_{11}, X_{12}, ..., X_{1m}, X_{21}, X_{22}, ..., X_{2m}, ..., X_{mm}]$$

is the *vectorization* of matrix $X$, defined by

$$vec(X)_{i(m-1)+j} = X_{ij} \text{ for } 1 \leq i \leq m, 1 \leq j \leq n$$

Given two matrices $X, Y \in \mathbb{R}^{m \times n}$, the congruence coefficient can be recoded as the dot product between the two unit vectors $\vec{v}_X$ and $\vec{v}_Y$.

$$r_c(X,Y) = \frac{tr(XY^T)}{\sqrt{XX^T}\sqrt{YY^T}} = \frac{vec(X) \cdot vec(Y)}{\|vec(X)\|\|vec(Y)\|} = \vec{v}_X \cdot \vec{v}_Y$$

The two unit vectors $\vec{v}_X, \vec{v}_Y$ can be seen as points on the surface of the $N$-dimensional unit sphere, where $N = |\vec{v}_X| = |\vec{v}_Y| = mn$ is the size of the two vectors. In the same spirit, the congruence coefficient can be seen as the cosine of the angle $\theta$ between the two unit vectors

$$r_c(X,Y) = \vec{v} \cdot \vec{v} = \|\vec{v}_X\| \, \|\vec{v}_Y\| \cos(\theta) = \cos(\theta)$$

Note that for any given $N$-dimensional unit vector $\vec{v}$, there are infinite matrices $X \in \mathbb{R}^{m \times n}$ such that $\vec{v}_X = \vec{v}$, e.g. for any constant $a \in \mathbb{R}, a \neq 0$

$$\vec{v}_{aX} = \vec{v}_X \text{ and } r_c(aX, Y) = r_c(X, Y)$$

hence, the surface of the $N$-dimensional unit sphere is completely represented by $m \times m$ matrices. The statistical hypothesis testing for the congruence coefficient can be then recoded as a statistical hypothesis testing on the angle between two unit vectors on the $N$-dimensional unit sphere, under the null hypothesis that the two vectors are orthogonal. Strangely enough, for the unit $N$-dimensional sphere the surface area reaches a maximum and then decreases toward zero as the dimension $N$ increases. As a counter effect, if we fix a reference surface point $\vec{u}$ (e.g. $\vec{u} = \{1/\sqrt{N}, ..1/\sqrt{N}\}, |\vec{v}| = N$) and an angle $\theta$, the density of surface points $\vec{v}$, such that $\vec{u} \cdot \vec{v} \geq \cos(\theta)$, decreases as $N$ increases. Hence, a statistical hypothesis test on the angle between two unit vectors on the $N$-dimensional unit sphere need to include the degree of freedom $N$ in order to estimate the probability of rejecting the null hypothesis. The simples solution is to consider the fraction of the area of the hyper-spherical cap identified by the angle between the $\vec{v}_X, \vec{v_Y}$ vectors (see Fig. 1.a).

The area of the $N$-dimensional unit sphere is defined by

$$A_N = \frac{2\pi^{N/2}}{\Gamma(N/2)} \tag{19}$$

where $\Gamma$ is the *gamma function*. The area of the hyper-spherical cap identified by some $\theta$ angle on a $N$-dimensional unit sphere is defined by [Li, 2011]

$$A_N^{cap(\theta)} = \frac{1}{2} A_N I_{\sin^2(\theta)} \left( \frac{N-1}{2}, \frac{1}{2} \right) \tag{20}$$

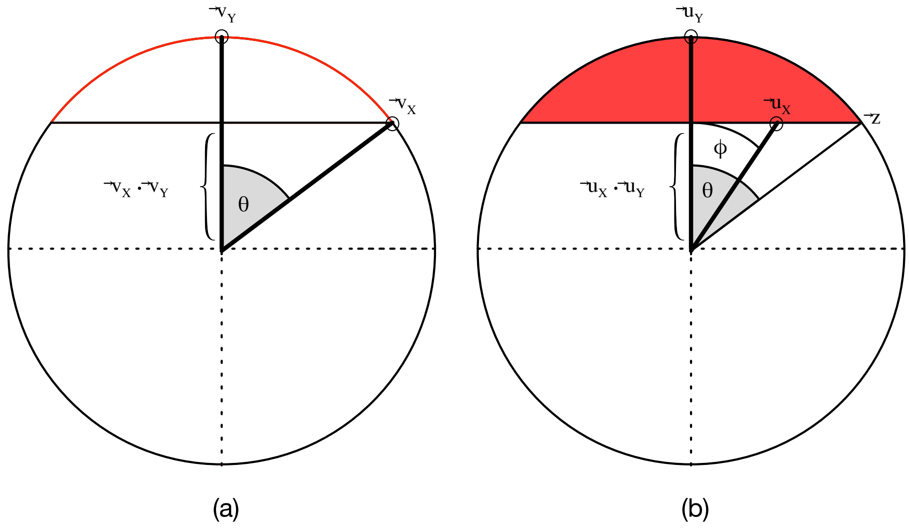


Figure 1: Hyperspherical caps: Area (a) and Volume (b)

where $I$ is the *regularized incomplete beta function*. By combining Equations 19 and 20 and by using the well known trigonometric equivalences

$$\theta = \arccos(\cos(\theta)), \quad \sin^2(\theta) = 1 - \cos^2(\theta)$$

we can define the two-tailed test of the congruence coefficient as the ratio between the hyperspherical cap area and the total sphere area:

$$Pr(|r_c| > p) = \frac{A_N^{cap(\arccos(p))}}{A_N} = I_{1-p^2}\left(\frac{N-1}{2}, \frac{1}{2}\right) \tag{21}$$

where $p = r_c(X, Y) = \cos(\theta)$ and $N = mn$ is the degree of freedom. The right-tailed statistical test is defined by:

$$Pr(r_c > p) = \begin{cases} \frac{1}{2} I_{1-p^2}\left(\frac{N-1}{2}, \frac{1}{2}\right) & p \geq 0 \\ \\ 1 - \frac{1}{2} I_{1-p^2}\left(\frac{N-1}{2}, \frac{1}{2}\right) & p < 0 \end{cases} \tag{22}$$

As it happens for the Pearson's correlation statistical test, the drawback of Equations 21 and 22 is that for large degrees of freedom (i.e. large matrices) small coefficients different from 0 are highly statistically significant. However, the degree of freedom can be reduced in some special cases. For example, if two matrices $X, Y \in \mathbb{R}^{m \times m}$ are symmetric and their first $k$ main diagonals are zero, the dimensionality $N$ can be reduced to $(m-k+1)(m-k)/2$ since the congruence coefficient can be recoded for the non-zero half of the matrices. For instance, if $vec(X)$ (resp. $vec(Y)$) is the vectorization of the the upper triangular part of $X$, starting from diagonal $k+1$,

$$vec(X) = [X_{1(k+1)}, X_{1(k+2)}, ..., X_{1m}, X_{2(k+2)}, X_{2(k+3)}, ..., X_{2m}, ...]$$

it is easy to see that $\vec{v}_X = \frac{vec(X)}{\|vec(X)\|}$ is a point on the surface of the $N$-dimensional unit sphere, where $N = (m-k+1)(m-k)/2$. Furthermore,

$$tr(XY) = 2\ vec(X) \cdot vec(Y) \text{ and } tr(XX) = 2\ vec(X) \cdot vec(X)$$

which implies the following equivalence:

$$r_c(X,Y) = \frac{2\ vec(X) \cdot vec(Y)}{\sqrt{2\ \|vec(X)\|}\sqrt{2\ \|vec(Y)\|}} = \vec{v}_X \cdot \vec{v}_Y$$

An even stronger reduction of the degree of freedom can be achieved for sparse symmetric matrices. In this case the statistical hypothesis test is based on the volume of the $N$-dimensional unit sphere. Formally, let $X, Y \in R^{m \times m}$ be two symmetric matrices with (at least) zero main diagonal and let $\vec{v}_X, \vec{v}_Y$ the normalized vectors corresponding to their upper triangular parts, respectively. Assume that $Y$ is a sparse matrix and let $\mathcal{I}$ be the set of indices, defined by

$$\mathcal{I} \subset \{1, ..., |\vec{v}_Y|\} \text{ such that } i \in \mathcal{I} \text{ if and only if } (\vec{v}_Y)_i \neq 0$$

That is, $\mathcal{I}$ encodes the indices of the non-zero elements in $\vec{v}_Y$, that correspond to the non-zero elements in the upper triangular part of matrix $Y$. Now, let $\vec{u}_X$ and $\vec{u}_Y$ be the sub-vectors of $\vec{v}_X$ and $\vec{v}_Y$, respectively, corresponding to $\mathcal{I}$ indices:

$$\vec{u}_X = \{(\vec{v}_X)_{I_1}, (\vec{v}_X)_{I_2}, ...\} \text{ and } \vec{u}_Y = \{(\vec{v}_Y)_{I_1}, (\vec{v}_Y)_{I_2} ...\}$$

It is easy to see that, since we removed only the zero elements in $\vec{v}_Y$, and the corresponding elements in $\vec{v}_X$,

$$r_c(X,Y) = \vec{v}_X \cdot \vec{v}_Y = \vec{u}_X \cdot \vec{u}_Y$$

and

$$0 \leq \|\vec{u}_X\| \leq \|\vec{u}_Y\| = 1$$

That is, $\vec{u}_Y$ has still length 1, while $\vec{u}_X$ can have a length smaller than 1. The two vectors $\vec{u}_X, \vec{u}_Y$ can then be seen as an inner point and a surface point, respectively, of the $N$-dimensional unit sphere, where $N = |\vec{u}_X| = |\vec{u}_Y|$ is the number of non-zero elements in the upper triangular part of $Y$. As before, for any given vector $\vec{u}$ such that $0 \leq \|\vec{u}\| \leq 1$ and $|\vec{u}| = N$, there are infinite symmetric matrices $X$, such that $\vec{u}_X = \vec{u}$. Then all the surface and inner point of the $N$-dimensional unit sphere are completely covered by symmetric matrices.

The vector norm $\|\vec{u}_X\|$ and the angle $\phi$ between $\vec{u}_X$ and $\vec{v}_Y$ identify a hyper-spherical cap on the $N$-dimensional unit sphere (see Fig. 1.b). The angle $\theta$, defining the hyper-spherical cap, is easily computed by using some geometric properties of the dot product between two vectors. First of all,

$$r_c(X,Y) = \vec{u}_X \cdot \vec{u}_Y = \|\vec{u}_X\|\,\|\vec{u}_Y\| \cos(\phi) = \|\vec{u}_X\| \cos(\phi)$$

where $\phi$ is the angle between $\vec{u}_X$ and $\vec{u}_Y$, and $\|\vec{u}_X\| \cos(\phi)$ is the length of the scalar projection of the $\vec{u}_X$ vector in the direction of the $\vec{u}_Y$ vector (see Fig. 1.b). We need to identify the hyper-spherical cap containing all (inner and surface) points $\vec{w}$ of the sphere such that $\vec{w} \cdot \vec{u}_Y \geq \vec{u}_X \cdot \vec{u}_Y$, i.e. the length of the scalar projection of $\vec{w}$ in the direction of $\vec{u}_Y$ is greater than or equal to $\|\vec{u}_X\| \cos(\phi)$. Such hyper-spherical cap is identified by the angle between any surface point, say $\vec{z}$, such that (see Fig. 1.b)

$$\vec{z} \cdot \vec{u}_Y = \vec{u}_X \cdot \vec{u}_Y = \|\vec{u}_X\| \cos(\phi).$$

Then we have that:

$$\cos(\theta) = \frac{\vec{z}}{\|\vec{z}\|} \cdot \frac{\vec{u}_Y}{\|\vec{u}_Y\|} = \vec{z} \cdot \vec{u}_Y = \vec{u}_X \cdot \vec{u}_Y = r_c(X,Y)$$

Then, if $Y$ is a symmetric and sparse matrix, we can compute the statistical significance of the $r_c(X,Y)$ coefficient as the ratio between the volume of the $N$-hyper-spherical cap identified by the angle $\theta$ and the volume of the $N$-dimensional unit sphere, where $\cos(\theta) = r_c(X,Y)$

and $N$ is the number of non-zero elements in the upper triangular part of $Y$. The volume of the $N$-dimensional unit sphere is defined by

$$V_N = \frac{\pi^{N/2}}{\Gamma(N/2+1)} \tag{23}$$

where $\Gamma$ is the *gamma function*. The volume of the hyper-spherical cap identified by angle $\theta$ in a $N$-dimensional unit sphere is defined by [Li, 2011]

$$V_N^{cap(\theta)} = \frac{1}{2} V_N I_{\sin^2(\theta)}\left(\frac{N+1}{2}, \frac{1}{2}\right) \tag{24}$$

where $I$ is the *regularized incomplete beta function*. By combining Equations 23 and 24 we obtain the two-tailed statistical significance formula for the congruence coefficient:

$$P(|r_c| > p) = \frac{V_N^{cap(\arccos p)}}{V_N} = I_{1-p^2}\left(\frac{N+1}{2}, \frac{1}{2}\right) \tag{25}$$

where $p = r_c(X,Y)$ and $N$ is the number of non-zero elements in the upper triangular part of $Y$. The right-tailed formula is defined by:

$$Pr(r_c > p) = \begin{cases} \frac{1}{2} I_{1-p^2}\left(\frac{N+1}{2}, \frac{1}{2}\right) & p \geq 0 \\ 1 - \frac{1}{2} I_{1-p^2}\left(\frac{N+1}{2}, \frac{1}{2}\right) & p < 0 \end{cases} \tag{26}$$

where, again, $p = r_c(X,Y)$ and $N$ is the number of non-zero elements in the upper triangular part of $Y$.

Note that Equations 25 and 26 can be used also for assessing the statistical significance of the congruence coefficient between two aligned maps $r_c^\alpha(X,Y)$, as defined in Equation 7. In this case, since $tr(XY^\alpha) = tr(X^{\alpha^{-1}}Y)$ the degree of freedom $N$ can be chosen to be the number of non-zero elements in the upper triangular part of either $X \in \mathbb{R}^{m\times m}$ or $Y \in \mathbb{R}^{n\times n}$.

In database searches, if $X$ is the target map, $Y$ some template map and $\alpha$ and alignments between $X$ and $Y$, by choosing

$$N = \textit{ number of non-zero elements in the upper triangular part of } X$$

we have that $Pr(r_c > r_c^\alpha(X,Y))$ is equivalent to computing $Pr(r_c > r_c^\alpha(X,Y) \mid X)$, which gives the probability of (uniformly) sampling a random matrix $Y' \in \mathbb{R}^{m\times m}$ such that

$$r_c(X, Y') > r_c^\alpha(X, Y).$$

Then, given two symmetric matrices with zero main diagonal $X, Y$ and an alignment $\alpha$ between $X$ and $Y$, we can detect whether there is a statistically significantly similarity (at standard significance level 0.05) between $X$ and $Y$ if

$$Pr(r_c > r_c^\alpha(X, Y) \mid X) < 0.05 \text{ and } Pr(r_c > r_c^\alpha(X, Y) \mid Y) < 0.05$$

We remark that, if $Y \in \mathbb{R}^{n \times n}$ (resp. for $X$) is a distance matrix, only the main diagonal is zero, then the degree of freedom $N = n * (n-1)/2$ can be quite large for large sizes $n$. This implies that, every congruence coefficient $r_c^\alpha(X, Y)$ sufficiently larger than 0 is statistically significant but this does not necessarily mean that $X$ and $Y$ are statistically significantly similar. The issue here is that

$$Pr(r_c > r_c^\alpha(X, Y) \mid Y)$$

gives the probability of observing a congruence coefficient greater than $r_c^\alpha(X, Y)$ if $X \in \mathbb{R}^{m \times m}$ is randomly sampled. In real case scenarios, $X$ can be a real or predicted distance map, while a randomly sampled $m \times m$ matrix is generally not a physical distance map. That is, the space of (real/predicted) $m \times m$ distance maps is much smaller than the space of random $m \times m$ symmetric matrices. We have the same problem when $Y$ is a contact map, but in this case the typically low degree of freedom we use for contact maps can still detect non significant similarities between two maps.

### 7 Tests

#### 7.1 Template and Benchmark Data

Benchmark data sets were obtained from the CASP repository (`http://predictioncenter.org`). For contact-based fold recognition assessment, we selected all residue-residue contact predictions submitted to the CASP12 and CASP13 experiments. When a group submitted multiple models for the same target, we selected only the first model. For distance-based fold recognition assessment, we decided to *simulate* predicted distance maps by recovering them from the structural predictions at CASP12 and CASP13 (likewise using only the first model). This was necessary since distance map predictions were used as an intermediary step, rather than as a standalone problem, and such predictions were not available. We considered only the CASP targets for which the experimentally determined structure was available in the PDB (`http://www.rcsb.org/`) and the fold annotation was available in the ECOD classification [Cheng *et al.*, 2014].

Template data were obtained from the ECOD database (version develop 238, 06/14/2019). ECOD provides a hierarchical classification of protein domains, from experimentally determined PDB protein structures, according to their evolutionary relationships. Protein domains are classified with respect to four groups: the *F-group* groups domains with significant sequence similarity, the *T-group* groups domains with similar topological connections, the *H-group* groups domain that are considered homologous based on different attributes (e.g. functional similarity, literature), the *X-group* groups domains that are potentially homologous although there is not yet adeguate evidence to support their homology relationship. The ECOD pipeline uses of the structural alignment tool DaliLite [Holm and Park, 2000] only if a protein chain remains unclassified after sequence- similarity-based searches. The ECOD database provides a broader coverage of the CASP targets compared to other structural classifications, such as SCOPe [Chandonia *et al.*, 2018].

We downloaded the ECOD pre-filtered subset at 40% sequence identity, consisting of 29,512 pdb-style domain files. We added 34 CASP13-related domains from the complete ECOD distribution which were not available in the subset. In order to prevent any sequence homology bias in our tests, we removed from the ECOD dataset all protein domains found by a hmmsearch [Eddy, 2011] scan of the CASP12 and CASP13 targets against the ECOD database. We further removed any domains shorter than 15 residues and domain files containing errors preventing parsing. In order to identify a subset of hard targets, we matched the FM (Free Modeling) domains of the CASP targets with the domains identified in ECOD. Performances on the FM targets were assessed on their FM domains only. Native contact and distance maps were extracted from the ECOD pdb domain files.

In predicted and native maps, we did not take into account contacts or distances between residues with sequence separation less than 6, as these are not informative of protein tertiary structure. The corresponding entries were set to 0. Furthermore un order to give more weight to short distances in Equation (5), we transformed the distance maps by taking the reciprocal of each distance (distances equal to 0 remain unchanged). Such a transformation basically converts distance maps into *weighted* contact maps. Preliminary tests showed that, on average, such transformation provides slightly better performance over using the raw distances (data not shown).

The exact number of ECOD templates used in our experiments, as well as the targets and FM targets in the two CASP benchmark sets, are shown in Table 1.

#### 7.2 Benchmark software

We considered three heuristic contact map overlap programs for performance comparison: AlEigen [Di Lena *et al.*, 2010], EigenThreader [Buchan and Jones, 2017], and Map_Align [Ovchinnikov *et al.*, 2017]. AlEigen, EigenThreader, and Map_Align have their own specific

| Dataset | #Targets (#FM) | #RR Pred | #REG Pred | #ECOD templates |
|---|---|---|---|---|
| CASP12 | 34 (12) | 1,109 | 2,567 | 27,677 |
| CASP13 | 24 (9) | 980 | 1,880 | 27,721 |

Table 1: Benchmark set statistics. #Targets: number of CASP targets with fold annotation. #FM: number of targets containing FM domains. RR Pred: residue-residue contact predictions. REG Pred: regular structure predictions. #ECOD templates: number of sequence homology-free ECOD templates

| Method | Avg Time CASP12 | Avg Time CASP13 |
|---|---|---|
| EigenThreader | 41m | 52m |
| AlEigen | 2.8h | 3.9h |
| Map_Align | 6d | 8.7d |
| CE | 4.7d | 4.9d |
| TM-align | 2.2h | 2.2h |

Table 2: Average running time per prediction. m=minutes, h=hours, d=days

scoring schemes for database searches. AlEigen simply uses the CMO (Contact Map Overlap) score [Godzik *et al.*, 1992, Goldman *et al.*, 1999], which is basically the trace of the product between two aligned matrices, i.e. an unnormalized version of the congruence coefficient score in Equation (5). Map_align score is also based on the CMO score, the only difference being that the matching contacts are weighted according to their sequence separation and the final score takes into account the number of gaps in the alignment. EigenThreader uses a score based on the t-test of the Pearson correlation between contact predictions and contact distances in the template. The EigenThreader score is actually a weighted function, fitted on some training data, of three variables: the t-statistic, the fraction of the target that is aligned, and the fraction of the template that is aligned. In some cases, EigenThreader's score function returns the same similarity score for thousands of different templates, which makes it impossible to fairly select the top-scored templates. For this reason, when selecting EigenThreader's top-scored templates, we use the t-statistic as a secondary sorting key.

All three tools return an alignment between two input contact maps. We used such alignments to calculate the corresponding congruence coefficient and its p-value, and thus re-score the overlap similarity between target and template maps with respect to statistically significant (at standard 0.05 threshold) $r_c$ scores.

The three contact map overlap tools were run as much as possible with standard/default parameters in order to make the comparison fair and the running times reasonable. AlEigen and EigenThreader approximate a two-dimensional map alignment by computing a one-dimensional alignment between sets of eigenvectors associated with the maps. With both AlEigen and EigenThreader we use 7 eigenvectors (the running time increases with the number of eigenvectors). Map_Align exploits an iterative double dynamic programming approach. It was run by limiting the number of iterations to 5, instead of the suggested 20, due to the long running time required to process a target against the ECOD dataset (see Table 2).

Unlike the maximum CMO problem, the standalone distance map alignment problem has received little or no attention in the literature. To the best of our knowledge, distance map alignment has only been used as a preprocessing step in some structural alignment tools, such as DaliLite [Holm and Park, 2000]. For this reason, for performance comparison, we decided to use two popular structural alignments tools, CE [Shindyalov and Bourne, 1998] and TM-Align [Zhang and Skolnick, 2005], and recover the distance map alignments from the structural alignments computed by both tools. Our choice of structural alignment tools has been driven mainly by speed considerations due to the very large number of comparison performed in our tests (over 52M comparisons). The fold recognition performances of CE are assessed by considering the Z-scores as the primary key, and the CE raw scores as the secondary key. For the TM-align performance, we use the TM-score [Xu and Zhang, 2010].

The average running times of the benchmarked methods are summarized in Table 2.

#### 7.3 Approximation of expectation and variance

We compare the true mean and standard deviation of the congruence coefficient (over all possible alignments) computed for all pairs of targets and templates in the CASP12 and CASP13 datasets against sampling/permutation mean and standard deviation. Due to the large computational times needed to compute the true standard deviation, in the comparison tests we just consider the true standard deviation computed for target and templates of small sizes: all templates of size $\leq 50$ and targets of size $\leq 150$ for contact maps, and all templates of size $\leq 50$ and targets of size $\leq 100$ for distance maps. Conversely, the true mean has been computed for all pairs of targets and templates.

For a pair $X \in \mathbb{R}^{m\times m}$ (target map) and $Y \in \mathbb{R}^{n\times n}$ (template map), the mean and standard deviation of the congruence coefficient with respect to all permutations of $Y$ has been computed with the closed formulas in Section 5, with $N = m + n$. The sampling mean and standard deviation has been computed by generating random alignments in the following way. Given two symmetric matrices $X \in \mathbb{R}^{m\times m}$ and $Y \in \mathbb{R}^{n\times n}$ we have that (see Section 4.1)

$$f_k(m, n) = \frac{\binom{m}{k}\binom{n}{k}}{\binom{m+n}{n}},\ 0 \leq k \leq \min\{m, n\}$$

| Comparison | Type | R | MAE | MAPE | MaxAE |
|---|---|---|---|---|---|
| True mean vs Sampling mean | Contact maps | 0.999 | 0.00004 | 0.55% | 0.0012 |
| True Sd vs Sampling Sd | | 0.999 | 0.00010 | 1.10% | 0.0052 |
| True mean vs Permutation mean | | 0.856 | 0.00217 | 31.50% | 0.0586 |
| True Sd vs Permutation Sd | | 0.851 | 0.00190 | 26.60% | 0.0592 |
| True mean vs Sampling mean | Distance maps | 0.999 | 0.00013 | 0.04% | 0.0011 |
| True Sd vs Sampling Sd | | 0.999 | 0.00010 | 0.35% | 0.0008 |
| True mean vs Permutation mean | | 0.993 | 0.00779 | 5.8% | 0.0979 |
| True mean vs Permutation mean | | 0.968 | 0.00320 | 11.33% | 0.0177 |

Table 3: R = Pearson Correlation, MAE = Mean Absolute Error, MAPE = Mean Absolute Percentage Error, MaxAE = Maximum Absolute Error

is the fraction of alignments that match exactly $k$ positions in $X$ with $k$ positions in $Y$. If $K$ is the number of random alignments that we want to sample, for each $0 \leq k \leq \min\{m, n\}$, we randomly sample $\lceil f_k(m, n) * K \rceil$ distinct alignments. That is, the number of sampled alignments with respect to matching size $k$ is proportional to the fraction of total alignments of size $k$ between $X$ and $Y$. The mean and standard deviation of the congruence coefficient has been estimated by computing the $r_c^\alpha(X, Y)$ for each sampled alignment $\alpha$. In our tests we fix $K = 10,000$ for all target/template pairs.

The scatter plot of the comparison true vs sampling/permutation mean and standard deviation for contact maps are in Figures 2 and 3, respectively. The equivalent scatter plots for distance maps are in Figures 4 and 5. It is clear that sampling mean and standard deviation from 10k random alignments are a very good approximation of the true mean and standard deviation. This is also confirmed by the statistical measures shown in Table 3, where we can see that the errors of the sampling mean/sd are on the average less than 1% of the true value (MAPE metrics). Conversely, the permutation mean and standard deviations do not provide good approximations, although its mean absolute error (MAE) is quite small (see Table 3).

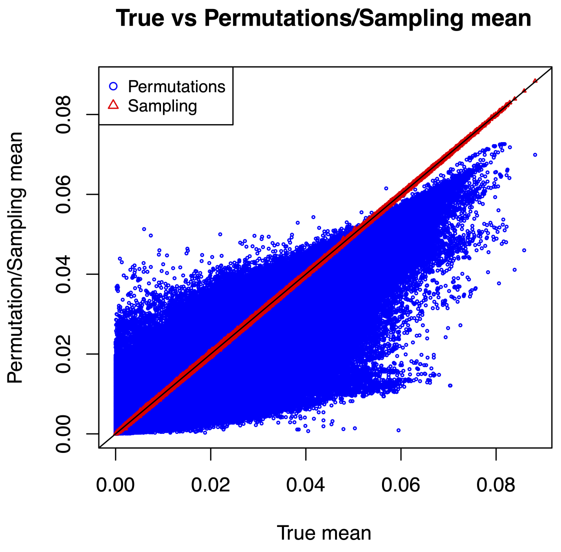


Figure 2: Contact maps: mean

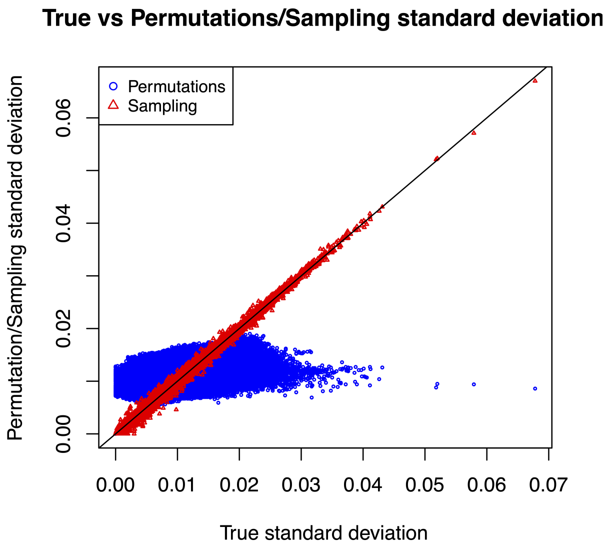


Figure 3: Contact maps: standard deviation

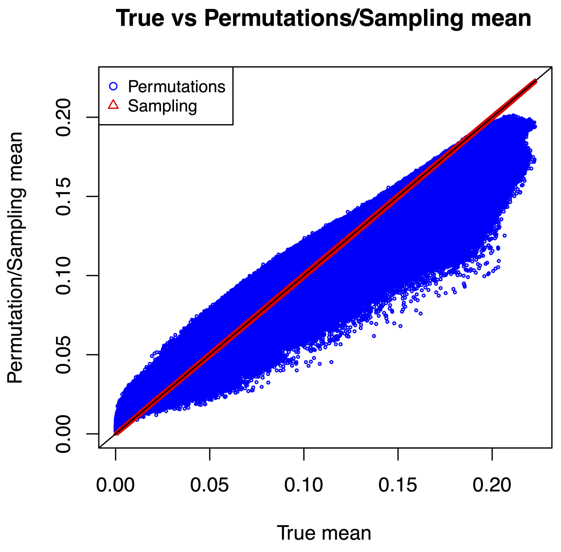


Figure 4: Distance maps: mean

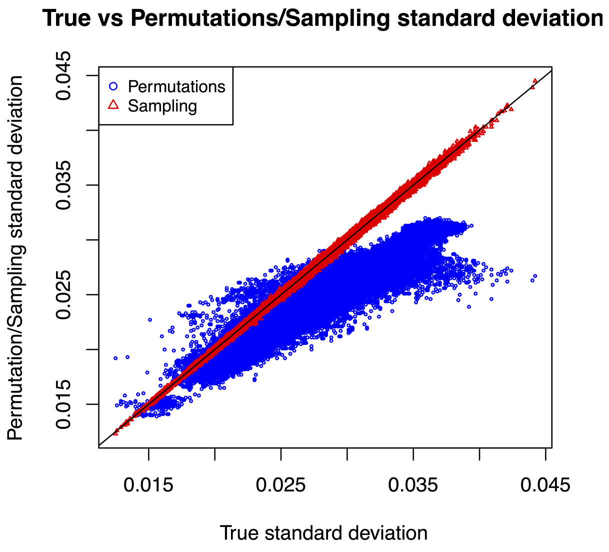


Figure 5: Distance maps: standard deviation

#### 7.4 Fold recognition performances

##### 7.4.1 Fold recognition with predicted contacts

For performance comparison, we search all residue-residue contact predictions submitted at CASP for a single target against the ECOD templates. This implies a maximum number of 38 predictions per target at CASP12 and 46 at CASP13, i.e. the number of residue-residue prediction groups in the two CASP editions. We consider only the first model if a single group submitted more predictions for the same target. True Positive Rate (TPR) fold recognition performances have been assessed by selecting the top-1, top-5, top-10 and top-20 unique templates identified by the search with multiple predicted maps. For each top-$k$ set, the TPR score is computed by counting the fraction of targets for which at least one template with similar fold is in the top-$k$ hits. We assess the TPR performances separately for the four ECOD classes, *Family Level* (F), *Topology Level* (T), *Homology Level* (H) and *Possible Homology Level* (X), which implies that, for example, for TPR assessment at the Family Level we consider only the CASP targets that have been annotated at the Family Level in ECOD. The TPR performances on CASP12 and CASP13 benchmark sets, with respect to the three map alignment tools AlEigen, EigenTHREADER and Map_Align are summarized in Table 4. The TPR performances restricted to FM (Free Modelling) targets only are shown in Table 5. The two tables compares the performances of the three tools with their specific scoring schemes against those obtained by using the congruence coefficient, indicated by AlEigen+$r_c$, EigenTHREADER+$r_c$ and Map_Align+$r_c$, respectively. In Tables 4 and 5 we can notice that the fold recognition precision is overall dramatically improved by the usage of the congruence coefficient for fitness ranking. We point out that in Table 5, fold recognition at the family level should be ignored since FM targets are classified as completely new folds. However, we found that a couple of targets for the older CASP12 benchmark sets have been annotated at the family level in ECOD. None of the benchmarked approaches can identify such related templates.

In Tables 6-9 we consider fold recognition performance by restricting contact predictions to the top-5, top-10, top15 and top-20 best performing contact predictors at CASP12 and CASP13 (obtained by the official CASP rankings). Best predictors provide generally better contact predictions, which should improve fold recognition performances. In fact, in Table 6-9, we can notice a general improvements in fold recognition accuracy for all methods with their original ranking scores. Interestingly, the improvement is not relevant (or even absent) with the usage of the congruence coefficient, which provides, anyway, the best fold recognition performances.

| | | top-1 hit | | | | top-5 hits | | | | top-10 hits | | | | top-20 hits | | | |
|---|---|---|---|---|---|---|---|---|---|---|---|---|---|---|---|---|---|
| Method | Dataset | F | T | H | X | F | T | H | X | F | T | H | X | F | T | H | X |
| AlEigen | CASP12 | 0.00 | 0.07 | 0.07 | 0.06 | 0.00 | 0.07 | 0.07 | 0.09 | 0.00 | 0.07 | 0.10 | 0.12 | 0.00 | 0.07 | 0.10 | 0.15 |
| AlEigen+$r_c$ | | **0.25** | 0.29 | 0.37 | 0.38 | **0.25** | 0.39 | 0.43 | 0.47 | **0.25** | 0.39 | 0.47 | 0.56 | **0.25** | 0.54 | 0.57 | 0.62 |
| EigenTHREADER | | 0.00 | 0.00 | 0.00 | 0.00 | 0.00 | 0.07 | 0.07 | 0.06 | 0.00 | 0.07 | 0.07 | 0.09 | 0.00 | 0.11 | 0.10 | 0.15 |
| EigenTHREADER+$r_c$ | | **0.25** | 0.32 | 0.40 | 0.41 | **0.25** | 0.54 | 0.57 | 0.62 | **0.25** | 0.57 | 0.60 | 0.68 | **0.25** | 0.61 | 0.63 | 0.68 |
| Map_Align | | 0.00 | 0.07 | 0.07 | 0.06 | 0.00 | 0.07 | 0.07 | 0.06 | 0.00 | 0.07 | 0.10 | 0.12 | 0.00 | 0.07 | 0.10 | 0.15 |
| Map_Align+$r_c$ | | **0.25** | **0.57** | **0.60** | **0.65** | **0.25** | **0.61** | **0.63** | **0.65** | **0.25** | **0.68** | **0.70** | **0.71** | **0.25** | **0.71** | **0.73** | **0.74** |
| AlEigen | CASP13 | 0.00 | 0.15 | 0.23 | 0.25 | 0.00 | 0.25 | 0.32 | 0.38 | 0.00 | 0.25 | 0.32 | 0.42 | 0.00 | 0.25 | 0.32 | 0.42 |
| AlEigen+$r_c$ | | 0.00 | 0.35 | 0.36 | 0.46 | 0.00 | 0.45 | 0.45 | 0.62 | 0.00 | 0.60 | 0.59 | 0.67 | 0.50 | 0.70 | 0.73 | 0.79 |
| EigenTHREADER | | 0.00 | 0.10 | 0.09 | 0.12 | 0.00 | 0.15 | 0.18 | 0.25 | 0.00 | 0.15 | 0.18 | 0.29 | 0.00 | 0.20 | 0.23 | 0.33 |
| EigenTHREADER+$r_c$ | | 0.00 | 0.50 | 0.50 | 0.58 | 0.00 | 0.70 | 0.68 | **0.71** | 0.00 | 0.80 | 0.77 | 0.79 | 0.00 | 0.80 | 0.77 | **0.83** |
| Map_Align | | **0.50** | 0.35 | 0.41 | 0.50 | 0.50 | 0.40 | 0.45 | 0.54 | 0.50 | 0.40 | 0.45 | 0.54 | 0.50 | 0.40 | 0.45 | 0.54 |
| Map_Align+$r_c$ | | **0.50** | **0.60** | **0.59** | **0.62** | **1.00** | **0.75** | **0.73** | **0.71** | **1.00** | **0.85** | **0.82** | **0.83** | **1.00** | **0.85** | **0.86** | **0.83** |

Table 4: **Fold recognition performances with predicted contacts from all contact predictors**. True Positive Rate (TPR) fold recognition performances on CASP12 and CASP13 benchmark sets. The TPR performances are assessed with respect to the **top-1**, **top-5**, **top10** and **top-20** ranked hits. ECOD hierarchy: (F) Family Level (4 targets in CASP12, 2 targets in CASP13), (T) Topology Level (28 targets in CASP12, 20 targets in CASP13), (H) Homology Level (30 targets in CASP12, 22 targets in CASP13), (X) Possible Homology Level (34 targets in CASP12, 24 targets in CASP13). EigenTHREADER, Map_Align and AlEigen use their own scoring system EigenTHREADER+$r_c$, Map_Align+$r_c$ and AlEigen+$r_c$ use statistically significant congruence coefficient. Best TPR performances per column on CASP12 and CASP13 benchmark sets are highlighted in bold.

| | | top-1 hit | | | | top-5 hits | | | | top-10 hits | | | | top-20 hits | | | |
|---|---|---|---|---|---|---|---|---|---|---|---|---|---|---|---|---|---|
| Method | Dataset | F | T | H | X | F | T | H | X | F | T | H | X | F | T | H | X |
| AlEigen | CASP12 | 0.00 | **0.14** | 0.12 | 0.08 | 0.00 | **0.14** | 0.12 | 0.08 | 0.00 | 0.14 | 0.12 | 0.08 | 0.00 | 0.14 | 0.12 | 0.08 |
| AlEigen+$r_c$ | | 0.00 | 0.00 | 0.12 | 0.17 | 0.00 | 0.00 | 0.12 | 0.17 | 0.00 | 0.00 | 0.25 | 0.33 | 0.00 | 0.14 | 0.25 | 0.33 |
| EigenTHREADER | | 0.00 | 0.00 | 0.00 | 0.00 | 0.00 | 0.00 | 0.00 | 0.00 | 0.00 | 0.00 | 0.00 | 0.08 | 0.00 | 0.00 | 0.00 | 0.17 |
| EigenTHREADER+$r_c$ | | 0.00 | 0.00 | 0.12 | 0.17 | 0.00 | **0.14** | **0.25** | **0.25** | 0.00 | 0.29 | 0.38 | **0.42** | 0.00 | 0.29 | 0.38 | **0.42** |
| Map_Align | | 0.00 | **0.14** | 0.12 | 0.08 | 0.00 | **0.14** | 0.12 | 0.08 | 0.00 | 0.14 | 0.12 | 0.08 | 0.00 | 0.14 | 0.12 | 0.08 |
| Map_Align+$r_c$ | | 0.00 | **0.14** | **0.25** | **0.25** | 0.00 | **0.14** | **0.25** | **0.25** | 0.00 | **0.43** | **0.50** | **0.42** | 0.00 | **0.43** | **0.50** | **0.42** |
| AlEigen | | 0.00 | 0.14 | **0.25** | **0.22** | 0.00 | 0.14 | 0.25 | 0.22 | 0.00 | 0.14 | 0.25 | 0.22 | 0.00 | 0.14 | 0.25 | 0.22 |
| AlEigen+$r_c$ | | 0.00 | 0.14 | 0.12 | **0.22** | 0.00 | 0.29 | 0.25 | **0.44** | 0.00 | 0.29 | 0.25 | 0.44 | 0.00 | 0.43 | 0.38 | 0.56 |
| EigenTHREADER | | 0.00 | 0.14 | 0.12 | 0.11 | 0.00 | 0.14 | 0.12 | 0.11 | 0.00 | 0.14 | 0.12 | 0.11 | 0.00 | 0.14 | 0.12 | 0.11 |
| EigenTHREADER+$r_c$ | | 0.00 | 0.14 | 0.12 | **0.22** | 0.00 | **0.43** | **0.38** | **0.44** | 0.00 | **0.57** | **0.50** | **0.67** | 0.00 | **0.57** | 0.50 | **0.67** |
| Map_Align | | 0.00 | 0.14 | **0.25** | **0.22** | 0.00 | 0.14 | 0.25 | 0.22 | 0.00 | 0.14 | 0.25 | 0.22 | 0.00 | 0.14 | 0.25 | 0.22 |
| Map_Align+$r_c$ | | 0.00 | **0.29** | **0.25** | **0.22** | 0.00 | 0.29 | 0.25 | 0.33 | 0.00 | **0.57** | **0.50** | 0.56 | 0.00 | **0.57** | **0.62** | 0.56 |

Table 5: **Fold recognition performances on FM targets with predicted contacts from all contact predictors**. True Positive Rate (TPR) fold recognition performances on CASP12 and CASP13 benchmark sets. The TPR performances are assessed with respect to the **top-1**, **top-5**, **top10** and **top-20** ranked hits. ECOD hierarchy: (F) Family Level (2 targets in CASP12, 0 targets in CASP13), (T) Topology Level (7 targets in both CASP12 and CASP13), (H) Homology Level (8 targets in both CASP12 and CASP13), (X) Possible Homology Level (12 targets in CASP12, 9 targets in CASP13). EigenTHREADER, Map_Align and AlEigen use their own scoring system EigenTHREADER+$r_c$, Map_Align+$r_c$ and AlEigen+$r_c$ use statistically significant congruence coefficient. Best TPR performances per column on CASP12 and CASP13 benchmark sets are highlighted in bold.

| | | top-1 hit | | | | top-5 hits | | | | top-10 hits | | | | top-20 hits | | | |
|---|---|---|---|---|---|---|---|---|---|---|---|---|---|---|---|---|---|
| Method | Dataset | F | T | H | X | F | T | H | X | F | T | H | X | F | T | H | X |
| AlEigen | CASP12 | 0.00 | 0.18 | 0.23 | 0.24 | 0.00 | 0.18 | 0.23 | 0.26 | 0.00 | 0.18 | 0.27 | 0.29 | 0.00 | 0.29 | 0.33 | 0.32 |
| AlEigen+$r_c$ | | 0.00 | 0.25 | 0.33 | 0.38 | 0.00 | 0.39 | 0.47 | 0.44 | 0.00 | 0.46 | 0.50 | 0.50 | 0.00 | 0.46 | 0.50 | 0.50 |
| EigenTHREADER | | 0.00 | 0.11 | 0.13 | 0.15 | 0.00 | 0.18 | 0.23 | 0.32 | 0.00 | 0.25 | 0.30 | 0.35 | 0.00 | 0.29 | 0.40 | 0.50 |
| EigenTHREADER+$r_c$ | | 0.00 | 0.29 | 0.33 | 0.38 | 0.00 | 0.39 | 0.43 | 0.44 | 0.00 | 0.39 | 0.43 | 0.44 | 0.00 | 0.43 | 0.47 | 0.50 |
| Map_Align | | 0.00 | 0.21 | 0.27 | 0.26 | 0.00 | 0.25 | 0.30 | 0.29 | 0.00 | 0.25 | 0.30 | 0.32 | 0.00 | 0.32 | 0.33 | 0.38 |
| Map_Align+$r_c$ | | 0.00 | **0.39** | **0.47** | **0.50** | 0.00 | **0.57** | **0.60** | **0.62** | 0.00 | **0.64** | **0.67** | **0.65** | 0.00 | **0.64** | **0.67** | **0.68** |
| AlEigen | CASP13 | 0.00 | 0.35 | 0.41 | 0.50 | **0.50** | 0.35 | 0.41 | 0.50 | 0.50 | 0.35 | 0.41 | 0.50 | **0.50** | 0.35 | 0.41 | 0.50 |
| AlEigen+$r_c$ | | **0.50** | 0.45 | 0.45 | 0.54 | **0.50** | 0.50 | 0.55 | 0.58 | **0.50** | 0.60 | **0.64** | 0.58 | **0.50** | 0.60 | 0.64 | 0.67 |
| EigenTHREADER | | 0.00 | 0.35 | 0.36 | 0.42 | 0.00 | 0.50 | 0.50 | 0.46 | 0.00 | **0.50** | 0.50 | 0.46 | 0.00 | 0.55 | 0.55 | 0.58 |
| EigenTHREADER+$r_c$ | | 0.00 | 0.35 | 0.41 | 0.54 | 0.00 | 0.50 | 0.55 | 0.54 | 0.00 | 0.55 | 0.59 | 0.54 | 0.00 | 0.55 | 0.59 | 0.54 |
| Map_Align | | 0.00 | 0.55 | **0.59** | **0.67** | **0.50** | 0.55 | 0.59 | **0.67** | **0.50** | 0.60 | **0.64** | **0.71** | **0.50** | 0.60 | 0.64 | 0.71 |
| Map_Align+$r_c$ | | **0.50** | 0.50 | 0.50 | 0.58 | **0.50** | **0.65** | **0.64** | 0.62 | **0.50** | **0.65** | **0.64** | **0.71** | **0.50** | **0.70** | **0.73** | **0.75** |

Table 6: **Fold recognition performances with predicted contacts from top-5 contact predictors**. True Positive Rate (TPR) fold recognition performances on CASP12 and CASP13 benchmark sets. The TPR performances are assessed with respect to the **top-1**, **top-5**, **top10** and **top-20** ranked hits. ECOD hierarchy: (F) Family Level (4 targets in CASP12, 2 targets in CASP13), (T) Topology Level (28 targets in CASP12, 20 targets in CASP13), (H) Homology Level (30 targets in CASP12, 22 targets in CASP13), (X) Possible Homology Level (34 targets in CASP12, 24 targets in CASP13). EigenTHREADER, Map_Align and AlEigen use their own scoring system EigenTHREADER+$r_c$, Map_Align+$r_c$ and AlEigen+$r_c$ use statistically significant congruence coefficient. Best TPR performances per column on CASP12 and CASP13 benchmark sets are highlighted in bold.

| | | top-1 hit | | | | top-5 hits | | | | top-10 hits | | | | top-20 hits | | | |
|---|---|---|---|---|---|---|---|---|---|---|---|---|---|---|---|---|---|
| Method | Dataset | F | T | H | X | F | T | H | X | F | T | H | X | F | T | H | X |
| AlEigen | CASP12 | 0.00 | 0.18 | 0.23 | 0.24 | 0.00 | 0.18 | 0.23 | 0.26 | 0.00 | 0.18 | 0.27 | 0.29 | 0.00 | 0.29 | 0.33 | 0.32 |
| AlEigen+$r_c$ | | 0.00 | 0.25 | 0.37 | 0.41 | 0.00 | 0.43 | 0.50 | 0.50 | 0.00 | 0.50 | 0.53 | 0.53 | 0.00 | 0.54 | 0.57 | 0.56 |
| EigenTHREADER | | 0.00 | 0.21 | 0.30 | 0.32 | 0.00 | 0.29 | 0.33 | 0.44 | 0.00 | 0.32 | 0.37 | 0.47 | 0.00 | 0.39 | 0.43 | 0.53 |
| EigenTHREADER+$r_c$ | | 0.00 | 0.36 | 0.40 | 0.44 | 0.00 | 0.46 | 0.50 | 0.53 | 0.00 | 0.54 | 0.57 | 0.56 | 0.00 | 0.57 | 0.60 | 0.59 |
| Map_Align | | 0.00 | 0.21 | 0.27 | 0.26 | 0.00 | 0.25 | 0.30 | 0.29 | 0.00 | 0.25 | 0.30 | 0.32 | 0.00 | 0.32 | 0.33 | 0.38 |
| Map_Align+$r_c$ | | 0.00 | **0.50** | **0.63** | **0.62** | 0.00 | **0.64** | **0.67** | **0.68** | 0.00 | **0.75** | **0.77** | **0.74** | 0.00 | **0.75** | **0.77** | **0.74** |
| AlEigen | CASP13 | 0.00 | 0.40 | 0.45 | 0.54 | 0.00 | 0.40 | 0.45 | 0.54 | **0.50** | 0.40 | 0.45 | 0.54 | **0.50** | 0.40 | 0.45 | 0.54 |
| AlEigen+$r_c$ | | **0.50** | 0.45 | 0.45 | 0.54 | **0.50** | 0.55 | 0.55 | 0.62 | **0.50** | 0.65 | 0.64 | 0.62 | **0.50** | 0.65 | 0.64 | 0.67 |
| EigenTHREADER | | 0.00 | 0.40 | 0.41 | 0.46 | 0.00 | 0.50 | 0.50 | 0.46 | 0.00 | 0.50 | 0.50 | 0.46 | 0.00 | 0.55 | 0.55 | 0.58 |
| EigenTHREADER+$r_c$ | | 0.00 | 0.50 | 0.55 | 0.62 | 0.00 | **0.70** | **0.73** | **0.71** | 0.00 | **0.75** | **0.77** | **0.75** | 0.00 | **0.75** | **0.77** | **0.75** |
| Map_Align | | 0.00 | **0.55** | **0.59** | **0.67** | **0.50** | 0.55 | 0.59 | 0.67 | **0.50** | 0.60 | 0.64 | 0.71 | **0.50** | 0.60 | 0.64 | 0.71 |
| Map_Align+$r_c$ | | **0.50** | **0.55** | 0.55 | **0.62** | **0.50** | **0.70** | 0.68 | **0.71** | **0.50** | 0.70 | 0.73 | 0.71 | **0.50** | **0.75** | **0.77** | **0.75** |

Table 7: **Fold recognition performances with predicted contacts from top-10 contact predictors**. True Positive Rate (TPR) fold recognition performances on CASP12 and CASP13 benchmark sets. The TPR performances are assessed with respect to the **top-1**, **top-5**, **top10** and **top-20** ranked hits. ECOD hierarchy: (F) Family Level (4 targets in CASP12, 2 targets in CASP13), (T) Topology Level (28 targets in CASP12, 20 targets in CASP13), (H) Homology Level (30 targets in CASP12, 22 targets in CASP13), (X) Possible Homology Level (34 targets in CASP12, 24 targets in CASP13). EigenTHREADER, Map_Align and AlEigen use their own scoring system EigenTHREADER+$r_c$, Map_Align+$r_c$ and AlEigen+$r_c$ use statistically significant congruence coefficient. Best TPR performances per column on CASP12 and CASP13 benchmark sets are highlighted in bold.

| | | top-1 hit | | | | top-5 hits | | | | top-10 hits | | | | top-20 hits | | | |
|---|---|---|---|---|---|---|---|---|---|---|---|---|---|---|---|---|---|
| Method | Dataset | F | T | H | X | F | T | H | X | F | T | H | X | F | T | H | X |
| AlEigen | CASP12 | 0.00 | 0.18 | 0.23 | 0.24 | 0.00 | 0.18 | 0.23 | 0.26 | 0.00 | 0.18 | 0.27 | 0.29 | 0.00 | 0.29 | 0.33 | 0.32 |
| AlEigen+$r_c$ | | 0.00 | 0.25 | 0.33 | 0.38 | 0.00 | 0.36 | 0.43 | 0.44 | 0.00 | 0.43 | 0.50 | 0.53 | 0.00 | 0.46 | 0.50 | 0.53 |
| EigenTHREADER | | 0.00 | 0.21 | 0.30 | 0.32 | 0.00 | 0.32 | 0.37 | 0.47 | 0.00 | 0.39 | 0.43 | 0.53 | 0.00 | 0.43 | 0.47 | 0.56 |
| EigenTHREADER+$r_c$ | | 0.00 | 0.32 | 0.37 | 0.38 | 0.00 | 0.39 | 0.43 | 0.47 | 0.00 | 0.43 | 0.47 | 0.50 | 0.00 | 0.54 | 0.57 | 0.59 |
| Map_Align | | 0.00 | 0.21 | 0.27 | 0.26 | 0.00 | 0.25 | 0.30 | 0.26 | 0.00 | 0.25 | 0.30 | 0.32 | 0.00 | 0.32 | 0.33 | 0.38 |
| Map_Align+$r_c$ | | 0.00 | **0.43** | **0.57** | **0.65** | 0.00 | **0.54** | **0.60** | **0.68** | 0.00 | **0.64** | **0.67** | **0.74** | 0.00 | **0.71** | **0.73** | **0.74** |
| AlEigen | CASP13 | 0.00 | 0.25 | 0.32 | 0.33 | 0.00 | 0.30 | 0.36 | 0.46 | 0.00 | 0.35 | 0.41 | 0.50 | 0.00 | 0.35 | 0.41 | 0.50 |
| AlEigen+$r_c$ | | 0.50 | 0.40 | 0.41 | 0.50 | 0.50 | 0.55 | 0.55 | 0.62 | 0.50 | 0.65 | 0.64 | 0.62 | 0.50 | 0.65 | 0.64 | 0.67 |
| EigenTHREADER | | 0.00 | 0.35 | 0.41 | 0.46 | 0.00 | 0.45 | 0.45 | 0.46 | 0.00 | 0.45 | 0.45 | 0.46 | 0.00 | 0.50 | 0.50 | 0.54 |
| EigenTHREADER+$r_c$ | | 0.00 | 0.45 | 0.45 | 0.58 | 0.00 | **0.70** | **0.68** | 0.67 | 0.00 | **0.70** | **0.73** | **0.71** | 0.00 | **0.75** | **0.77** | **0.75** |
| Map_Align | | 0.50 | 0.40 | 0.45 | 0.54 | 0.50 | 0.45 | 0.50 | 0.58 | 0.50 | 0.50 | 0.55 | 0.62 | 0.50 | 0.50 | 0.55 | 0.62 |
| Map_Align+$r_c$ | | **1.00** | **0.55** | **0.55** | **0.62** | **1.00** | **0.70** | **0.68** | **0.71** | **1.00** | **0.70** | **0.73** | **0.71** | **1.00** | **0.75** | **0.77** | **0.75** |

Table 8: **Fold recognition performances with predicted contacts from top-15 contact predictors**. True Positive Rate (TPR) fold recognition performances on CASP12 and CASP13 benchmark sets. The TPR performances are assessed with respect to the **top-1**, **top-5**, **top10** and **top-20** ranked hits. ECOD hierarchy: (F) Family Level (4 targets in CASP12, 2 targets in CASP13), (T) Topology Level (28 targets in CASP12, 20 targets in CASP13), (H) Homology Level (30 targets in CASP12, 22 targets in CASP13), (X) Possible Homology Level (34 targets in CASP12, 24 targets in CASP13). EigenTHREADER, Map_Align and AlEigen use their own scoring system EigenTHREADER+$r_c$, Map_Align+$r_c$ and AlEigen+$r_c$ use statistically significant congruence coefficient. Best TPR performances per column on CASP12 and CASP13 benchmark sets are highlighted in bold.

| | | top-1 hit | | | | top-5 hits | | | | top-10 hits | | | | top-20 hits | | | |
|---|---|---|---|---|---|---|---|---|---|---|---|---|---|---|---|---|---|
| Method | Dataset | F | T | H | X | F | T | H | X | F | T | H | X | F | T | H | X |
| AlEigen | CASP12 | 0.00 | 0.18 | 0.23 | 0.24 | 0.00 | 0.18 | 0.23 | 0.26 | 0.00 | 0.18 | 0.27 | 0.29 | 0.00 | 0.29 | 0.33 | 0.32 |
| AlEigen+$r_c$ | | **0.25** | 0.25 | 0.33 | 0.38 | **0.25** | 0.43 | 0.47 | 0.50 | **0.25** | 0.43 | 0.50 | 0.56 | **0.25** | 0.57 | 0.60 | 0.62 |
| EigenTHREADER | | 0.00 | 0.18 | 0.27 | 0.29 | 0.00 | 0.32 | 0.37 | 0.47 | 0.00 | 0.39 | 0.43 | 0.53 | 0.00 | 0.43 | 0.47 | 0.59 |
| EigenTHREADER+$r_c$ | | **0.25** | 0.29 | 0.37 | 0.38 | **0.25** | 0.50 | 0.53 | 0.59 | **0.25** | 0.50 | 0.53 | 0.62 | **0.25** | 0.57 | 0.60 | 0.65 |
| Map_Align | | 0.00 | 0.21 | 0.27 | 0.26 | 0.00 | **0.25** | 0.30 | 0.26 | 0.00 | **0.25** | 0.30 | 0.32 | 0.00 | 0.32 | 0.33 | 0.38 |
| Map_Align+$r_c$ | | **0.25** | **0.50** | **0.57** | **0.62** | **0.25** | **0.64** | **0.67** | **0.68** | **0.25** | **0.71** | **0.73** | **0.74** | **0.25** | **0.75** | **0.77** | **0.76** |
| AlEigen | CASP13 | 0.00 | 0.25 | 0.32 | 0.29 | 0.00 | 0.30 | 0.36 | 0.42 | 0.00 | 0.30 | 0.36 | 0.42 | 0.00 | 0.35 | 0.41 | 0.46 |
| AlEigen+$r_c$ | | **0.50** | 0.45 | 0.45 | 0.54 | 0.50 | 0.55 | 0.55 | 0.67 | 0.50 | 0.65 | 0.64 | 0.67 | 0.50 | **0.75** | 0.73 | 0.75 |
| EigenTHREADER | | 0.00 | 0.35 | 0.41 | 0.46 | 0.00 | 0.45 | 0.45 | 0.46 | 0.00 | 0.45 | 0.45 | 0.46 | 0.00 | 0.45 | 0.45 | 0.54 |
| EigenTHREADER+$r_c$ | | 0.00 | 0.45 | 0.45 | **0.58** | 0.00 | 0.65 | 0.64 | **0.71** | 0.00 | **0.75** | **0.73** | **0.75** | 0.00 | **0.75** | **0.77** | **0.79** |
| Map_Align | | **0.50** | 0.35 | 0.41 | 0.50 | 0.50 | 0.40 | 0.45 | 0.54 | 0.50 | 0.45 | 0.50 | 0.58 | 0.50 | 0.45 | 0.50 | 0.58 |
| Map_Align+$r_c$ | | **0.50** | **0.55** | **0.55** | **0.58** | **1.00** | **0.70** | **0.68** | 0.67 | **1.00** | 0.70 | 0.68 | 0.71 | **1.00** | **0.75** | **0.77** | **0.79** |

Table 9: **Fold recognition performances with predicted contacts from top-20 contact predictors**. True Positive Rate (TPR) fold recognition performances on CASP12 and CASP13 benchmark sets. The TPR performances are assessed with respect to the **top-1**, **top-5**, **top10** and **top-20** ranked hits. ECOD hierarchy: (F) Family Level (4 targets in CASP12, 2 targets in CASP13), (T) Topology Level (28 targets in CASP12, 20 targets in CASP13), (H) Homology Level (30 targets in CASP12, 22 targets in CASP13), (X) Possible Homology Level (34 targets in CASP12, 24 targets in CASP13). EigenTHREADER, Map_Align and AlEigen use their own scoring system EigenTHREADER+$r_c$, Map_Align+$r_c$ and AlEigen+$r_c$ use statistically significant congruence coefficient. Best TPR performances per column on CASP12 and CASP13 benchmark sets are highlighted in bold.

##### 7.4.2 Fold recognition with predicted distances/structures

As we have done with predicted contacts, we search all structure predictions submitted at CASP for a single target against the ECOD templates. The database search has been performed with two structural alignment tools, CE and TM-Align. Also in this case, for each group we consider only the first submitted prediction. Distance maps have been recovered from the predicted structures and distance map alignments from the structural alignments returned by CE and TM-Align. Also here, we compare the folding recognition capabilities of CE and TM-Align with their own scoring schemes against those obtained by using the congruence coefficient, CE+$r_c$ and TM-Align+$r_c$, respectively. In order to give more weight to short distances, we transformed the distance maps by taking the reciprocal of each distance (distances equal to 0 remain unchanged). Such transformation basically converts distance maps as *weighted* contact map. Preliminary tests showed that on the average such transformation provides slightly better performances than by using the real distances (data not shown). The true positive rate performances on CASP12 and CASP13 benchmark sets are summarized in Table 10. The TPR performances restricted to FM (Free Modelling) targets only are shown in Table 11. In Tables 12-15 we show fold recognition performances by restricting contact predictions to the top-5, top-10, top15 and top-20 best performing strucutre predictors at CASP12 and CASP13 (obtained by the official CASP rankings).

In these tests we cannot detect any approach that is overall better than all the others in all cases. We point out, however, that in Table 11, TM-align is the only method that can correctly identify the correct template at the family level for one FM CASP12 target. The restriction to the best performing predictors in Tables 12-15 gives better fold recognition accuracies for the CASP12 benchmark set but not for CASP13.

| | | top-1 hit | | | | top-5 hits | | | | top-10 hits | | | | top-20 hits | | | |
|---|---|---|---|---|---|---|---|---|---|---|---|---|---|---|---|---|---|
| Method | Benchmark set | F | T | H | X | F | T | H | X | F | T | H | X | F | T | H | X |
| CE | CASP12 | 0.00 | 0.39 | 0.43 | **0.41** | 0.00 | **0.57** | **0.63** | **0.62** | 0.00 | 0.57 | **0.63** | **0.62** | 0.00 | 0.57 | **0.63** | **0.62** |
| CE+$r_c$ | | 0.00 | **0.43** | **0.47** | **0.41** | 0.00 | 0.54 | 0.57 | 0.56 | 0.00 | 0.57 | 0.60 | 0.59 | 0.00 | 0.61 | **0.63** | **0.62** |
| TM-Align | | 0.00 | 0.29 | 0.37 | 0.38 | 0.00 | 0.50 | 0.53 | 0.56 | 0.00 | 0.57 | 0.60 | 0.59 | **0.50** | **0.64** | **0.63** | **0.62** |
| TM-Align+$r_c$ | | 0.00 | 0.39 | **0.47** | **0.41** | **0.25** | 0.50 | 0.57 | 0.53 | **0.25** | **0.61** | **0.63** | 0.59 | **0.50** | 0.61 | **0.63** | 0.59 |
| CE | CASP13 | **0.50** | **0.60** | **0.64** | **0.67** | **0.50** | 0.60 | 0.64 | 0.67 | **0.50** | 0.60 | 0.64 | 0.71 | **0.50** | 0.60 | 0.64 | 0.71 |
| CE+$r_c$ | | **0.50** | 0.50 | 0.50 | 0.54 | **0.50** | 0.65 | 0.68 | 0.67 | **0.50** | 0.65 | 0.68 | 0.71 | **0.50** | 0.70 | 0.73 | 0.79 |
| TM-Align | | 0.00 | **0.60** | **0.64** | **0.67** | **0.50** | 0.60 | 0.64 | **0.71** | **0.50** | 0.60 | 0.68 | **0.75** | **0.50** | 0.60 | 0.68 | 0.75 |
| TM-Align+$r_c$ | | **0.50** | 0.50 | 0.50 | 0.54 | **0.50** | **0.70** | **0.73** | **0.71** | **0.50** | **0.70** | **0.73** | **0.75** | **0.50** | **0.85** | **0.86** | **0.83** |

Table 10: **Fold recognition performances with predicted distances/structures**. True Positive Rate (TPR) fold recognition performances on CASP12 and CASP13 benchmark sets. The TPR performances are assessed with respect to the **top-1**, **top-5**, **top10** and **top-20** ranked hits. ECOD hierarchy: (F) Family Level (2 targets in CASP12, 0 targets in CASP13), (T) Topology Level (7 targets in both CASP12 and CASP13), (H) Homology Level (8 targets in both CASP12 and CASP13), (X) Possible Homology Level (12 targets in CASP12, 9 targets in CASP13). CE and TM-Align use their own scoring system. CE+$r_c$, and TM-Align+$r_c$ use statistically significant congruence coefficient. Best TPR performances per column on CASP12 and CASP13 benchmark sets are highlighted in bold.

| Method | Benchmark set | top-1 hit | | | | top-5 hits | | | | top-10 hits | | | | top-20 hits | | | |
|---|---|---|---|---|---|---|---|---|---|---|---|---|---|---|---|---|---|
| | | F | T | H | X | F | T | H | X | F | T | H | X | F | T | H | X |
| CE | CASP12 | 0.00 | 0.00 | 0.00 | 0.00 | 0.00 | **0.29** | **0.38** | **0.33** | 0.00 | **0.29** | **0.38** | **0.33** | 0.00 | **0.29** | **0.38** | 0.33 |
| CE+$r_c$ | | 0.00 | **0.14** | 0.12 | 0.08 | 0.00 | **0.29** | **0.38** | **0.33** | 0.00 | **0.29** | **0.38** | **0.33** | 0.00 | **0.29** | **0.38** | **0.42** |
| TM-Align | | 0.00 | **0.14** | **0.25** | **0.17** | 0.00 | **0.29** | **0.38** | **0.33** | 0.00 | **0.29** | **0.38** | **0.33** | **0.50** | **0.29** | **0.38** | 0.33 |
| TM-Align+$r_c$ | | 0.00 | **0.14** | 0.12 | 0.08 | 0.00 | 0.14 | 0.25 | 0.25 | 0.00 | **0.29** | **0.38** | **0.33** | 0.00 | **0.29** | **0.38** | 0.33 |
| CE | | 0.00 | **0.43** | **0.50** | **0.44** | 0.00 | **0.43** | **0.50** | 0.44 | 0.00 | **0.43** | **0.50** | **0.56** | 0.00 | **0.43** | **0.50** | **0.56** |
| CE+$r_c$ | | 0.00 | 0.29 | 0.25 | 0.22 | 0.00 | **0.43** | **0.50** | 0.44 | 0.00 | **0.43** | **0.50** | **0.56** | 0.00 | **0.43** | **0.50** | **0.56** |
| TM-Align | | 0.00 | **0.43** | **0.50** | **0.44** | 0.00 | **0.43** | **0.50** | **0.56** | 0.00 | **0.43** | **0.50** | **0.56** | 0.00 | **0.43** | **0.50** | **0.56** |
| TM-Align+$r_c$ | | 0.00 | 0.43 | 0.38 | 0.33 | 0.00 | **0.43** | **0.50** | 0.44 | 0.00 | **0.43** | **0.50** | **0.56** | 0.00 | **0.43** | **0.50** | **0.56** |

Table 11: **Fold recognition performances on FM targets with predicted distances/structures**. True Positive Rate (TPR) fold recognition performances on CASP12 and CASP13 benchmark sets. The TPR performances are assessed with respect to the **top-1**, **top-5**, **top10** and **top-20** ranked hits. ECOD hierarchy: (F) Family Level (4 targets in CASP12, 2 targets in CASP13), (T) Topology Level (28 targets in CASP12, 20 targets in CASP13), (H) Homology Level (30 targets in CASP12, 22 targets in CASP13), (X) Possible Homology Level (34 targets in CASP12, 24 targets in CASP13). CE and TM-Align use their own scoring system. CE+$r_c$, and TM-Align+$r_c$ use statistically significant congruence coefficient. Best TPR performances per column on CASP12 and CASP13 benchmark sets are highlighted in bold.

| Method | Benchmark set | top-1 hit | | | | top-5 hits | | | | top-10 hits | | | | top-20 hits | | | |
|---|---|---|---|---|---|---|---|---|---|---|---|---|---|---|---|---|---|
| | | F | T | H | X | F | T | H | X | F | T | H | X | F | T | H | X |
| CE | CASP12 | 0.00 | **0.21** | **0.23** | **0.24** | 0.00 | **0.25** | **0.27** | **0.26** | 0.00 | **0.25** | **0.27** | **0.29** | 0.00 | **0.29** | **0.30** | **0.29** |
| CE+$r_c$ | | 0.00 | 0.18 | 0.17 | 0.18 | 0.00 | 0.21 | 0.23 | 0.24 | 0.00 | **0.25** | **0.27** | 0.26 | 0.00 | 0.25 | 0.27 | 0.26 |
| TM-Align | | 0.00 | 0.14 | 0.13 | 0.15 | 0.00 | 0.21 | 0.20 | 0.21 | 0.00 | 0.21 | 0.20 | 0.21 | 0.00 | 0.21 | 0.20 | 0.21 |
| TM-Align+$r_c$ | | 0.00 | 0.18 | 0.17 | 0.18 | 0.00 | 0.18 | 0.17 | 0.18 | 0.00 | 0.18 | 0.20 | 0.21 | 0.00 | 0.25 | 0.27 | 0.26 |
| CE | CASP13 | **0.50** | 0.50 | 0.50 | 0.62 | **0.50** | 0.60 | 0.64 | **0.71** | **0.50** | 0.60 | 0.64 | **0.79** | **1.00** | 0.60 | 0.64 | **0.79** |
| CE+$r_c$ | | **0.50** | 0.50 | 0.50 | 0.62 | **0.50** | 0.60 | 0.59 | 0.67 | **0.50** | **0.70** | **0.68** | **0.79** | 0.50 | **0.80** | **0.77** | **0.79** |
| TM-Align | | 0.00 | **0.55** | **0.64** | **0.67** | **0.50** | 0.60 | **0.68** | **0.71** | **0.50** | 0.60 | **0.68** | 0.71 | 0.50 | 0.60 | 0.68 | 0.71 |
| TM-Align+$r_c$ | | 0.50 | 0.55 | 0.55 | 0.58 | **0.50** | **0.65** | 0.64 | 0.67 | **0.50** | 0.65 | 0.64 | 0.67 | 0.50 | 0.70 | 0.68 | 0.71 |

Table 12: **Fold recognition performances with predicted distances/structures from top-5 structure predictors**. True Positive Rate (TPR) fold recognition performances on CASP12 and CASP13 benchmark sets. The TPR performances are assessed with respect to the **top-1**, **top-5**, **top10** and **top-20** ranked hits. ECOD hierarchy: (F) Family Level (4 targets in CASP12, 2 targets in CASP13), (T) Topology Level (28 targets in CASP12, 20 targets in CASP13), (H) Homology Level (30 targets in CASP12, 22 targets in CASP13), (X) Possible Homology Level (34 targets in CASP12, 24 targets in CASP13). CE and TM-Align use their own scoring system. CE+$r_c$, and TM-Align+$r_c$ use statistically significant congruence coefficient. Best TPR performances per column on CASP12 and CASP13 benchmark sets are highlighted in bold.

| | | top-1 hit | | | | top-5 hits | | | | top-10 hits | | | | top-20 hits | | | |
|---|---|---|---|---|---|---|---|---|---|---|---|---|---|---|---|---|---|
| Method | Benchmark set | F | T | H | X | F | T | H | X | F | T | H | X | F | T | H | X |
| CE | CASP12 | **0.25** | **0.61** | **0.63** | **0.65** | **0.25** | **0.68** | **0.70** | **0.68** | **0.50** | **0.68** | **0.70** | **0.68** | **0.50** | **0.68** | **0.70** | **0.71** |
| CE+$r_c$ | | 0.00 | 0.50 | 0.50 | 0.59 | 0.00 | 0.61 | 0.63 | 0.62 | 0.00 | 0.61 | 0.63 | 0.62 | 0.00 | **0.68** | **0.70** | 0.68 |
| TM-Align | | 0.00 | 0.54 | 0.53 | 0.50 | 0.00 | 0.61 | 0.60 | 0.59 | 0.00 | 0.61 | 0.60 | 0.59 | 0.00 | 0.61 | 0.60 | 0.59 |
| TM-Align+$r_c$ | | 0.00 | 0.57 | 0.60 | 0.56 | **0.25** | 0.64 | 0.63 | 0.59 | 0.25 | **0.68** | 0.67 | 0.62 | 0.25 | **0.68** | **0.70** | 0.68 |
| CE | CASP13 | **0.50** | 0.55 | 0.59 | **0.67** | **0.50** | 0.60 | 0.64 | 0.71 | **0.50** | 0.65 | 0.68 | **0.75** | **1.00** | 0.75 | 0.77 | **0.83** |
| CE+$r_c$ | | **0.50** | 0.55 | 0.55 | 0.62 | **0.50** | 0.70 | 0.68 | **0.75** | **0.50** | 0.70 | 0.73 | 0.83 | 0.50 | **0.80** | **0.82** | **0.83** |
| TM-Align | | 0.00 | 0.60 | **0.64** | **0.67** | **0.50** | 0.60 | 0.68 | 0.71 | **0.50** | 0.60 | 0.68 | 0.71 | 0.50 | 0.60 | 0.68 | 0.71 |
| TM-Align+$r_c$ | | **0.50** | **0.60** | 0.59 | **0.67** | **0.50** | **0.75** | **0.77** | **0.75** | **0.50** | **0.75** | **0.77** | **0.75** | 0.50 | **0.80** | **0.82** | 0.75 |

Table 13: **Fold recognition performances with predicted distances/structures from top-10 structure predictors**. True Positive Rate (TPR) fold recognition performances on CASP12 and CASP13 benchmark sets. The TPR performances are assessed with respect to the **top-1**, **top-5**, **top10** and **top-20** ranked hits. ECOD hierarchy: (F) Family Level (4 targets in CASP12, 2 targets in CASP13), (T) Topology Level (28 targets in CASP12, 20 targets in CASP13), (H) Homology Level (30 targets in CASP12, 22 targets in CASP13), (X) Possible Homology Level (34 targets in CASP12, 24 targets in CASP13). CE and TM-Align use their own scoring system. CE+$r_c$, and TM-Align+$r_c$ use statistically significant congruence coefficient. Best TPR performances per column on CASP12 and CASP13 benchmark sets are highlighted in bold.

| | | top-1 hit | | | | top-5 hits | | | | top-10 hits | | | | top-20 hits | | | |
|---|---|---|---|---|---|---|---|---|---|---|---|---|---|---|---|---|---|
| Method | Benchmark set | F | T | H | X | F | T | H | X | F | T | H | X | F | T | H | X |
| CE | CASP12 | 0.00 | **0.64** | **0.67** | **0.65** | **0.25** | **0.68** | **0.70** | **0.68** | **0.25** | **0.68** | **0.70** | **0.71** | **0.50** | **0.71** | **0.73** | **0.74** |
| CE+$r_c$ | | 0.00 | 0.54 | 0.57 | 0.59 | 0.00 | 0.61 | 0.63 | 0.62 | 0.00 | 0.61 | 0.63 | 0.62 | 0.00 | 0.64 | 0.70 | 0.68 |
| TM-Align | | 0.00 | 0.54 | 0.53 | 0.50 | 0.00 | 0.64 | 0.60 | 0.59 | 0.00 | 0.64 | 0.60 | 0.59 | 0.00 | 0.64 | 0.60 | 0.59 |
| TM-Align+$r_c$ | | 0.00 | 0.57 | 0.60 | 0.56 | 0.00 | 0.57 | 0.60 | 0.56 | **0.25** | **0.68** | 0.67 | 0.62 | 0.25 | 0.68 | 0.70 | 0.65 |
| CE | CASP13 | **0.50** | 0.50 | 0.55 | 0.62 | **0.50** | 0.55 | 0.59 | 0.67 | **0.50** | 0.60 | 0.64 | 0.71 | **1.00** | 0.70 | 0.73 | **0.83** |
| CE+$r_c$ | | **0.50** | 0.55 | 0.55 | 0.62 | **0.50** | 0.70 | 0.68 | **0.75** | **0.50** | 0.70 | 0.73 | **0.83** | 0.50 | **0.75** | **0.77** | **0.83** |
| TM-Align | | 0.00 | **0.60** | **0.64** | **0.67** | **0.50** | 0.60 | 0.68 | 0.71 | **0.50** | 0.60 | 0.68 | 0.71 | 0.50 | 0.60 | 0.68 | 0.71 |
| TM-Align+$r_c$ | | **0.50** | **0.60** | 0.59 | **0.67** | **0.50** | **0.75** | **0.73** | 0.71 | **0.50** | **0.75** | **0.77** | 0.75 | 0.50 | **0.75** | **0.77** | 0.75 |

Table 14: **Fold recognition performances with predicted distances/structures from top-15 structure predictors**. True Positive Rate (TPR) fold recognition performances on CASP12 and CASP13 benchmark sets. The TPR performances are assessed with respect to the **top-1**, **top-5**, **top10** and **top-20** ranked hits. ECOD hierarchy: (F) Family Level (4 targets in CASP12, 2 targets in CASP13), (T) Topology Level (28 targets in CASP12, 20 targets in CASP13), (H) Homology Level (30 targets in CASP12, 22 targets in CASP13), (X) Possible Homology Level (34 targets in CASP12, 24 targets in CASP13). CE and TM-Align use their own scoring system. CE+$r_c$, and TM-Align+$r_c$ use statistically significant congruence coefficient. Best TPR performances per column on CASP12 and CASP13 benchmark sets are highlighted in bold.

| Method | Benchmark set | top-1 hit | | | | top-5 hits | | | | top-10 hits | | | | top-20 hits | | | |
|---|---|---|---|---|---|---|---|---|---|---|---|---|---|---|---|---|---|
| | | F | T | H | X | F | T | H | X | F | T | H | X | F | T | H | X |
| CE | CASP12 | 0.00 | **0.64** | **0.67** | **0.65** | **0.25** | **0.68** | **0.70** | **0.68** | **0.25** | **0.71** | **0.73** | **0.71** | **0.50** | **0.71** | **0.73** | **0.74** |
| CE+$r_c$ | | 0.00 | 0.46 | 0.50 | 0.56 | 0.00 | 0.57 | 0.57 | 0.56 | 0.00 | 0.57 | 0.57 | 0.56 | 0.00 | 0.64 | 0.70 | 0.68 |
| TM-Align | | 0.00 | 0.57 | 0.57 | 0.53 | 0.00 | 0.64 | 0.60 | 0.59 | 0.00 | 0.64 | 0.60 | 0.59 | 0.00 | 0.64 | 0.60 | 0.59 |
| TM-Align+$r_c$ | | 0.00 | 0.54 | 0.57 | 0.53 | 0.00 | 0.54 | 0.57 | 0.53 | **0.25** | 0.68 | 0.67 | 0.62 | 0.25 | 0.68 | 0.70 | 0.65 |
| CE | CASP13 | **0.50** | 0.55 | 0.59 | **0.67** | **0.50** | 0.60 | 0.64 | 0.71 | **0.50** | 0.65 | 0.68 | 0.75 | **0.50** | 0.70 | 0.73 | 0.83 |
| CE+$r_c$ | | **0.50** | **0.65** | **0.64** | **0.67** | **0.50** | **0.75** | **0.73** | **0.79** | **0.50** | **0.75** | **0.77** | **0.88** | **0.50** | **0.80** | **0.82** | **0.88** |
| TM-Align | | 0.00 | 0.60 | **0.64** | **0.67** | **0.50** | 0.60 | 0.68 | 0.71 | **0.50** | 0.60 | 0.68 | 0.71 | **0.50** | 0.60 | 0.68 | 0.71 |
| TM-Align+$r_c$ | | **0.50** | 0.60 | 0.59 | **0.67** | **0.50** | **0.75** | **0.73** | 0.71 | **0.50** | **0.75** | **0.77** | 0.75 | **0.50** | 0.75 | 0.77 | 0.75 |

Table 15: **Fold recognition performances with predicted distances/structures from top-20 structure predictors**. True Positive Rate (TPR) fold recognition performances on CASP12 and CASP13 benchmark sets. The TPR performances are assessed with respect to the **top-1**, **top-5**, **top10** and **top-20** ranked hits. ECOD hierarchy: (F) Family Level (4 targets in CASP12, 2 targets in CASP13), (T) Topology Level (28 targets in CASP12, 20 targets in CASP13), (H) Homology Level (30 targets in CASP12, 22 targets in CASP13), (X) Possible Homology Level (34 targets in CASP12, 24 targets in CASP13). CE and TM-Align use their own scoring system. CE+$r_c$, and TM-Align+$r_c$ use statistically significant congruence coefficient. Best TPR performances per column on CASP12 and CASP13 benchmark sets are highlighted in bold.

##### 7.4.3 The impact of detecting statistical significant similarities

The possibility to detect statistically significant similarities between target and template maps has some impact in fold recognition performances. In table 16 we show how the fold recognition performances with the congruence coefficient as fitness function change if we remove the statistical similarity filter, i.e. if we include in fold recognition evaluation also those templates that do not show statistically significantly similarity with the target map (with respect to some given alignment). In table 16, performances that include non-statistically significant comparisons are indicated with $^*$. It is clear that the statistically significant filtering has a relevant impact in fold recognition performances with predicted contacts. Conversely, when dealing with distance maps, the degree of freedom of the P-value in Equation is usually quite large, thus most of the alignments between two maps are evaluated as statistically significant. Thus, in the fold recognition tests with distance maps the statistically significant filtering has no effect and there is no change in fold recognition performances.

AlEigen, EigenTHREADER and Map_Align do not evaluate whether two aligned maps are significantly similar, thus their template ranking include also uninteresting templates that can affect fold recognition performances. In Table 17 we show how the True Positive Rate performances change if we exclude (i.e. filter out) from the evaluation those templates that do not show statistically significantly similarity with the target map (with respect to the computed alignment). In Table 17, where we use the $^*$ mark to indicate performances that rank only statistically significant comparisons, we can notice that EigenTHREADER performances are overall improved by such filtering. The performances of Map_Align are also improved in some cases, while AlEigen is unaffected.

| | | top-1 hit | | | | top-5 hits | | | | top-10 hits | | | | top-20 hits | | | |
|---|---|---|---|---|---|---|---|---|---|---|---|---|---|---|---|---|---|
| Method | Dataset | F | T | H | X | F | T | H | X | F | T | H | X | F | T | H | X |
| AlEigen+$r_c$ | CASP12 | 0.25 | **0.29** | **0.37** | **0.38** | 0.25 | **0.39** | **0.43** | **0.47** | 0.25 | 0.39 | **0.47** | **0.56** | 0.25 | **0.54** | **0.57** | **0.62** |
| AlEigen+$r_c^*$ | | 0.25 | 0.25 | 0.30 | 0.35 | 0.25 | 0.32 | 0.37 | 0.41 | 0.25 | 0.39 | 0.43 | 0.47 | 0.25 | 0.46 | 0.50 | 0.53 |
| EigenTHREADER+$r_c$ | CASP12 | 0.25 | **0.32** | 0.40 | 0.41 | 0.25 | **0.54** | **0.57** | **0.62** | 0.25 | **0.57** | **0.60** | **0.68** | 0.25 | **0.61** | **0.63** | **0.68** |
| EigenTHREADER+$r_c^*$ | | 0.25 | 0.29 | 0.40 | 0.41 | 0.25 | 0.43 | 0.50 | 0.56 | 0.25 | 0.46 | 0.50 | 0.56 | 0.25 | 0.50 | 0.53 | 0.62 |
| Map_Align+$r_c$ | CASP12 | 0.25 | 0.57 | 0.60 | **0.65** | 0.25 | 0.61 | 0.63 | **0.65** | 0.25 | **0.68** | **0.70** | **0.71** | 0.25 | **0.71** | **0.73** | **0.74** |
| Map_Align+$r_c^*$ | | 0.25 | 0.57 | 0.60 | 0.62 | 0.25 | 0.61 | 0.63 | 0.62 | 0.25 | 0.61 | 0.63 | 0.62 | 0.25 | 0.64 | 0.67 | 0.65 |
| AlEigen+$r_c$ | CASP13 | 0.00 | **0.35** | **0.36** | **0.46** | 0.00 | **0.45** | **0.45** | **0.62** | 0.00 | **0.60** | **0.59** | **0.67** | **0.50** | **0.70** | **0.73** | **0.79** |
| AlEigen+$r_c^*$ | | 0.00 | 0.30 | 0.32 | 0.42 | 0.00 | 0.40 | 0.41 | 0.58 | 0.00 | 0.50 | 0.50 | 0.58 | 0.00 | 0.55 | 0.55 | 0.62 |
| EigenTHREADER+$r_c$ | CASP13 | 0.00 | **0.50** | **0.50** | **0.58** | 0.00 | **0.70** | **0.68** | **0.71** | 0.00 | **0.80** | **0.77** | **0.79** | 0.00 | **0.80** | **0.77** | **0.83** |
| EigenTHREADER+$r_c^*$ | | 0.00 | 0.35 | 0.36 | 0.46 | 0.00 | 0.65 | 0.64 | 0.67 | 0.00 | 0.70 | 0.68 | 0.67 | 0.00 | 0.70 | 0.68 | 0.75 |
| Map_Align+$r_c$ | CASP13 | 0.50 | **0.60** | **0.59** | **0.62** | 1.00 | 0.75 | 0.73 | 0.71 | 1.00 | **0.85** | **0.82** | **0.83** | 1.00 | **0.85** | **0.86** | **0.83** |
| Map_Align+$r_c^*$ | | 0.50 | 0.55 | 0.55 | 0.58 | 1.00 | 0.75 | 0.73 | 0.71 | 1.00 | 0.75 | 0.73 | 0.75 | 1.00 | 0.75 | 0.77 | 0.75 |

Table 16: **Fold recognition performances with predicted contacts from all contact predictors**. True Positive Rate (TPR) fold recognition performances on CASP12 and CASP13 benchmark sets. The TPR performances are assessed with respect to the **top-1**, **top-5**, **top10** and **top-20** ranked hits. ECOD hierarchy: (F) Family Level (4 targets in CASP12, 2 targets in CASP13), (T) Topology Level (28 targets in CASP12, 20 targets in CASP13), (H) Homology Level (30 targets in CASP12, 22 targets in CASP13), (X) Possible Homology Level (34 targets in CASP12, 24 targets in CASP13). EigenTHREADER+$r_c^*$, Map_Align+$r_c^*$ and AlEigen+$r_c^*$ indicate that non-significant similarities between target and template maps have been considered for evaluating the TPR performances. The changes between original and unfiltered performances are highlighted in bold.

| | | top-1 hit | | | | top-5 hits | | | | top-10 hits | | | | top-20 hits | | | |
|---|---|---|---|---|---|---|---|---|---|---|---|---|---|---|---|---|---|
| Method | Dataset | F | T | H | X | F | T | H | X | F | T | H | X | F | T | H | X |
| AlEigen | CASP12 | 0.00 | 0.07 | 0.07 | 0.06 | 0.00 | 0.07 | 0.07 | 0.09 | 0.00 | 0.07 | 0.10 | 0.12 | 0.00 | 0.07 | 0.10 | 0.15 |
| AlEigen* | | 0.00 | 0.07 | 0.07 | 0.06 | 0.00 | 0.07 | 0.07 | 0.09 | 0.00 | 0.07 | 0.10 | 0.12 | 0.00 | 0.07 | 0.10 | 0.15 |
| EigenTHREADER | CASP12 | 0.00 | 0.00 | 0.00 | 0.00 | 0.00 | 0.07 | 0.07 | 0.06 | 0.00 | 0.07 | 0.07 | 0.09 | 0.00 | 0.11 | 0.10 | 0.15 |
| EigenTHREADER* | | 0.00 | 0.00 | 0.00 | **0.03** | 0.00 | 0.07 | 0.07 | **0.15** | 0.00 | **0.29** | **0.27** | **0.29** | 0.00 | **0.32** | **0.33** | **0.38** |
| Map_Align | CASP12 | 0.00 | 0.07 | 0.07 | 0.06 | 0.00 | 0.07 | 0.07 | 0.06 | 0.00 | 0.07 | 0.10 | 0.12 | 0.00 | 0.07 | 0.10 | 0.15 |
| Map_Align* | | 0.00 | 0.07 | 0.07 | 0.06 | 0.00 | 0.07 | 0.07 | 0.06 | 0.00 | 0.07 | 0.10 | 0.12 | 0.00 | 0.07 | 0.10 | 0.15 |
| AlEigen | CASP13 | 0.00 | 0.15 | 0.23 | 0.25 | 0.00 | 0.25 | 0.32 | 0.38 | 0.00 | 0.25 | 0.32 | 0.42 | 0.00 | 0.25 | 0.32 | 0.42 |
| AlEigen* | | 0.00 | 0.15 | 0.23 | 0.25 | 0.00 | 0.25 | 0.32 | 0.38 | 0.00 | 0.25 | 0.32 | 0.42 | 0.00 | 0.25 | 0.32 | 0.42 |
| EigenTHREADER | CASP13 | 0.00 | 0.10 | 0.09 | 0.12 | 0.00 | 0.15 | 0.18 | 0.25 | 0.00 | 0.15 | 0.18 | 0.29 | 0.00 | 0.20 | 0.23 | 0.33 |
| EigenTHREADER* | | 0.00 | **0.15** | **0.27** | **0.33** | 0.00 | **0.25** | **0.27** | **0.42** | 0.00 | **0.40** | **0.45** | **0.50** | 0.00 | **0.45** | **0.50** | **0.54** |
| Map_Align | CASP13 | 0.50 | 0.35 | 0.41 | 0.50 | 0.50 | 0.40 | 0.45 | 0.54 | 0.50 | 0.40 | 0.45 | 0.54 | 0.50 | 0.40 | 0.45 | 0.54 |
| Map_Align* | | 0.50 | 0.35 | 0.41 | 0.50 | 0.50 | 0.40 | 0.45 | 0.54 | 0.50 | **0.45** | **0.50** | **0.58** | 0.50 | **0.45** | **0.50** | **0.58** |

Table 17: **Fold recognition performances with predicted contacts from all contact predictors by filtering out non-significant similarities**. True Positive Rate (TPR) fold recognition performances on CASP12 and CASP13 benchmark sets. The TPR performances are assessed with respect to the **top-1**, **top-5**, **top10** and **top-20** ranked hits. ECOD hierarchy: (F) Family Level (4 targets in CASP12, 2 targets in CASP13), (T) Topology Level (28 targets in CASP12, 20 targets in CASP13), (H) Homology Level (30 targets in CASP12, 22 targets in CASP13), (X) Possible Homology Level (34 targets in CASP12, 24 targets in CASP13). EigenTHREADER*, Map_Align* and AlEigen* indicate that non-significant similarities between target and template maps have not been considered for evaluating the TPR performances. The changes between original and filtered performances are highlighted in bold.

### Appendix A Summations

#### Notation

- $X \in \mathbb{R}^{n \times n}$ is a symmetric matrix
- Squared matrix $X^2$

$$X^2_{ij} = (X_{ij})^2$$

- Matrix product $XY$

$$(XY)_{ij} = \sum_{k=0}^{n} X_{ik} Y_{kj}$$

- Sum of all the elements in a matrix:

$$\Sigma(X) = \sum_{i=1}^{n} \sum_{j=1}^{n} X_{ij}$$

- Diagonal of X in vectorial form:

$$D = [X_{11}, X_{22}, ...X_{nn}]$$

- Diagonal matrix from the main diagonal of X, $D_X$

$$D_X = DI$$

where $I \in \{0,1\}^{n \times n}$ is the identity matrix.

- Matrix $X$ with main diagonal set to zero, $\mathring{X}$

$$\mathring{X} = X - D_X$$

#### Summations over one index

1. $$\sum_{i=1}^{n} X_{ii} X_{ii} = tr(X^2) = tr(D_X D_X) = \Sigma(D_X D_X) = \Sigma((X - \mathring{X})(X - \mathring{X})) =$$
$$= \Sigma(XX) - 2\Sigma(X\mathring{X}) + \Sigma(\mathring{X}\mathring{X})$$

**Summations over two indexes**

2. $$\sum_{i=1}^{n}\sum_{j=1}^{n} X_{ii}X_{jj} = tr(X)^2 = \Big(\Sigma(X) - \Sigma(\mathring{X})\Big)^2 = $$
$$= \Sigma(X)^2 - 2\Sigma(X)\Sigma(\mathring{X}) + \Sigma(\mathring{X})^2$$

3. $$\sum_{i=1}^{n}\sum_{j=1}^{n} X_{ii}X_{ij} = \sum_{i=1}^{n}\sum_{j=1}^{n} X_{jj}X_{ij} = \Sigma(XD_X) = \Sigma(X(X - \mathring{X}))$$
$$= \Sigma(XX) - \Sigma(X\mathring{X})$$

4. $$\sum_{i=1}^{n}\sum_{j=1}^{n} X_{ij}X_{ij} = \sum_{i=1}^{n}\sum_{j=1}^{n} X_{ij}^2 = \Sigma(X^2)$$

5. $$\sum_{i=1}^{n}\sum_{i\neq j=1}^{n} X_{ii}X_{jj} = \sum_{i=1}^{n}\left(\sum_{j=1}^{n} X_{ii}X_{jj} - X_{ii}X_{ii}\right) =$$
$$= \sum_{i=1}^{n}\sum_{j=1}^{n} X_{ii}X_{jj} - \sum_{i=1}^{n} X_{ii}X_{ii} = \quad \text{(by 2,1)}$$
$$= tr(X)^2 - tr(X^2) =$$
$$= \Sigma(X)^2 - 2\Sigma(X)\Sigma(\mathring{X}) + \Sigma(\mathring{X})^2 - \Sigma(XX) + 2\Sigma(X\mathring{X}) - \Sigma(\mathring{X}\mathring{X})$$

6. $$\sum_{i=1}^{n}\sum_{i\neq j=1}^{n} X_{ij}X_{ii} = \sum_{i=1}^{n}\left(\sum_{j=1}^{n} X_{ij}X_{ii} - X_{ii}X_{ii}\right) = \sum_{i=1}^{n}\sum_{j=1}^{n} X_{ij}X_{ii} - \sum_{i=1}^{n} X_{ii}X_{ii} = \text{(by 3,1)}$$
$$= \Sigma(XX) - \Sigma(X\mathring{X}) - \Sigma(XX) + 2\Sigma(X\mathring{X}) - \Sigma(\mathring{X}\mathring{X}) =$$
$$= \Sigma(\mathring{X}X) - \Sigma(\mathring{X}\mathring{X})$$

7. $$\sum_{i=1}^{n}\sum_{i\neq j=1}^{n} X_{ij}X_{jj} = \sum_{i=1}^{n}\left(\sum_{j=1}^{n} X_{ij}X_{jj} - X_{ii}X_{ii}\right) = \sum_{i=1}^{n}\sum_{j=1}^{n} X_{ij}X_{jj} - \sum_{i=1}^{n} X_{ii}X_{ii} = \text{(by 3,1)}$$
$$= \Sigma(XX) - \Sigma(X\mathring{X}) - \Sigma(XX) + 2\Sigma(X\mathring{X}) - \Sigma(\mathring{X}\mathring{X}) =$$
$$= \Sigma(\mathring{X}X) - \Sigma(\mathring{X}\mathring{X})$$

$$8.\ \sum_{i=1}^{n}\sum_{i\neq j=1}^{n} X_{ij}X_{ji} = \sum_{i=1}^{n}\left(\sum_{j=1}^{n} X_{ij}X_{ji} - X_{ii}X_{ii}\right) = \sum_{i=1}^{n}\sum_{j=1}^{n} X_{ij}X_{ji} - \sum_{i=1}^{n} X_{ii}X_{ii} = \text{(by 4,1)}$$
$$= \Sigma(X^2) - tr(X^2) =$$
$$= \Sigma(\mathring{X}^2)$$

**Summations over three indexes**

$$9.\ \sum_{i=1}^{n}\sum_{j=1}^{n}\sum_{k=1}^{n} X_{ii}X_{jk} = \sum_{i=1}^{n} X_{ii}\left(\sum_{j=1}^{n}\sum_{k=1}^{n} X_{jk}\right) = tr(X)\Sigma(X) =$$
$$= \Sigma(X - \mathring{X})\Sigma(X) = (\Sigma(X) - \Sigma(\mathring{X}))\Sigma(X) =$$
$$= \Sigma(X)^2 - \Sigma(\mathring{X})\Sigma(X)$$

$$10.\ \sum_{i=1}^{n}\sum_{j=1}^{n}\sum_{k=1}^{n} X_{ij}X_{ik} = \sum_{j=1}^{n}\sum_{k=1}^{n}\left(\sum_{i=1}^{n} X_{ji}X_{ik}\right) = \Sigma(XX)$$

$$11.\ \sum_{i=1}^{n}\sum_{i\neq j=1}^{n}\sum_{k=1}^{n} X_{ij}X_{ik} = \sum_{i=1}^{n}\sum_{k=1}^{n}\left(\sum_{j=1}^{n} X_{ij}X_{ik} - X_{ii}X_{ik}\right) =$$
$$= \sum_{i=1}^{n}\sum_{j=1}^{n}\sum_{k=1}^{n} X_{ij}X_{ik} - \sum_{i=1}^{n}\sum_{k=1}^{n} X_{ii}X_{ik} = \quad \text{(by 10,3)}$$
$$= \Sigma(XX) - \Sigma(XX) + \Sigma(X\mathring{X}) =$$
$$= \Sigma(X\mathring{X})$$

$$12.\ \sum_{i=1}^{n}\sum_{i\neq j=1}^{n}\sum_{k=1}^{n} X_{ij}X_{jk} = \sum_{i=1}^{n}\sum_{k=1}^{n}\left(\sum_{j=1}^{n} X_{ij}X_{jk} - X_{ii}X_{ik}\right) =$$
$$= \sum_{i=1}^{n}\sum_{j=1}^{n}\sum_{k=1}^{n} X_{ij}X_{jk} - \sum_{i=1}^{n}\sum_{k=1}^{n} X_{ii}X_{ik} = \quad \text{(by 10,3)}$$
$$= \Sigma(XX) - \Sigma(XX) + \Sigma(X\mathring{X}) =$$
$$= \Sigma(X\mathring{X})$$

13. $$\sum_{i=1}^{n}\sum_{i\neq j=1}^{n}\sum_{k=1}^{n}X_{ij}X_{kk} = \sum_{i=1}^{n}\sum_{k=1}^{n}\left(\sum_{j=1}^{n}X_{ij}X_{kk} - X_{ii}X_{kk}\right) =$$
$$= \sum_{i=1}^{n}\sum_{j=1}^{n}\sum_{k=1}^{n}X_{ij}X_{kk} - \sum_{i=1}^{n}\sum_{k=1}^{n}X_{ii}X_{kk} = \quad \text{(by 9,2)}$$
$$= \Sigma(X)^2 - \Sigma(\mathring{X})\Sigma(X) - \Sigma(X)^2 + 2\Sigma(\mathring{X})\Sigma(X) - \Sigma(\mathring{X})^2 =$$
$$= \Sigma(\mathring{X})\Sigma(X) - \Sigma(\mathring{X})^2$$

14. $$\sum_{i=1}^{n}\sum_{i\neq j=1}^{n}\sum_{k=1}^{n}X_{ii}X_{jk} = \sum_{i=1}^{n}\sum_{k=1}^{n}\left(\sum_{j=1}^{n}X_{ii}X_{jk} - X_{ii}X_{ik}\right) =$$
$$= \sum_{i=1}^{n}\sum_{j=1}^{n}\sum_{k=1}^{n}X_{ii}X_{jk} - \sum_{i=1}^{n}\sum_{k=1}^{n}X_{ii}X_{ik} = \quad \text{(by 9,3)}$$
$$= \Sigma(X)^2 - \Sigma(\mathring{X})\Sigma(X) - \Sigma(XX) + \Sigma(X\mathring{X})$$

15. $$\sum_{i=1}^{n}\sum_{i\neq j=1}^{n}\sum_{i,j\neq k=1}^{n}X_{ij}X_{ki} = \sum_{i=1}^{n}\sum_{i\neq j=1}^{n}\left(\sum_{k=1}^{n}X_{ij}X_{ki} - X_{ij}X_{ii} - X_{ij}X_{ji}\right) =$$
$$= \sum_{i=1}^{n}\sum_{i\neq j=1}^{n}\sum_{k=1}^{n}X_{ij}X_{ki} - \sum_{i=1}^{n}\sum_{i\neq j=1}^{n}X_{ij}X_{ii} - \sum_{i=1}^{n}\sum_{i\neq j=1}^{n}X_{ij}X_{ji} = \quad \text{(by 11,6,8)}$$
$$= \Sigma(X\mathring{X}) - \Sigma(\mathring{X}X) + \Sigma(\mathring{X}\mathring{X}) - \Sigma(\mathring{X}^2) =$$
$$= \Sigma(\mathring{X}\mathring{X}) - \Sigma(\mathring{X}^2)$$

16. $$\sum_{i=1}^{n}\sum_{i\neq j=1}^{n}\sum_{i,j\neq k=1}^{n}X_{ij}X_{kj} = \sum_{i=1}^{n}\sum_{i\neq j=1}^{n}\left(\sum_{k=1}^{n}X_{ij}X_{kj} - X_{ij}X_{ij} - X_{ij}X_{jj}\right) =$$
$$= \sum_{i=1}^{n}\sum_{i\neq j=1}^{n}\sum_{k=1}^{n}X_{ij}X_{kj} - \sum_{i=1}^{n}\sum_{i\neq j=1}^{n}X_{ij}X_{ij} - \sum_{i=1}^{n}\sum_{i\neq j=1}^{n}X_{ij}X_{jj} = \quad \text{(by 12,8,7)}$$
$$= \Sigma(X\mathring{X}) - \Sigma(\mathring{X}^2) - \Sigma(\mathring{X}X) + \Sigma(\mathring{X}\mathring{X}) =$$
$$= \Sigma(\mathring{X}\mathring{X}) - \Sigma(\mathring{X}^2)$$

17. $$\sum_{i=1}^{n}\sum_{i\neq j=1}^{n}\sum_{i,j\neq k=1}^{n} X_{ij}X_{kk} = \sum_{i=1}^{n}\sum_{i\neq j=1}^{n}\left(\sum_{k=1}^{n} X_{ij}X_{kk} - X_{ij}X_{ii} - X_{ij}X_{jj}\right) =$$
$$= \sum_{i=1}^{n}\sum_{i\neq j=1}^{n}\sum_{k=1}^{n} X_{ij}X_{kk} - \sum_{i=1}^{n}\sum_{i\neq j=1}^{n} X_{ij}X_{ii} - \sum_{i=1}^{n}\sum_{i\neq j=1}^{n} X_{ij}X_{jj} = \quad \text{(by 13,6,7)}$$
$$= \Sigma(\mathring{X})\Sigma(X) - \Sigma(\mathring{X})^2 - 2\Sigma(\mathring{X}X) + 2\Sigma(\mathring{X}\mathring{X})$$

18. $$\sum_{i=1}^{n}\sum_{i\neq j=1}^{n}\sum_{i,j\neq k=1}^{n} X_{ii}X_{jk} = \sum_{i=1}^{n}\sum_{i\neq j=1}^{n}\left(\sum_{k=1}^{n} X_{ii}X_{jk} - X_{ii}X_{ji} - X_{ii}X_{jj}\right) =$$
$$= \sum_{i=1}^{n}\sum_{i\neq j=1}^{n}\sum_{k=1}^{n} X_{ii}X_{jk} - \sum_{i=1}^{n}\sum_{i\neq j=1}^{n} X_{ii}X_{ji} - \sum_{i=1}^{n}\sum_{i\neq j=1}^{n} X_{ii}X_{jj} = \quad \text{(by 14,6,5)}$$
$$= \Sigma(X)^2 - \Sigma(\mathring{X})\Sigma(X) - \Sigma(XX) + \Sigma(X\mathring{X}) +$$
$$- \Sigma(\mathring{X}X) + \Sigma(\mathring{X}\mathring{X}) +$$
$$- \Sigma(X)^2 + 2\Sigma(X)\Sigma(\mathring{X}) - \Sigma(\mathring{X})^2 + \Sigma(XX) - 2\Sigma(X\mathring{X}) + \Sigma(\mathring{X}\mathring{X}) =$$
$$= 2\Sigma(\mathring{X}\mathring{X}) + \Sigma(X)\Sigma(\mathring{X}) - 2\Sigma(X\mathring{X}) - \Sigma(\mathring{X})^2$$

**Summations over four indexes**

19. $$\sum_{i=1}^{n}\sum_{j=1}^{n}\sum_{k=0}^{n}\sum_{l=1}^{n} X_{ij}X_{kl} = \left(\sum_{i=1}^{n}\sum_{j=1}^{n} X_{ij}\right)\left(\sum_{k=1}^{n}\sum_{l=1}^{n} X_{kl}\right) = \Sigma(X)^2$$

20. $$\sum_{i=1}^{n}\sum_{i\neq j=1}^{n}\sum_{k=1}^{n}\sum_{l=1}^{n} X_{ij}X_{kl} = \sum_{i=1}^{n}\sum_{k=1}^{n}\sum_{l=1}^{n}\left(\sum_{j=1}^{n} X_{ij}X_{kl} - X_{ii}X_{kl}\right) =$$
$$= \sum_{i=1}^{n}\sum_{j=1}^{n}\sum_{k=1}^{n}\sum_{l=1}^{n} X_{ij}X_{kl} - \sum_{i=1}^{n}\sum_{k=1}^{n}\sum_{l=1}^{n} X_{ii}X_{kl} = \quad \text{(by 19,9)}$$
$$= \Sigma(X)^2 - \Sigma(X)^2 + \Sigma(\mathring{X})\Sigma(X) =$$
$$= \Sigma(X)\Sigma(\mathring{X})$$

21. $$\sum_{i=1}^{n}\sum_{i\neq j=1}^{n}\sum_{i,j\neq k=1}^{n}\sum_{l=1}^{n}X_{ij}X_{kl} = \sum_{i=1}^{n}\sum_{i\neq j=1}^{n}\sum_{l=1}^{n}\left(\sum_{k=1}^{n}X_{ij}X_{kl}-X_{ij}X_{il}-X_{ij}X_{jl}\right)=$$
$$=\sum_{i=1}^{n}\sum_{i\neq j=1}^{n}\sum_{k=1}^{n}\sum_{l=1}^{n}X_{ij}X_{kl}-\sum_{i=1}^{n}\sum_{i\neq j=1}^{n}\sum_{l=1}^{n}X_{ij}X_{il}+$$
$$-\sum_{i=1}^{n}\sum_{i\neq j=1}^{n}\sum_{l=1}^{n}X_{ij}X_{jl}= \text{ (by 20,11,12)}$$
$$=\Sigma(X)\Sigma(\mathring{X})-2\Sigma(X\mathring{X})$$

22. $$\sum_{i=1}^{n}\sum_{i\neq j=1}^{n}\sum_{i,j\neq k=1}^{n}\sum_{i,j,k\neq l=1}^{n}X_{ij}X_{kl} = \sum_{i=1}^{n}\sum_{i\neq j=1}^{n}\sum_{i,j\neq k=1}^{n}\left(\sum_{l=1}^{n}X_{ij}X_{kl}-X_{ij}X_{ki}-X_{ij}X_{kj}-X_{ij}X_{kk}\right)=$$
$$=\sum_{i=1}^{n}\sum_{i\neq j=1}^{n}\sum_{i,j\neq k=1}^{n}\sum_{l=1}^{n}X_{ij}X_{kl}-\sum_{i=1}^{n}\sum_{i\neq j=1}^{n}\sum_{i,j\neq k=1}^{n}X_{ij}X_{ki}+$$
$$-\sum_{i=1}^{n}\sum_{i\neq j=1}^{n}\sum_{i,j\neq k=1}^{n}X_{ij}X_{kj}-\sum_{i=1}^{n}\sum_{i\neq j=1}^{n}\sum_{i,j\neq k=1}^{n}X_{ij}X_{kk}=$$
(by 21,15,16,17)
$$=\Sigma(X)\Sigma(\mathring{X})-2\Sigma(X\mathring{X})-2\Sigma(\mathring{X}\mathring{X})+2\Sigma(\mathring{X}^2)+$$
$$-\Sigma(\mathring{X})\Sigma(X)+\Sigma(\mathring{X})^2+2\Sigma(\mathring{X}X)-2\Sigma(\mathring{X}\mathring{X})$$
$$=\Sigma(\mathring{X})^2-4\Sigma(\mathring{X}\mathring{X})+2\Sigma(\mathring{X}^2)$$
